# Toxin-associated genes cluster beyond gene family size expectations in both venomous and non-venomous animals

**DOI:** 10.64898/2026.10.26.761191

**Authors:** Juan Zhao, Fang Zhao, Feng Zhao

## Abstract

Tandem arrays of toxin genes are widely read as a genomic signature of venom evolution. However, large gene families cluster whatever their function, so the baseline against which toxin clustering should be judged is unclear. We compared the clustering of toxin-associated genes with random genes and with non-toxin families of the same size in 14 independent venom origins, including a new chromosome-level genome and venom-gland proteome of the centipede *Scolopendra mutilans*, and applied the same analysis to toxin homologues of eight non-venomous animals. Against random genes, the odds of a toxin-associated gene having another such gene within 100 kb were 1.8–30 times higher. Against size-matched families, the odds ratio was 2.84 on average (95% CI 1.85–4.36), and its interval excluded 1 in 12 of 14 origins, with high heterogeneity (95% prediction interval 0.97–8.33). The excess consisted mainly of homologous copies lying next to one another. Toxin homologues of non-venomous animals were also clustered beyond the family-size expectation (pooled odds ratio 2.10, 95% CI 1.33–3.32; post hoc), and whether the excess was larger in venomous lineages remained undetermined. When origins were combined, the excess was within a pre-specified equivalence bound of that of secretory and defence families, although no single origin was resolved. Thus, a random-gene baseline greatly overstates how exceptional toxin clustering is, and much of the excess that remains after matching on family size is a property of the gene families concerned rather than of venomous lineages alone.

**Significance:** Venomous animals often carry their toxin genes in clusters, and these clusters are widely taken as evidence that venom genes evolve in a special way. We examined this idea in 14 animal lineages that evolved venom independently and in eight animals without venom. Compared with random genes, toxin-associated genes had up to 30 times the odds of being clustered. Compared with gene families of the same size, the difference shrank to about threefold on average and varied among lineages. Homologues of the same toxin families were also clustered in animals without venom. Much of what looks exceptional about venom-gene clusters therefore depends on the baseline, a point that applies to any gene family said to have an unusual genomic arrangement.

## Introduction

Venoms have evolved independently in many animal lineages, and venomous lineages provide a natural system for studying how new functions are assembled from existing genes (1–3). Toxin genes are often arranged in tandem arrays. Such arrays have been described in snakes, in which venom gene families expanded through repeated tandem duplication (4–6), and in ants (7), and duplicated toxin genes have been reported in spiders (8). These arrays are commonly interpreted as the footprint of repeated local duplication of toxin genes after their recruitment to venom, although this model has also been questioned (9).

However, clustering alone is a weak signature of venom evolution. Most toxins belong to large multigene families, and large families of any function tend to lie in tandem because local duplication is a general mechanism of gene-family growth. Toxins are also mostly secreted proteins, and families of secreted and defence proteins, such as proteases, lectins and antimicrobial peptides, are often expanded in tandem. In the habu genome, for example, non-venom paralogs of the major venom families were duplicated as extensively as the venom genes themselves (10). A toxin set compared with random genes will therefore appear clustered even if toxins behave like other families of the same size or of the same biological role. It remains unclear whether toxin-associated genes cluster more than equally sized non-toxin families, and whether any such excess is specific to venomous lineages.

In this study, we measured the clustering of toxin-associated genes against random genes and against non-toxin families of the same size in 14 independent venom origins, anchored on a new chromosome-level genome and venom-gland proteome of the medicinal centipede *Scolopendra mutilans*, which complements a published assembly of the species (11). We then compared toxins with secretory and defence families of the same genomes and, in a post hoc analysis, applied the same procedure to toxin homologues of eight non-venomous animals. We show that a random-gene baseline overstates the clustering of toxin-associated genes by roughly an order of magnitude in most lineages. Against families of the same size, an excess remained in 12 of 14 origins, averaging 2.84-fold in odds, and toxin homologues of non-venomous animals were also clustered beyond this expectation.

## Results

### Venom origins, representative species and toxin evidence tiers

Candidate venom origins were taken from a published survey of independent venom evolution (table S1 in ref. 1), and each origin was represented by one species. A species was eligible if it had an annotated genome assembled to chromosome scale or with a scaffold N50 of at least 0.5 Mb, BUSCO completeness of at least 80% and ToxProt homologues from which a toxin set could be built. Of 15 candidate origins, 14 met these criteria; the pseudoscorpion candidate was excluded because its proteome was 40.5% complete (SI Appendix, Table S1 and Fig. S7). Where several species of an origin were eligible, the representative was chosen by a fixed order: a published venom proteome that could be mapped to the assembly, chromosome-scale assembly, BUSCO completeness and contig N50.

The centipede representative is a new chromosome-level assembly of *S. mutilans* (1.49 Gb; 14 chromosome-scale scaffolds spanning 949.5 Mb; contig N50 50.0 Mb; Merqury QV 68.8; BUSCO 98.6%; SI Appendix, Note 2). Its toxin catalogue contains 188 genes that match UniProt ToxProt and are enriched in venom-gland transcripts. A data-independent-acquisition proteome of venom glands from 12 adults detected peptides for 163 of the 188 toxins and resolved 141 of them to single loci by genome-unique peptides (SI Appendix, Table S17).

Toxin identity was graded into three tiers defined independently of clustering. The strict tier (P) required at least one peptide unique to the gene in a published venom proteome or in our venom-gland proteome. The intermediate tier added venom-gland expression (E), and the broad tier added ToxProt homology alone (H). The broad tier is the only one available for all species and was used in all cross-origin analyses. The strict tier was estimable in the centipede (143 genes in 26 families with at least two toxins) but not in any of the 13 other species, in which at most eight such families were available (Fig. 1B). We therefore refer to the genes analysed across origins as toxin-associated genes.

**Fig. 1.**
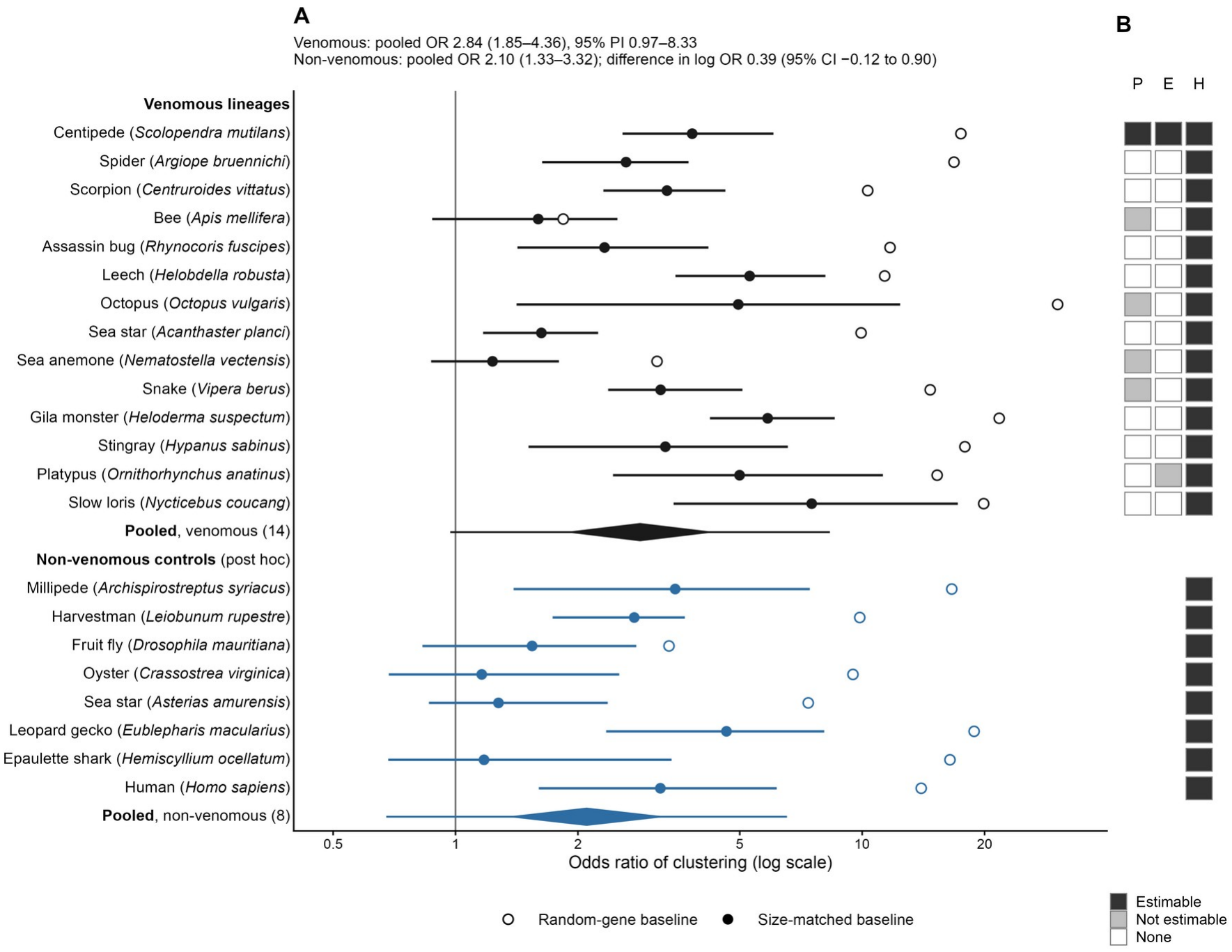
Clustering of toxin-associated genes against random and family-size-matched baselines in 14 venom origins and eight non-venomous animals. (*A*) Odds ratio for a toxin-associated gene having another such gene within 100 kb, relative to random protein-coding genes (open circles; point estimates) and to genes drawn from non-toxin families of the nearest size (filled circles; bars, 95% family-block bootstrap intervals from 2,000 resamples of toxin families). Each baseline used 1,000 draws (broad tier, primary family definition). Black, venomous lineages (one representative species per origin); blue, ToxProt homologues of non-venomous animals (post hoc). The black diamond shows the pooled estimate of the phylogenetic multilevel model (2.84, 95% CI 1.85–4.36; I^2^ = 82.7%), and the line through it the 95% prediction interval (0.97–8.33). The blue diamond shows the random-effects estimate for the non-venomous animals (2.10, 1.33–3.32; prediction interval 0.68–6.53). The difference in log odds ratio between the two groups (0.39, 95% CI −0.12 to 0.90) is from a meta-regression with the Hartung–Knapp adjustment. (*B*) Toxin evidence tiers: P, gene-unique venom peptide; E, venom-gland expression; H, ToxProt homology. Dark, estimable (at least ten gene families containing two or more genes of the tier); grey, genes present but not estimable; white, no genes. Assemblies and numbers of genes per tier are given in SI Appendix, Tables S1 and S26.

### Toxin-associated genes cluster beyond family-size expectations in most venom origins

Clustering was measured as the fraction of toxin-associated genes that have another such gene within 100 kb, and each toxin set was compared with two baselines (Materials and Methods). The first draws random protein-coding genes. The second draws, for every toxin family, a non-toxin family of the nearest available size and samples as many genes from it as the toxin family contributes.

Against random genes, the odds of a toxin-associated gene being clustered were 1.84 (honey bee) to 30.25 (octopus) times higher across the 14 origins (Fig. 1A; SI Appendix, Table S2). However, against size-matched families, the odds ratio fell to 1.23–7.52, and its family-block interval excluded 1 in 12 origins. The two exceptions were the sea anemone (1.23, 95% CI 0.87–1.80) and the honey bee (1.60, 0.88–2.50). The 14 log odds ratios were combined in a phylogenetic multilevel model, with between-origin correlations derived from TimeTree divergence times (12). The average odds ratio was 2.84 (95% CI 1.85–4.36). Heterogeneity was high (I^2^ = 82.7%), and the 95% prediction interval was 0.97–8.33, so an excess was present on average but its size differed among origins. Omitting each origin in turn gave odds ratios of 2.66–3.20, and every interval excluded 1.

These estimates were robust to the analytical choices that we could vary. For the spider–scorpion divergence, which is absent from TimeTree, the median of four published estimates (13–16) was used (455.5 Mya); the youngest and oldest estimates gave 3.11 (2.28–4.25) and 2.84 (1.86–4.35). Counting spiders and scorpions as one origin gave 2.85 (1.87–4.35). Removing genes within 10 kb of an assembly gap or overlapping other gene models left 13 estimable origins and an average of 2.98 (1.98–4.48). Restricting the panel to the six origins with proteome or venom-gland expression evidence gave 2.70 (1.46–5.02; post hoc), and in the centipede the strict tier gave 4.43 (3.03–6.75; post hoc), compared with 3.82 (2.57–6.05) for the full catalogue.

For three lineages, a second species had been analysed with the same procedure, and both species of each pair gave intervals excluding 1: *Crotalus adamanteus* 6.16 (3.94–10.42) and *Vipera berus* 3.19 (2.37–5.08), *Stegodyphus dumicola* 3.67 (1.89–6.54) and *Argiope bruennichi* 2.63 (1.63–3.74), and *Centruroides sculpturatus* 2.79 (2.12–3.92) and *C. vittatus* 3.31 (2.31–4.61). In addition, in these five-genome analyses, the excess was also found under two other gene-family definitions, for windows of 25, 50 and 200 kb, after additional matching on within-family protein identity, and with the venom genes annotated by the authors of the snake genome (17) (SI Appendix, Fig. S8 and Table S24).

### Toxin homologues of non-venomous animals are also clustered

To ask whether the excess is specific to venomous lineages, we applied the same procedure, in a post hoc analysis registered before any calculation, to ToxProt homologues of eight non-venomous animals chosen by fixed rules from the groups represented in the venom panel (Materials and Methods). Against random genes, these homologues had odds ratios of 3.35–18.85. Against size-matched families, the odds ratio was 1.16–4.64, and its interval excluded 1 in four species: the millipede *Archispirostreptus syriacus* (3.47, 1.39–7.43), the leopard gecko *Eublepharis macularius* (4.64, 2.35–8.07), human (3.19, 1.60–6.16) and the harvestman *Leiobunum rupestre* (2.75, 1.73–3.66) (Fig. 1A; SI Appendix, Table S26). Thus, the pooled odds ratio of the non-venomous animals was 2.10 (1.33–3.32; prediction interval 0.68–6.53). The difference in log odds ratio between venomous and non-venomous animals was 0.39 (95% CI −0.12 to 0.90), which under the pre-specified rule was classified as undetermined; restricting the venomous side to the 13 species analysed with public genomes, or to their homology tier, gave the same classification.

### Comparison with secreted and chemoreceptor gene families

Toxins are mostly secreted proteins, and secreted or defence-related families may themselves lie in tandem arrays. Each genome was therefore searched for two classes of non-toxin control families with the same Pfam and signal-peptide rules in all species (Materials and Methods). The secretory and defence class (S) comprised families dominated by antimicrobial peptides, lysozymes, C-type lectins, trypsin-like serine proteases, metalloproteases and secreted lipases. The lineage-expansion class (L) comprised chemoreceptor families, which were defined only for arthropods and vertebrates. Families containing any toxin-associated gene were removed from the control classes. For each class, the size-matched odds ratio was calculated with the same null model as for toxins, and the difference Δ = log OR_toxin − log OR_class was compared with a pre-specified equivalence bound of ±0.5.

However, the S class was estimable in 11 origins, and Δ was uncertain in each of them (Fig. 2). When the 11 differences were combined, the pooled Δ was 0.15 (90% CI −0.12 to 0.41; 95% CI −0.18 to 0.47). The 90% interval lay within the equivalence bound: averaged over origins, toxin-associated genes were clustered to a similar degree as secretory and defence families. The rule for combining origins was added after the five-genome comparison had been seen but before any control class of the added species was computed (SI Appendix, Note 1, D9). The phylogenetic variance component was estimated close to zero, so the pre-specified random-effects model with the Hartung–Knapp adjustment was used; the multilevel model gave the same classification. Counting only class members that carried the defining domain gave a pooled Δ of −0.01 (90% CI −0.28 to 0.25). The 95% prediction interval of the pooled difference (−0.41 to 0.70) extended beyond the bound. In a post hoc analysis restricted to the 143 proteome-supported centipede toxins, Δ was 0.69 (90% CI −0.03 to 1.57) for all class members and 0.77 (95% CI 0.03–1.93) for domain-bearing members (SI Appendix, Note 1, D11). The L class was estimable in only three origins. In the honey bee and the assassin bug, toxins were less clustered than chemoreceptors (Δ −2.17, 95% CI −3.66 to −0.97; and −1.59, −2.62 to −0.24). Because the plan defined only “more clustered” and “similar”, both comparisons are recorded as undetermined; we report the direction here.

**Fig. 2.**
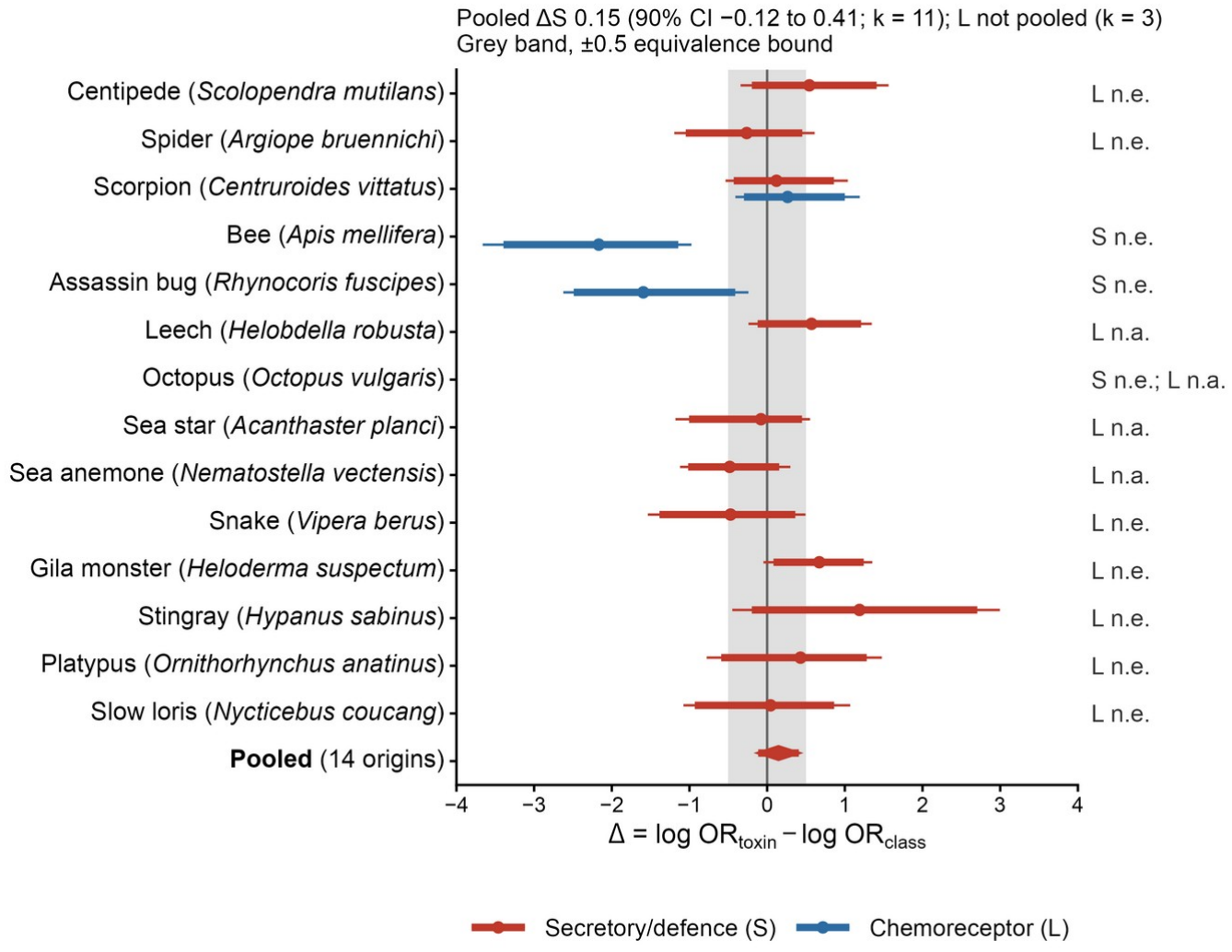
Comparison of toxin-associated genes with secretory and defence families and with chemoreceptor families of the same genomes. Δ, difference in log odds ratio between toxin-associated genes and secretory and defence families (S, red) or chemoreceptor families (L, blue); thick bars, 90% intervals; thin bars, 95% intervals (2,000 independent family-block resamples). The grey band marks the pre-specified equivalence bound (±0.5). n.e., not estimable (fewer than ten qualifying control families or an undefined control odds ratio); n.a., class not defined for the phylum. The red diamond shows the pooled S-class difference across 11 estimable origins (0.15, 90% CI −0.12 to 0.41; random-effects model with the Hartung–Knapp adjustment). The L class was estimable in three origins and was not pooled.

### The excess consists mainly of adjacent homologous copies

The composition of the excess was examined in the five genomes for which family-level decompositions were available (*S. mutilans, C. adamanteus, S. dumicola, C. sculpturatus* and *Nematostella vectensis*). Because the excess is the difference between observed and expected clustered genes, it can be divided exactly among toxin families (Materials and Methods). The largest single family carried 7% (centipede), 13% (snake), 32% (spider), 13% (scorpion) and 26% (sea anemone) of the positive excess, and 15, 5, 4 and 9 families were required to account for half of it in the four bilaterians (Fig. 3A). None of these shares was clearly larger than expected from the same decomposition applied to control draws (largest-family share, *P* = 0.088 in the spider and ≥ 0.43 elsewhere). When each of the five largest contributors was removed in turn, the excess remained in every bilaterian species (odds ratios 2.62–6.12).

**Fig. 3.**
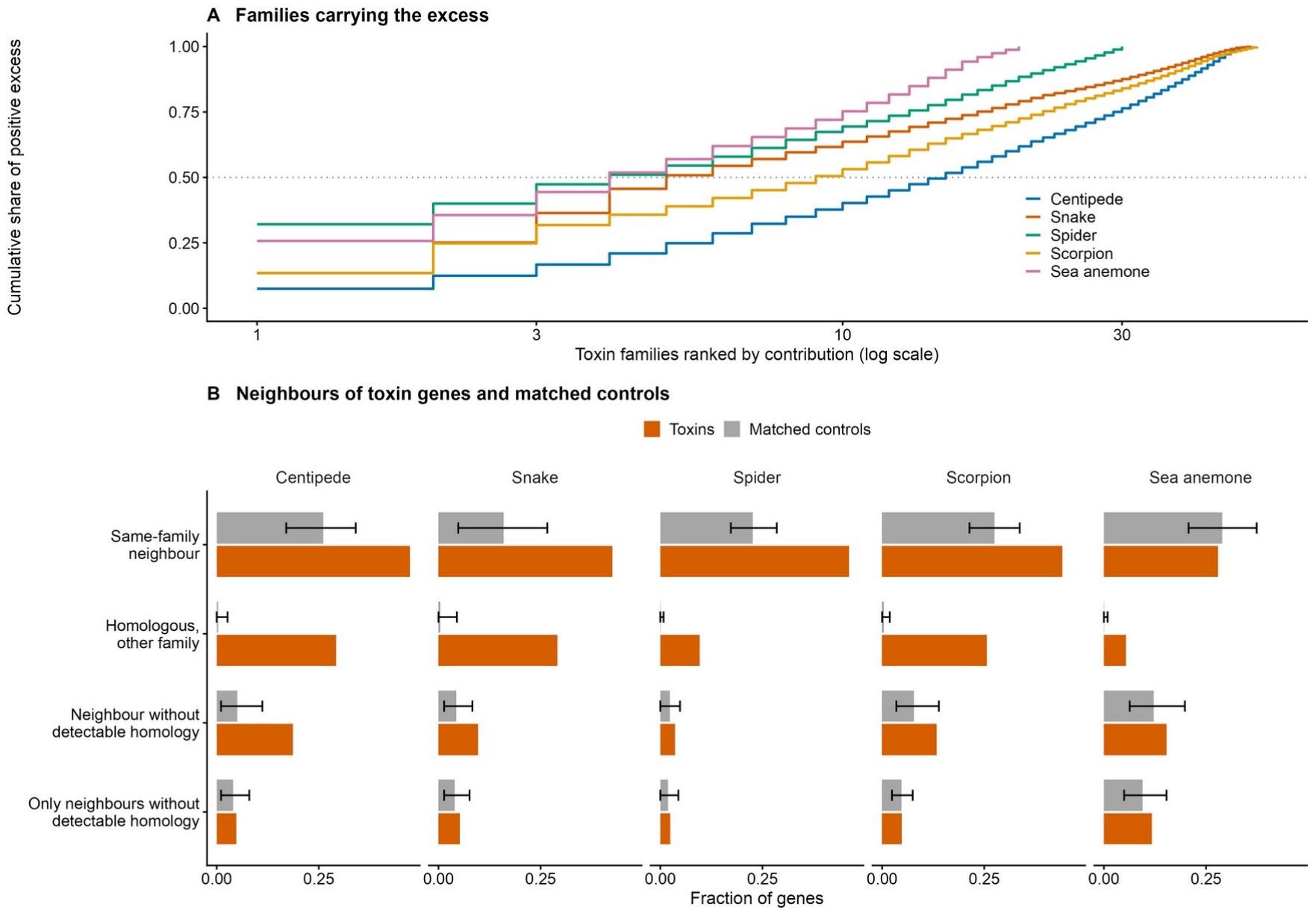
Distribution and composition of the excess in five venomous genomes. (*A*) Cumulative share of the positive excess carried by toxin families ranked by their contribution (primary definition). (*B*) Fraction of toxin genes (orange) and of genes sampled from size-matched non-toxin families (grey; mean and 2.5–97.5% range over 200 draws) that have, within 100 kb, a neighbour from their own family, a homologous neighbour from another family, a neighbour without detectable homology (DIAMOND *E* < 10^−3^), or only neighbours without detectable homology. The first three categories are not mutually exclusive, and the last is a subset of the third. Denominators are all toxin genes and all sampled control genes.

We next examined what kind of neighbour makes a toxin gene count as clustered. Toxins had a neighbour from their own family more often than matched controls (centipede 0.47 versus 0.26; snake 0.43 versus 0.16; spider 0.46 versus 0.23; scorpion 0.44 versus 0.28), and they also had more neighbours assigned to other families (Fig. 3B). When every pair of neighbouring genes from different families was searched for detectable homology (DIAMOND, *E* < 10^−3^), a homologous neighbour from another family was found for 0.29 of centipede toxins (≤ 0.005 of controls), 0.29 of snake toxins, 0.26 of scorpion toxins and 0.10 of spider toxins. Neighbours without detectable homology were modestly in excess in the centipede (0.19 versus 0.05), snake (0.10 versus 0.04) and scorpion (0.13 versus 0.08), but not in the spider or sea anemone. The excess is therefore carried mainly by homologous toxin copies lying next to one another.

### Duplication topology, expression and conservation of toxin and non-toxin arrays

Further tests addressed how toxin arrays might differ from other tandem families, and none separated toxin from control families (SI Appendix, Fig. S9). Strongly supported sister copies lay closer together than each family’s own layout predicts in both toxin families (median Δ*f* 0.23–0.60) and size-matched non-toxin families (0.31–0.49), with toxin-minus-control differences of +0.14 to +0.23 (Holm-adjusted *P* ≥ 0.41). The per-copy expression of tandem relative to non-tandem copies did not differ detectably between toxin and control families, in RNA or in venom-gland protein (Holm-adjusted *P* ≥ 0.067). *S. mutilans* toxin arrays were located in the congener *Scolopendra cretica* less often than non-toxin arrays (34 of 90 versus 425 of 893; family-level difference −0.17, 95% CI −0.30 to −0.03; Holm-adjusted *P* = 0.033), whereas copy-number discordance was uncertain. We also attempted to order array formation and venom recruitment by comparison with non-venomous outgroups; the order could be resolved for only 4 of 63 families, all in the snake, and the pre-specified stopping rule applied (SI Appendix, Note 6 and Table S4).

## Discussion

Toxin genes cluster far more than random genes, and this is often taken as a distinctive feature of venom genomes. In this study, the odds ratio against random genes was 1.8–30 across 14 venom origins and fell to 1.2–7.5 once non-toxin families of the same size were used as controls, averaging 2.84. This reduction reflects the choice of baseline and should not be read as a proportion of clustering attributable to family size. The size of the remaining excess differed among origins, and the prediction interval reached 1.

The non-venomous controls show that much of this remaining excess is not specific to venomous lineages. When the same procedure was applied to ToxProt homologues of eight animals without venom, these genes were also clustered beyond the family-size expectation, by a pooled odds ratio of 2.10, and the millipede, leopard gecko, human and harvestman each gave intervals excluding 1. However, the venomous lineages had a higher average, and the difference was not resolved. This result agrees with the habu genome, in which non-venom paralogs of venom families were duplicated as extensively as the venom genes (10), and with evidence that oral venom systems are built on secretory gene networks shared with other glands (18). In bees, most venom genes originated by co-option of single genes, and duplication contributed mainly to their later diversification (19). Thus, toxin families appear to be drawn from gene families that already tend to form local homologous arrays, and whether recruitment to venom adds to this tendency remains open. The order of recruitment and array formation could be resolved for only 4 of 63 families in our data.

The comparison with secretory and defence families points in the same direction. When the 11 estimable origins were combined, the difference between toxin-associated genes and these families lay within the pre-specified equivalence bound, but its prediction interval (−0.41 to 0.70) extended beyond the bound, and no single origin was resolved. In the 13 added species, toxin-associated genes were defined mainly by ToxProt homology, and families containing any such gene were removed from the control class. Secreted homologues that are not venom components may therefore have been counted as toxins, which would draw the two sets together. In agreement with this possibility, the proteome-supported toxins of the centipede were more clustered than its domain-bearing secretory and defence genes (Δ 0.77, 95% CI 0.03–1.93; post hoc), although the difference relative to all members of these families was uncertain.

The composition of the excess was similar across species. The excess was spread across many toxin families, and no concentration in a few families was clearly beyond that seen in control draws. It consisted mainly of homologous toxin copies lying next to one another. In the centipede, the largest toxin cluster is a 271-kb array of 15 scoloptoxin-like genes, of which 13 are resolved by genome-unique venom-gland peptides. In the snake, the three families contributing most to the excess are a myotoxin, a snake venom metalloproteinase and a serine-protease family, consistent with the tandem venom-gene arrays reported for this species (17). In addition, removing any of the five largest contributors left the excess in place.

Three analyses designed to discriminate how such arrays arise or persist did not separate toxin from control families. Sister copies lay close together in both, the expression of tandem copies did not differ detectably, and toxin arrays were located in a congener less often than other arrays. The excess is therefore compatible with toxin families forming local homologous arrays at a higher rate, retaining them for longer, or both, and differences in family age also remain possible.

Three limitations apply to these conclusions. Each origin and each control is represented by one species. The strict, proteome-supported tier was estimable only in the centipede, so across species the analysis concerns toxin-associated genes defined largely by homology, as it does in the non-venomous controls. The pooled models used standard errors derived from bootstrap intervals that are not always symmetric, and the non-venomous controls, as well as the present title, were added after the main results had been seen.

In summary, toxin-associated genes were clustered beyond the family-size expectation in most of 14 independent venom origins, by about threefold in odds on average, and homologues of the same families were also clustered in non-venomous animals. Claims of exceptional toxin-gene architecture therefore depend on comparison with families of the same size, with families of a similar secreted role and with the same families in animals without venom.

## Materials and Methods

### Study design and analysis plan

The analysis plan was frozen on 10 November 2026 before any new end point was computed. It was kept as a dated internal document and was not deposited in a public registry. It separated results already reported for five genomes in a preprint version of this study, post hoc analyses of those genomes and new prospective tests (evidence tiers in added species, control families, the origin panel and the order of recruitment). It specified inclusion rules, evidence tiers, family definitions, statistics, equivalence bounds, interpretation rules and stopping rules. Clarifications and corrections were appended to a dated deviation log before the affected analyses were run; the log is reproduced in SI Appendix, Note 1. One correction concerns the centipede: analyses run before 13 November 2026 had used an earlier assembly of the same reads, and all centipede results reported here were recomputed on the assembly described below. The non-venomous controls and the present title were added after all other results had been seen (SI Appendix, Note 1, D13 and D14).

### Centipede genome, toxin catalogue and venom-gland proteome

Adult *S. mutilans* were collected in Luoshan County, Xinyang, Henan Province, China, in 2023. All animal procedures were reviewed and approved by the Laboratory Animal Welfare and Ethics Committee of Pu’er University (Approval No. PEU-AEC-2023-119). Sampling, sequencing, assembly, annotation and proteomics are described in SI Appendix, Note 2. Briefly, the *S. mutilans* genome was assembled from PacBio HiFi and Hi-C reads with hifiasm (20, 21) in Hi-C phasing mode and scaffolded with YaHS (22). Predicted proteins were searched against UniProt ToxProt (23) with DIAMOND v2.2.2 (24) (*E* < 10^−5^), and hits were required to show venom-gland-enriched expression (venom gland/control ≥ 2; mean ≥ 5 TPM). Raw reads were available for only 3 of the 18 RNA-seq libraries, so expression values were transferred from an earlier annotation of the same reads (SI Appendix, Note 2). Six physiological proteins that matched ToxProt only through a shared domain were removed, leaving 188 toxins. A data-independent-acquisition proteome of dissected venom glands was obtained from 12 adults, and a toxin was resolved to a single locus when it carried a peptide found in no other predicted protein.

### Venom origins, representative species and evidence tiers

Candidate origins were taken from the table of independent venom origins in ref. 1. An origin was eligible if a species had an NCBI or Ensembl assembly at chromosome scale or with a scaffold N50 ≥ 0.5 Mb, a protein-coding annotation, BUSCO v6 (25) protein completeness ≥ 80% with the lineage-specific odb12 dataset and ToxProt homologues. One representative per origin was chosen in the registered order given in Results, and assemblies are listed in SI Appendix, Table S1. For each protein-coding gene, the longest isoform was kept.

Toxin-associated genes were assigned to three tiers without using clustering information. The broad tier (H) comprised genes whose proteins had a DIAMOND hit to ToxProt (*E* < 10^−5^, query coverage ≥ 50%, alignment ≥ 40 amino acids). The intermediate tier added genes listed in the venom-gland transcriptome toxin tables of the original studies or meeting the centipede expression criterion (E). The strict tier (P) required at least one gene-unique peptide in a published venom proteome or in our venom-gland proteome; published proteins were mapped through accessions or, when none were given, by ≥ 95% identity over ≥ 90% of both sequences. A tier was estimable when at least ten families contained two or more of its genes and at least 95% of bootstrap resamples were valid. For the four comparator genomes of the five-genome analysis, the broad tier was the primary input.

### Gene families and size-matched clustering

The primary gene-family definition was MMseqs2 (26) (release 18-8cc5c) clustering of canonical proteins at minimum identity 0.3 with bidirectional coverage 0.5 and cluster reassignment. For the five genomes, two further definitions were used as sensitivity analyses (SI Appendix, Note 3). The analysis plan had named the unidirectional-coverage definition as primary; the canonical-isoform definition is used as primary here, and both gave intervals excluding 1 in the four bilaterians (SI Appendix, Note 1 and Fig. S8). A gene was called clustered if another member of the same set lay within 100 kb on the same sequence. The statistic was the fraction of toxin-associated genes clustered. The random-gene baseline drew equal numbers of random protein-coding genes (1,000 draws). In each of 1,000 draws, the size-matched baseline chose for every toxin family a non-toxin family of the nearest available size that contained at least as many genes as the toxin family contributed, and sampled that number of genes from it. Clustering was then evaluated over the union of the sampled genes. Effects are odds ratios of the observed fraction against the mean control fraction. Intervals were obtained by resampling toxin families with replacement (2,000 resamples), carrying with each family its observed and expected clustered counts; spatially connected groups of families were also resampled as a check. The random seed was 20260724.

The excess of each toxin family was its observed number of clustered toxins minus the mean number expected from its matched controls, and these values sum to the total excess. Neighbours from different families were tested for homology with DIAMOND (*E* < 10^−3^) in toxins and controls alike. Duplication topology, expression and congener analyses followed the registered designs and are described in SI Appendix, Note 4.

### Control families

Control classes were defined with the same rules in all species. Proteins were annotated with hmmscan (HMMER v3.4) (27) against Pfam-A (28), keeping domains with independent *E* < 10^−5^, and signal peptides were predicted with DeepSig v1.2.5 (29) in eukaryote mode. The secretory and defence class (S) comprised families in which at least half of the members carried one of the registered domains (antimicrobial peptides and defensins, lysozymes, C-type lectins, trypsin-like serine proteases, metalloproteases and secreted lipases) and at least half carried a signal peptide. The lineage-expansion class (L) comprised chemoreceptor families, defined by 7tm_6, 7tm_7 and Lig_chan domains in arthropods and by 7tm_4 domains in vertebrates; it was not defined for other phyla. Qualifying families had at least two members and no toxin-associated gene, and families containing toxins were counted and excluded. All members of qualifying families formed the primary class set, and members carrying the defining domain were used in a sensitivity analysis. The odds ratio of each class was computed with the same size-matched null model, with toxin families and class families excluded from the control pool. The difference Δ = log OR_toxin − log OR_class was estimated with independent family-block resampling of toxin and class families (2,000 resamples). A class was not estimable with fewer than ten qualifying families. Toxins were classified as more clustered when the lower 95% limit of Δ exceeded 0, and as similar when the 90% interval lay within ±0.5; otherwise the comparison was classified as undetermined.

### Combination across origins

Effect sizes were log odds ratios, with standard errors taken as the width of the 95% family-block interval on the log scale divided by 3.92. Between-origin correlations followed a Brownian model on the time tree, R_ij = 1 − t_ij/T, in which t_ij is the median pairwise divergence time from the TimeTree public interface (12) and T = 721.45 Mya is the largest pairwise time. TimeTree returned no time for spiders and scorpions, so the median of four published fossil-calibrated estimates of their most recent common ancestor (13–16) was used (455.5 Mya), with the minimum and maximum as sensitivity analyses (SI Appendix, Note 5). A multilevel model with phylogenetic and origin random effects was fitted by restricted maximum likelihood with metafor 5.0.1 (30) in R. If either variance component was zero, smaller than 10^−4^ of the other or failed to converge, a random-effects model with the Hartung–Knapp adjustment (31) was used instead. We report the pooled estimate, its 95% CI and prediction interval, I^2^ and leave-one-origin-out estimates. Pooled estimates of Δ for the control classes used the same models and were computed only when at least ten origins were estimable. Spiders and scorpions were also counted as a single origin by giving them a shared origin effect.

A local quality check was applied to all species as a sensitivity analysis. Genes within 10 kb of an assembly gap (runs of at least ten N) or overlapping another gene model were removed from toxins and controls alike, and toxin genes flagged as possible split or merged models from their ToxProt alignments were also removed.

### Non-venomous controls (post hoc)

The rules for choosing controls, the end point and its interpretation were registered before any control genome was downloaded (SI Appendix, Note 1, D13). For each group of the venom panel, one animal outside all venomous lineages of ref. 1 was chosen by the eligibility criteria and the order of choice used for the representative species, preferring the same class and then the same phylum; the two non-venomous outgroups chosen earlier for the analysis of recruitment were used for myriapods and squamates, and human was used for mammals. No annelid met the criteria, and cnidarians have no non-venomous lineage. ToxProt homologues of the eight controls were identified and analysed with exactly the same procedure and parameters as the broad tier of the venom panel. The difference in log odds ratio between venomous origins and controls was estimated by meta-regression (random-effects model, restricted maximum likelihood, Hartung–Knapp adjustment) and classified with the same rule and bound (±0.5) as the control-class comparison.

### Order of recruitment

For pre-specified toxin families with published venom-peptide evidence, orthologous loci were sought in non-venomous outgroups (millipedes for the centipede; non-toxicoferan squamates for the snake). Gene trees, two-sided syntenic anchors and gene-conversion tests were used to classify each family as an ancestral array, an expansion after recruitment or not orderable (SI Appendix, Note 6). The analysis was to be reported only if at least half of the families, and at least six, could be ordered with a consistent direction.

### Use of generative AI

Generative AI tools (Claude Opus 5.5 and Claude Fable 5.2; Anthropic) were used to assist with writing and running analysis code, checking the consistency of numbers between result files and the text, and editing the English text. Figures were drawn by the analysis scripts from the result tables; no image was generated or edited with generative AI. The authors checked all code, results and text and take full responsibility for the content of this article.

## Supporting information

Supplementary Information (Notes 1-7, Figs. S1-S9, legends for Tables S1-S26 and Datasets S1-S4, SI references)

Datasets S1-S4 (analysis plan and deviation log with English translations; toxin-associated gene tiers; literature screen)

## Data, Materials, and Software Availability

This Whole Genome Shotgun project has been deposited at DDBJ/ENA/GenBank under the accession JCANLD000000000 (BioProject PRJNA1462551; BioSample SAMN61429659) (32). Version JCANLD010000000 is the earlier provider assembly (SI Appendix, Note 2), and the assembly and annotation described in this paper (Smutilans v3.1) are being deposited as an update of this project. The venom-gland DIA proteome (12 raw files and the DIA-NN (33) search report) is deposited in iProX under project accession IPX0020582000 and in ProteomeXchange under PXD086253 (34) (private until publication). The search database comprises the 16,703 primary predicted proteins analysed in this paper, 12 additional toxin isoforms and 116 cRAP contaminant entries (16,831 sequences). The four re-analysed shotgun datasets (proteome and peptidome of the venom gland and of the torso) are available at iProX (IPX0001117000) (35, 36); only the venom-gland proteome was analysed in the original publication. Raw PacBio HiFi, Iso-Seq, Hi-C, short-read WGS and small-RNA reads have been deposited in the SRA under BioProject PRJNA1462551 (19 BioSamples; 22 runs, SRR39514747–SRR39514768; private until publication). Raw reads of the three venom-gland RNA-seq libraries that remain available are being deposited under the same BioProject; raw reads of the other 15 libraries are no longer available. The assembly and annotation analysed in this paper (FASTA, GFF3, protein and CDS sequences) and the expression matrix of the 18 RNA-seq samples are deposited at Zenodo (https://doi.org/10.5281/zenodo.23975847; files restricted until publication) (37). Public assemblies are listed in SI Appendix, Table S1 (representative species), Note 3 (comparator genomes of the five-genome analysis), Note 6 (outgroups) and Table S26 (non-venomous controls); all public assemblies are cited in SI Appendix, SI References, and the per-species toxin sets in Dataset S3.

Analysis scripts for the cross-species comparison are available at https://github.com/zhaofeng-zyh/scolopendra-venom-genomics and archived at Zenodo (https://doi.org/10.5281/zenodo.21212745) under the MIT License. Scripts for the analyses added in this paper will be released at publication and are available from the corresponding author on request until then.

## Acknowledgments

Genome and transcriptome sequencing were performed by Shanghai Biozeron Biotechnology Co., Ltd. (Shanghai, China).

## Funding

This work was supported by the National Natural Science Foundation of China (82260851 and 81860696), the Joint Special Project for Basic Research of Local Universities in Yunnan Province (202101BA070001-143) and the Key Research and Development Program of Yunnan Province (202403AP140041).

## Author contributions

Feng Zhao and Juan Zhao designed research; Juan Zhao and Fang Zhao performed research; Feng Zhao and Juan Zhao analyzed data; and Feng Zhao wrote the paper. All authors reviewed and approved the final version.

## Competing interests

The authors declare no competing interests.

## Notes

### Competing Interest Statement

The authors have declared no competing interest.

### Summary of Updates

This version corresponds to the manuscript submitted to PNAS. The analysis was extended from five genomes to 14 independent venom origins, one representative species each, with three toxin evidence tiers, secretory/defence and chemoreceptor control families, and a phylogenetic multilevel model across origins. In a post hoc analysis, the same procedure was applied to ToxProt homologues of eight non-venomous animals, which were also clustered beyond the family-size expectation. The title, abstract and text were rewritten accordingly. The five-genome analyses of the previous version are retained, and the dated analysis plan and deviation log are provided as Datasets S1 and S2.

