## Supplementary Information (Notes 1-7, Figs. S1-S9, legends for Tables S1-S26 and Datasets S1-S4, SI references) for "Toxin-associated genes cluster beyond gene family size expectations in both venomous and non-venomous animals"

#### This PDF file includes

Notes 1 to 7

Figs. S1 to S9

Legends for Tables S1 to S26 (tables are provided in SI_Tables_S1-S26.xlsx)

Legends for Datasets S1 to S4

Public genome assemblies analysed

SI References

#### Note 1. Analysis plan, deviations and corrections

##### Registration

The analysis plan was written in Chinese and frozen on 10 November 2026, after an internal panel review and before any new end point was computed. The frozen plan and its dated deviation log are provided as Datasets S1 and S2, in the Chinese original with an English translation (SHA-256 of the Chinese originals: Dataset S1 a7a1d1a9e813f74e37ad96ebba5820f411441a93ecc8e3a2c52dfd17e32c89dc; Dataset S2 3b29f2643c97ae7b865d2dd0004a4b5757762f4860326abf2fa33fbd054bae7f). The plan separated three kinds of analysis. Existing results (E) are those reported in our earlier five-genome analysis, which had already been seen. Post hoc analyses (P) are new analyses of those five genomes. Prospective analyses (N) are end points that had never been computed: evidence tiers in added species, control families in all genomes, the origin panel and the order of recruitment. Only N analyses are treated as tests specified in advance.

The plan fixed the following elements before computation: inclusion rules for venom origins and representative species; the three toxin evidence tiers and the estimability threshold (ten families containing at least two toxin-associated genes, with at least 95% valid bootstrap resamples); the composition of the two control classes and their Pfam domains; the gene-family definition and statistic; the equivalence bound for control comparisons (±0.5 on the natural-log odds ratio scale); the stopping rules (the origin panel was to stop with fewer than six eligible origins, and the order of recruitment with fewer than half, or fewer than six, orderable families); the primary model for pooling across origins; and the wording to be used for each pattern of results. For the origin panel, pooled estimates were to be treated as sensitivity analyses with fewer than ten origins, and as the primary pooled analysis with ten or more.

#### Deviations and corrections

Each entry below was appended to the log before the affected analysis was run, unless stated otherwise. Entries D1–D9 are clarifications of points the frozen plan did not specify; D10 is a correction of an input error; D11 is a post hoc analysis made in response to an internal review; D12 is a deviation in the choice of title; D13 is a post hoc analysis of non-venomous controls whose rules were registered before computation; and D14 replaces the D12 title after D13.

| Entry | Date (2026) | Subject | Content | Timing |
|---|---|---|---|---|
| D1 | 10 Nov | Origin list | A single reference table of independent venom origins (table S1 in ref. 1) defined the candidate origins (15); snakes and helodermatid lizards, and spiders, scorpions and pseudoscorpions, counted | Before eligibility screening |

| Entry | Date (2026) | Subject | Content | Timing |
| --- | --- | --- | --- | --- |
| | | | separately in the primary count and merged only in sensitivity analyses. | |
| D2 | 11 Nov | Control-class membership | Membership of the control-class set was specified (domain-bearing members, signal peptide for S), together with the Pfam evalue column, the not-applicable L class in cnidarians and the bootstrap settings for Δ. The primary set was later changed to all members of qualifying families, with domain-bearing members as a sensitivity analysis. | Specification before computation; the change of primary set was decided after the five-genome results had been seen, and both sets are reported |
| D3 | 11 Nov | First rank of representative choice | A "published venom proteome that can be mapped" required a peer-reviewed mass-spectrometry study of venom listing identified proteins, of which ≥50% mapped to the selected annotation. Two points were clarified after mapping results were known: one qualifying paper suffices, and only identifications | Rule before mapping; two clarifications after, with results under both readings retained |

| Entry | Date (2026) | Subject | Content | Timing |
|---|---|---|---|---|
| | | | classified as venom components by the original authors count. | |
| D4 | 11–12 Nov | Strict and intermediate tiers in added species | P counted only identifications classified as venom components, using gene-unique peptides of ≥7 residues (I/L equivalent) or, without peptides, protein identity ≥95% over ≥90% of both sequences; E used only venom-gland toxin tables from the original studies. | Before any clustering end point in added species |
| D5 | 12 Nov | Order of recruitment | Outgroups (millipedes for the centipede; *Eublepharis macularius* and *Podarcis raffonei* for the snake), source arrays, syntenic anchoring, copy counting, gene-tree criteria and gene-conversion screening (GENECONV 1.81a) were specified; families with too few sequences or too few polymorphisms were classified as not orderable. | Before any classification |
| D6 | 12 Nov | Pooling across origins | TimeTree divergence times, the correlation matrix, standard errors from | Before the pooled model was fitted |

| Entry | Date (2026) | Subject | Content | Timing |
|---|---|---|---|---|
| | | | bootstrap intervals, the multilevel model and its fallback were specified. TimeTree returned no time for spiders and scorpions; alternative species pairs were registered and also failed, and the median of published estimates was then registered (screening criteria in Note 5). | |
| D7 | 13 Nov | Spiders and scorpions as one origin | Implemented as a shared origin effect for the two species, with the phylogenetic effect unchanged. | Before computation |
| D8 | 13 Nov | Local quality check | Gap definition (≥10 consecutive N within 10 kb), gene-model overlap, split- and merged-model criteria from ToxProt alignments, and recomputation through filtered coordinates. | Before any flag was produced |
| D9 | 13 Nov | Control classes in the origin panel | Extension of control classes to all representatives; the panel-level verdict pooled Δ across estimable origins only when at least ten were estimable; the title wording | Before any annotation of added species was used |

| Entry | Date (2026) | Subject | Content | Timing |
|---|---|---|---|---|
| | | | referring to secreted families was tied to the panel-level verdict for the S class. | |
| D10 | 13 Nov | Correction of centipede input | Analyses of control classes and of the origin panel run before 13 November had used an earlier assembly of the same centipede reads. All centipede results, including the order of recruitment, were recomputed on the assembly reported here; earlier results were archived. | Correction made after those results had been seen; conclusions unchanged (Note 1 text) |
| D11 | 13 Nov | Proteome-supported centipede toxins versus control classes | The S- and L-class differences were recomputed with the 143 proteome-supported centipede toxins. S class: 0.69 (90% CI −0.03 to 1.57) for all members (undetermined) and 0.77 (95% CI 0.03–1.93) for domain-bearing members (toxins more clustered); L class not estimable. | Post hoc, after all panel results had been seen; descriptive only |
| D12 | 13 Nov | Title | The plan required the T1 title because the order-of- | Decided after all results had been |

| Entry | Date (2026) | Subject | Content | Timing |
|---|---|---|---|---|
| | | | recruitment analysis met its stopping rule. The authors chose a title that extends T1 from four bilaterians to the 14 origins of the panel and omits the comparison with secreted families; the abstract states the heterogeneity and the prediction interval. | seen |
| D13 | 13 Nov | Non-venomous controls and evidence-supported subset (post hoc) | Prompted by an internal pre-submission review. For each group of the venom panel, one animal outside all venomous lineages of the reference table was chosen by the eligibility rules and order of choice of the representative species; ToxProt homologues were analysed with the same procedure. End point: difference in log odds ratio between venomous origins and controls (meta-regression with the Hartung–Knapp adjustment), classified with the bound of ±0.5. Eight controls were chosen (no | Rules, choice of species and end point registered before any control genome was downloaded |

| Entry | Date (2026) | Subject | Content | Timing |
|---|---|---|---|---|
| | | | annelid was eligible). Result: controls pooled 2.10 (1.33–3.32); difference 0.39 (95% CI −0.12 to 0.90), undetermined; the six origins with proteome or expression evidence gave 2.70 (1.46–5.02). | |
| D14 | 13 Nov | Title and central claim | After D13, the D12 title was replaced by the present title, which states that toxin-associated genes cluster beyond family-size expectations in both venomous and non-venomous animals. Registered verdicts are unchanged; the abstract states that whether the excess is larger in venomous lineages remains undetermined. | Decided after all results, including D13, had been seen |

#### Effect of the correction

With the earlier assembly, the centipede odds ratio was 4.18 (2.60–7.48) and the pooled odds ratio across origins was 2.86 (1.85–4.41; prediction interval 0.96–8.48). With the assembly reported here, the centipede odds ratio is 3.82 (2.57–6.05) and the pooled odds ratio is 2.84 (1.85–4.36; prediction interval 0.97–8.33). The pooled S-class difference was 0.147 (90% CI −0.120 to 0.413) before and 0.146 (−0.115 to 0.408) after the correction, and the classification did not change. For the order of recruitment, the centipede main set comprised 45 families on the earlier assembly and 51 on the assembly reported here; none was orderable in either case, and the stopping rule applied in both (4 of 57 and 4 of 63 families orderable).

#### Completion status

All registered prospective analyses were run, with two exceptions. In the order of recruitment, the spider and scorpion genomes contributed no family, because no published venom proteome was available for these species. Evidence tiers in the four comparator genomes of the earlier analysis were recomputed only for the centipede. The strict tier was not estimable in any added species. The L control class was estimable in too few origins to be pooled. The order of recruitment met its stopping rule (4 of 63 families orderable).

#### Deviations recorded in the earlier analysis

The earlier five-genome analysis had its own analysis plan and deviation record, reproduced below without change.

Codes refer to elements of the analysis plan: Q1, family-size-matched clustering; P1, decomposition and composition of the excess; P3, local duplication topology; P4, expression endpoints; P5, repeat environment; P6, congener comparison; H3, an exploratory hypothesis on coordinated expression; E1, the pre-specified endpoint of the congener comparison.

| **Pre-specified element** | **Status** |
|---|---|
| Q1: family-size-matched clustering, three family definitions, family- and spatial-block intervals | Completed; primary family definition changed (next row); spatial-block intervals from the deterministic re-run (row after) |
| Q1: primary gene-family definition | Deviation. The frozen plan named the unidirectional-coverage definition as primary and the canonical-isoform definition as a sensitivity analysis, while citing the latter for intervals. The paper uses the canonical-isoform definition as primary, because it represents each gene once by its canonical isoform and was the definition with family-block intervals in the earlier |

| Pre-specified element | Status |
| --- | --- |
| | implementation. All three definitions are reported in full (Table S18). The spider and sea anemone estimates are lower under the canonical definition than under the unidirectional one, so the change does not systematically favour the conclusion |
| Q1: reproducibility of spatial-block intervals | A source of non-determinism in the spatial resampling was removed and the analysis was re-run (on the earlier assembly, for example, centipede 2.74–7.29 to 2.68–7.26, with identical point estimates). All values reported here come from the deterministic version |
| Q1: symmetric quality filter (overlapping models, gap proximity) in the centipede | Completed. The gap-proximity filter initially removed no gene, because gap and gene coordinates used different sequence names. After correction, it removes seven genes, none a toxin (canonical odds ratio 3.60 to 3.62; unidirectional 3.68 to 3.70). The additional filter on structurally anomalous models (coding length below half the family median, incomplete ends) was not applied |
| Q1 sensitivity: windows of 25, 50 and 200 kb; centipede Tier A and Tier B; matching on within-family sequence similarity; snake author-annotated venom-gene sets | Completed as specified (Table S24) |
| Q1: subset restricted to proteome-supported toxins | Not performed, because it would select toxins and controls asymmetrically |
| P1: decomposition, concentration null, removal of the five largest contributors, neighbour classes, homology of cross-family neighbours | Completed; removal was done with recomputed matching only, not with fixed matched sets |
| P3: local duplication topology | Completed; families with alignments shorter than 100 amino acids are excluded as specified (this removed two |

| Pre-specified element | Status |
|---|---|
| | evaluable snake toxin families) |
| P3: matching on sequence similarity, family age or duplication span as sensitivity analyses | Not performed (similarity matching was applied to Q1; see above) |
| P3: support threshold UFBoot ≥ 80 | Completed on the same trees (main text). The delta80 column of Table S21 was computed only for families with a UFBoot ≥ 95 cherry and is superseded by this re-run |
| P4: endpoints 1 and 2, RNA and protein, Holm correction across species within layer and endpoint | Completed |
| P4: sensitivity using toxins defined independently of expression evidence | Not performed |
| H3 (exploratory): matching of control pairs on distance, sequence identity and expression level | Not performed; controls are tandem pairs from size-matched non-toxin families only |
| Family-level link between the excess and expression endpoints | Not performed; only the link with $\Delta f$ was computed |
| P5: repeat environment | Stopped under the pre-specified rule (no duplication boundaries); reported descriptively in Note 7 |
| P6: congener comparison | The primary matched comparison was evaluable (14 toxin arrays from 12 families) and is reported in the main text. On the earlier assembly and annotation, it fell below the information threshold, which prompted the post hoc stratified comparison without the family-size restriction and excluding regions with assembly gaps; that comparison is reported as exploratory |
| Independent re-assembly from the same reads | Completed (Note 2). After the pre-specified analyses had been run on the provider assembly, the re-assembly replaced it as the reference, with a new annotation, and the analyses were re-run on the new reference and |

| Pre-specified element | Status |
|---|---|
| | annotation. Exceptions, code corrections and the results that changed are listed in Note 2 |

### Note 2. Centipede genome, toxin catalogue and venom-gland proteome

#### Methods

##### *Sampling, ethics and vouchers*

Adult *Scolopendra mutilans* (Chilopoda: Scolopendridae) were collected in Luoshan County, Xinyang, Henan Province, China, in 2023. They were identified morphologically, maintained at room temperature and fed raw chicken meat. No voucher specimen was deposited. Genomic DNA for the assembly was extracted from legs and muscle (isolate S-Q-1; sex not determined, recorded as "pooled" in the BioSample record). For transcriptome sampling, venom glands (V) and a non-venom tissue (T) were dissected at rest, immediately after venom extraction by mild electrical stimulation of the body (0 h), and 1 h later, with three replicates per tissue and time point (18 samples). The two tissues served as reciprocal controls. The tissues were snap-frozen in liquid nitrogen and stored at −80 °C until use. *S. mutilans* is an invertebrate not listed under Chinese wildlife-protection regulations, and no collection permit was required. Under the Regulations of the People's Republic of China for the Implementation of Laboratory Animal Management (State Council Decree No. 2, revised 2017) and the national standard GB/T 35892-2018, ethical review is not legally mandated for invertebrate research. Notwithstanding this exemption, all animal procedures were reviewed and approved by the Laboratory Animal Welfare and Ethics Committee of Pu'er University (Approval No. PEU-AEC-2023-119).

##### *Genome sequencing*

Library preparation and sequencing were carried out by Shanghai Biozeron Biotechnology Co., Ltd. (Shanghai, China), whose delivery report is dated 2024-06-04. Short-insert whole-genome libraries were built with the Illumina TruSeq Nano DNA Sample Prep Kit from 1 μg of genomic DNA. The DNA was sheared to 300–500 bp on a Covaris M220, end-repaired, 3′-A-tailed and ligated to indexed adapters. The libraries were enriched by 8 PCR cycles, size-selected on 2% agarose, quantified by PicoGreen (TBS380) and sequenced as 2 × 150 bp (insert size 450 bp). For HiFi sequencing, genomic DNA was fragmented (G-tube or Megaruptor), repaired and ligated to hairpin adapters to form SMRTbell templates. Linear and internally damaged circular molecules were digested enzymatically, and the templates were size-selected (BluePippin or Sage ELF) and sequenced in CCS mode. Hi-C libraries were prepared by formaldehyde cross-linking, restriction digestion to leave sticky ends, biotinylated fill-in, proximity ligation, protease de-cross-linking, shearing to 500–700 bp and streptavidin-bead capture, followed by short-read sequencing.

Short reads were quality-trimmed with Trimmomatic v0.39 (2). Adapters and non-ACGT 5′ bases were removed, read ends below Q20 were trimmed, reads with ≥ 10% N were removed, and fragments shorter than 75 bp after trimming were discarded. HiFi polymerase reads shorter than 200 bp or of quality below 0.80 were discarded, CCS reads were extracted with adapter removal, and CCS reads shorter than 200 bp were discarded. Assembly input comprised one PacBio Sequel II HiFi whole-genome run (delivered as three FASTQ parts), one Illumina NovaSeq 6000 Hi-C run and one Illumina NovaSeq 6000 short-read WGS run. A PacBio Sequel II Iso-Seq (FL-cDNA) run and 18 NovaSeq 6000 small-RNA runs were generated for annotation and regulatory analyses only (22 runs in total; SRA accessions SRR39514747–SRR39514768 under BioProject PRJNA1462551). The provider report gives 51.82 Gb of HiFi data (read N50 20,744 bp, mean 20,127 bp, longest 69,539 bp). The HiFi read set analysed in this study comprised 2,631,872 reads, and the two sources could not be reconciled to a single read set. Clean short-read whole-genome data amounted to 32.50 Gb (Q20 97.63%, Q30 94.72%), and clean Hi-C data to 92.78 Gb (Q20 98.36%, Q30 95.69%). These data correspond to 34.8×, 21.8× and 62.4× of the 1,488.0-Mb non-N assembly span for HiFi, short-read and Hi-C data, respectively (119.0× in total). The provider's 21-mer estimate of genome size (976.3 Mb; heterozygosity 0.51%; repeat content 55.3%) lies below the other k-mer-based estimates (1.23–1.49 Gb; Note 2).

#### *Transcriptome sequencing*

According to the provider's methods description, total RNA was extracted with TRIzol reagent (Invitrogen) and treated with DNase I (TaKaRa), and RNA quality was assessed with a 2100 Bioanalyzer (Agilent) and an ND-2000 spectrophotometer (NanoDrop). Libraries were prepared from 1 μg of total RNA with the Illumina TruSeq RNA Sample Preparation Kit by poly(A) selection, size-selected for 200–300-bp cDNA fragments, amplified for 15 PCR cycles and sequenced as 2 × 150 bp on an Illumina NovaSeq 6000 by Shanghai Biozeron Biotechnology. Each of the 18 libraries yielded 5.77–6.93 Gb of clean data. Reads were trimmed with Trimmomatic v0.36 (SLIDINGWINDOW:4:15, MINLEN:75), aligned to the provider assembly with HISAT2 using default parameters and counted per gene with HTSeq, and TPM values were computed by the provider on the earlier annotation.

#### *Genome assembly and quality*

Contigs were assembled from PacBio HiFi reads with hifiasm v0.25.0-r726 (3, 4) in Hi-C-integrated phasing mode with default purging of haplotigs. The primary contigs were scaffolded with YaHS v1.2.2 (5) using Hi-C reads mapped with chromap v0.3.2 (6) (preset hic; PCR duplicates removed; contacts with MAPQ < 10 discarded). YaHS contig error correction was disabled, because when enabled it introduced 543 contig breaks, of which only 27 were supported by HiFi reads. One mis-join was identified jointly from the Hi-C contact map, whole-genome alignment to the provider assembly (minimap2; ref. 7) and HiFi read-back. It was corrected by breaking contig ptg000010l at 21,466,000 bp and appending the 21.5-Mb segment to the scaffold of the chromosome to which it aligns. The 14 chromosome-scale scaffolds were named after the chromosomes of the provider assembly by whole-genome alignment and oriented as in that assembly (10 were reverse-complemented); the unplaced sequences keep their scaffold names. Contaminant sequence was identified with FCS-GX v0.5.5 (8). Thirteen unplaced scaffolds were removed and terminal segments of four were trimmed (4.71 Mb in total). Ten intervals within scaffolds assigned to *Spiroplasma* were retained after read-level verification (Note 2; Table S14).

Completeness was assessed with BUSCO v6.1.0 (9) (arthropoda_odb10, $n$ = 1,013; genome mode). As an independent check, compleasm v0.2.8 (10) was run with arthropoda_odb12; its results are reported separately because the tool and lineage database differ and the two are not directly comparable. Consensus accuracy was assessed with meryl/merqury (r977, $k$ = 21) (11), and small-scale and structural errors with Inspector (12). Terminal telomeric (TTAGG)n arrays were detected with a purpose-written scan (Supplementary Code). An earlier chromosome-level assembly of the same reads produced by the provider (Hifiasm and ALLHiC, versions not stated; 979.96 Mb) is compared in Note 2 and Table S25. A previous version of this study was based on that assembly.

#### Genome annotation

Repeats were modelled with RepeatModeler2 (13) and soft-masked with RepeatMasker 4.2.3 (rmblast 2.14.1). Genes were predicted with BRAKER3 v3.0.8 (14) in ETP mode from the soft-masked assembly, RNA-seq alignments (HISAT2; ref. 15) of the three venom-gland libraries whose raw reads were available, and OrthoDB v12 Arthropoda proteins (16). GeneMark-ETP and AUGUSTUS predictions were combined by TSEBRA (17), giving 14,743 genes. Helixer (18) predictions without a BRAKER3 counterpart on the same strand were added when supported by RNA-seq reads (≥ 10 fragments) together with eggNOG orthology or intron support, or when they recovered a BUSCO gene missing from the BRAKER3 set (1,935 genes after removing three transposase models). At toxin loci, gene models of the earlier annotation were projected onto the assembly (miniprot; ref. 19). These projections were used to split seven merged models into 17 and to add 15 missing models (none with an internal stop codon). The final annotation (Smutilans v3.1) contains 16,703 genes and 20,529 transcripts (BUSCO protein mode with the longest isoform per gene, arthropoda_odb12: 98.9% complete, 7.0% duplicated). Functional annotation used InterProScan 5 (20) (92.1% of transcripts with a hit) and eggNOG-mapper v2 (21). Genes of the earlier annotation were related to v3.1 genes by reciprocal best protein hits and by locus overlap of their projections. This correspondence table was used only to transfer expression values (see below) and to compare the toxin catalogues. The analyses used this annotation of 16,703 genes. Two gene models on a trimmed contaminant segment are absent from the deposited version (16,701 genes). Neither is a toxin or a member of a gene family used for matched controls, and excluding them leaves the family-size-matched odds ratio unchanged (3.82).

#### Toxin identification and curation

Predicted proteins were searched against UniProt ToxProt with DIAMOND v2.2.2 blastp (22) ($E < 1 \times 10^{-5}$), and hits were intersected with venom-gland-enriched expression (18-

sample RNA-seq; ratio ≥ 2, mean ≥ 5 TPM). Raw reads were available for only three of the 18 libraries. Expression values (provider TPM of the earlier annotation) were therefore transferred to v3.1 genes through the correspondence table. Genes matched by both reciprocal best hit and locus, by either alone, or as fragments of an earlier gene received that gene's values. When several earlier genes matched one v3.1 gene, the one with the highest mean venom-gland TPM was used (12,775 of 16,703 genes received values). ToxProt homologues without transferable expression (191) cannot be assessed for enrichment. The 74 of them with at least 5 TPM in the three re-quantifiable venom-gland libraries are listed as candidates in Table S6 and were excluded from all analyses. Signal peptides were predicted with DeepSig v1.2.5 (23) in eukaryote mode, and Pfam domains were annotated with HMMER v3.4 (24) against Pfam-A (ref. 25; 30,134 profiles). Candidates with a non-toxin best domain matching ToxProt over < 30% of the alignment were removed (six), yielding 188 toxins (Table S6).

##### *Venom-proteome re-mapping*

Four *S. mutilans* shotgun MS/MS proteomes (8,480 de-duplicated peptides) were mapped to the 188 toxin proteins by exact substring matching. One of them (B2; 879 de-duplicated peptides) is the published venom proteome (ref. 26; iProX IPX0001117000). Its identity was inferred from the exact agreement of its peptide count, and of its 52 peptide-compatible loci on the earlier annotation, with the published figures. The other three (the venom-gland peptidome and the torso proteome and peptidome) were generated in the same project and deposited in the same iProX project, but were not analysed in that publication. Sample identity (venom or body tissue) was inferred by intersecting each sample's protein identifiers with the venom (86 proteins) and body-tissue (784 proteins) GO reference sets supplied with the vendor report, and the larger fraction was taken as the call. The two sets share 51 identifiers, so the call rests on the difference between the two fractions and not on either fraction alone. The figures for the published sample alone (51 peptide-compatible and 34 locus-resolved toxins; 52 and 37 on the earlier annotation) are given in Table S7 for comparability with the earlier report. An I/L-equivalence sensitivity analysis on the published sample yields 53. Toxins were classed as peptide-compatible (≥ 1 matching peptide), locus-resolved (≥ 1 peptide not shared with any other of the 188 toxin proteins) or protein-group-level only. The published peptide-level FDR of 1% is cited, and no protein-level FDR is claimed for the re-search.

#### *Venom-gland DIA proteome*

Twelve adult *S. mutilans* were assigned to two treatments of six animals (three males and three females each): venom extraction by electrical stimulation 24 h before dissection, or no stimulation. The animals were collected in Luoshan County, Xinyang, Henan Province, China, on 3 September 2026 and maintained as described above for the genome animals (room temperature; fed raw chicken meat). Sex was assigned morphologically among animals of the same age and batch (females have a broader dorsum and a longer body) and was not confirmed anatomically. Venom had not been extracted from any animal before. Venom was extracted by stimulating the forcipules with a 3 V alternating current as described previously (26), and unstimulated controls were handled and sampled in parallel. Animals were chilled on ice without anaesthetic and killed by decapitation. The ethics approval given above (PEU-AEC-2023-119) covers the sampling and euthanasia procedures used for these animals. Both venom glands were dissected, snap-frozen in liquid nitrogen and stored at −80 °C.

Protein extraction, digestion and LC-MS/MS were performed by a commercial provider (Shanghai Biozeron Biotechnology). Tissue was ground in liquid nitrogen and lysed in 8 M urea, 1 mM PMSF and 2 mM EDTA with sonication. Protein (100 μg) was reduced (5 mM DTT, 37 °C, 45 min), alkylated (11 mM iodoacetamide, 15 min, dark), digested overnight with trypsin (Promega V5280) and desalted on C18. Peptides (200 ng) were separated on a Vanquish Neo UHPLC (PepMap Neo trap, 300 μm × 5 mm; Easy-Spray PepMap Neo column, 150 μm × 15 cm, 2 μm, 55 °C; 2.5 μL/min; 6.9-min effective gradient, 8 min per run). They were analysed on an Orbitrap Astral mass spectrometer in DIA mode (MS1 380–980 m/z at 240,000 resolution, normalised AGC target 500%, maximum injection time 5 ms; 299 isolation windows of 2 Th, HCD 25%, normalised AGC target 500%, maximum injection time 3 ms). Raw files were searched with DIA-NN 2.7.0 (27) in library-free mode, with a spectral library predicted in silico from the database in a separate step. The database comprised the 16,703 primary predicted proteins (one per gene), 12 curated toxin isoforms (16,715 entries; identical sequences were kept as separate entries, so tandem-cluster members remain distinct) and the 116-

entry cRAP contaminant set. Search settings were trypsin/P with at most one missed cleavage; peptides of 7–30 residues; precursor charge 1–4 and m/z 380–980; carbamidomethyl-Cys as a fixed modification; Met oxidation, protein N-terminal acetylation and protein C-terminal amidation as variable modifications (at most two per peptide); N-terminal Met excision; match-between-runs; and precursor and protein-group FDR of 1%.

For toxin mapping, stripped peptide sequences were matched to all database sequences by exact substring search, with isoleucine and leucine treated as equivalent. A toxin was peptide-compatible if at least one peptide matched it, and locus-resolved if at least one peptide matched that gene and no other gene in the database. The same code was applied to the 8,480 shotgun peptides under the earlier rules (exact matching, longest isoform per gene) and the earlier annotation. It reproduced the earlier pooled four-sample re-mapping of this study (60 peptide-compatible, 42 locus-resolved and 6 clusters with two or more resolved members), and on v3.1 it gave 60, 37 and 6 (Table S17). For clusters with two or more locus-resolved members, pairwise protein identity of those members was computed from global alignments (Biopython 1.87 PairwiseAligner; BLOSUM62, gap opening −10, extension −0.5; identical residues divided by the length of the shorter protein; ref. 28). The unique peptides of each member were compared with the other copy to count differing positions. Differential abundance was analysed with MaxLFQ protein intensities (log2, median-normalised, no imputation), restricted to proteins quantified in at least two of three animals in every sex × treatment group. limma (29) was used with the additive model ~ sex + treatment and robust empirical Bayes, and *P* values were BH-adjusted. The toxin-set test used camera (30) with the inter-gene correlation estimated from the data. Analysis plans, including acceptance criteria, were fixed before the results obtained with the earlier annotation were examined; at that stage, the pre-specified analysis used the provider's protein matrix. Because the provider's search used the earlier protein database, the present version reports the local search only.

#### Results in detail

The *S. mutilans* genome was assembled from PacBio HiFi and Hi-C reads with hifiasm in Hi-C phasing mode, and the primary contigs were scaffolded with YaHS. After removal of bacterial contaminant sequence, the assembly comprised 1.49 Gb in 475 sequences, of which 14 chromosome-scale scaffolds spanned 949.5 Mb (Table 1). The contig N50 was 50.0 Mb (482 contigs). The Merqury QV was 68.8, and 98.6% of BUSCOs were complete; both values were measured before the final one-contig correction and contamination cleaning. The assembly is gap-poor, with seven gaps of 100 bp (five of them on four chromosomes), but it is not telomere-to-telomere. One mis-join was identified jointly from the Hi-C contact map, whole-genome alignment to an earlier assembly of the same reads and HiFi read-back, and was corrected. The chromosomes were named and oriented after that earlier assembly. Ten intervals of *Spiroplasma*-like sequence within scaffolds (1.11 Mb in total) are joined to centipede sequence by continuous HiFi reads. They were retained as probable integrated sequence, and none of them contains a toxin gene (Note 2). The 538 Mb outside the chromosome-scale scaffolds has almost no Hi-C signal. On our reading of the k-mer spectra, this is repeat-rich unplaced sequence whose copy number and haplotypic redundancy remain unresolved (Note 2). An earlier assembly produced by the sequencing provider from the same reads (0.98 Gb; QV 65.9) was used in a previous version of these analyses. The conclusion of the family-size-matched comparison was the same on that assembly, whereas several secondary results changed (Note 2; Table S25). All 188 curated toxin genes lie on the chromosome-scale scaffolds.

To define the venom arsenal, the 20,529 predicted proteins of the 16,703 annotated genes were searched against UniProt ToxProt (31). Candidates were required to show venom-gland-enriched expression (venom-gland/control ≥ 2; mean ≥ 5 TPM), and six physiological proteins that matched ToxProt only through a shared domain were removed. A total of 188 curated toxins were obtained (Table S6). Of these, 121 had a predicted signal peptide (Tier A), and 67 had a toxin or venom-enzyme domain or scoloptoxin homology but no detected signal peptide (Tier B). Raw reads were available for only three of the 18 RNA-seq libraries (all three from venom glands). Expression values were therefore transferred from the earlier annotation by reciprocal best hits and locus overlap (Materials and Methods). A further 74 ToxProt homologues are transcribed in the venom gland, but their expression could not be transferred. They are listed separately as candidates and were excluded from all analyses. Twenty-eight of the 188 genes derive from 12 genes that the earlier annotation modelled as single genes. All 28 begin with methionine, 24 carry a predicted signal peptide and 19 are resolved by their own genome-unique venom-gland peptides, consistent with tandem copies that the earlier annotation had merged (Table S6, Part B). When these genes were reduced to one locus per earlier gene (172 toxin loci), the family-size-matched odds ratio was 3.27 (95% family-block interval 2.18–5.22; Table S18). The most common Pfam domains were serine protease (15 toxins), serpin (12), γ-glutamyltransferase (11), CAP/CRISP (9) and Kazal (5). In addition, 118 of the toxins lie in 32 tandem clusters (toxins ≤ 100 kb apart; Fig. S5).

A data-independent-acquisition (DIA) proteome of dissected venom glands was obtained from 12 adults (three males and three females per treatment; venom extracted by electrical stimulation 24 h before dissection, or not stimulated). A total of 6,687 non-contaminant protein groups were identified (1% FDR). Peptides were recovered for 163 of the 188 toxins, and 141 of them were resolved to a single locus by peptides found in no other predicted protein (Table S17). Four earlier shotgun proteomes, pooled and analysed with the same rule of uniqueness in the whole predicted proteome, gave 60 peptide-compatible and 37 locus-resolved toxins (Table S17). Among them, the published collected-venom proteome gave 51 peptide-compatible and 34 locus-resolved toxins when uniqueness was assessed among the 188 toxin proteins (Table S7). In 25 of the 32 clusters, at least two members are each resolved by their own genome-unique peptides. Every one of these clusters contains a pair of resolved members below 95% protein identity, and each member of the pair carries a unique peptide that differs from the other copy at two or more positions. These clusters therefore contain distinct translated copies, a pattern not explained by near-identical alleles of a single locus. The proteome supports their translation, while their physical arrangement rests on the assembly.

Protein abundance of locus-resolved toxins was correlated with gland transcription (Spearman $\rho = 0.56$, $P = 7.8 \times 10^{-13}$, $n = 141$). The 25 toxins without peptides were mostly weakly transcribed (median TPM 30.3 versus 294.1). The exception was one toxin with high transferred TPM (SMUT163040, 2,112 TPM), a fragment of an earlier merged model (Note 2). No protein differed detectably between treatments or sexes (limma, BH-adjusted $P \geq 0.59$). With three animals per sex and treatment, the size of effect that can be excluded is not defined.

#### Assembly, contamination screen and quality

##### *Assembly*

The reference assembly was built from the PacBio HiFi and Hi-C reads of isolate S-Q-1 (Materials and Methods). The BioSample records the sex as pooled, and a single-individual origin of all libraries is not documented. Primary contigs were produced by hifiasm v0.25.0 in Hi-C phasing mode and scaffolded by YaHS v1.2.2 without contig error correction. With correction enabled, YaHS introduced 543 breaks, of which only 27 had support from HiFi reads. One mis-join was identified jointly from the Hi-C contact map, whole-genome alignment to the earlier provider assembly (Note 2) and HiFi read-back. It was corrected by breaking one contig at 21,466,000 bp and joining the 21.5-Mb segment to the scaffold of the chromosome to which it aligns. The 14 chromosome-scale scaffolds were named after the chromosomes of the provider assembly by whole-genome alignment and oriented as in that assembly (10 were reverse-complemented). Chromosome numbers are therefore comparable between the two assemblies, but coordinates are not. The Hi-C contact map of the corrected chromosomes is shown in Fig. S3a.

After the contamination screen described below, the assembly spans 1,488,016,704 bp in 475 sequences (482 contigs; contig N50 50.0 Mb), of which the 14 chromosome-scale scaffolds span 949.5 Mb (63.8%). It contains seven gaps of 100 bp, five on chromosomes (Chr3, Chr6, Chr8 and two on Chr13) and two on unplaced scaffolds. Consensus quality, k-mer completeness, gene-set completeness and error counts were measured on the scaffolds before the one-contig correction and the contamination cleaning, which together changed one junction and removed 4.71 Mb (0.32%) of sequence. The Merqury QV was 68.8. The k-mer completeness was 92.6% and rose to 99.1% for the union of the two phased haplotype assemblies, so the shortfall of a single assembly reflects haplotype-specific k-mers. BUSCO completeness was 98.6% (0.6% duplicated; arthropoda_odb10, $n$ = 1,013), and Inspector reported 6.9 small-scale errors per Mb and 62 structural errors.

HiFi reads mapped back to the final assembly give a primary alignment for 99.86% of reads (2,628,094 of 2,631,872).

##### Genome size and unplaced sequence

k-mer analysis of the reads gives discordant haploid size estimates: 1.23 Gb at 0.40–0.45% heterozygosity from the HiFi reads, and 1.45–1.49 Gb at 1.33–1.81% from the Illumina survey reads. Both estimates have modest model fit (Note 2), so we use them only as supporting information. The assembly (1.49 Gb) lies at their upper end, whereas the provider assembly (0.98 Gb) lies below all of them. On our reading, the 538.5 Mb outside the chromosome-scale scaffolds is mainly repeat-rich, and whether it consists of further repeat copies or partly of redundant haplotype remains unresolved. Relative to the provider assembly, k-mers present in the reads but absent from the assembly fall only from 44.5 to 40.6 million. This is far fewer than the ~40 million haplotype-specific k-mers expected if the extra sequence were a redundant haplotype at 0.4% heterozygosity. In contrast, k-mers at an assembly copy number above four rise from 14.5 to 17.5 million. This sequence also has almost no Hi-C signal and no counterpart with MAPQ ≥ 5 in the provider assembly. This interpretation is an inference, and the additional copies were not validated individually. All 188 curated toxin genes lie on the chromosome-scale scaffolds.

##### Telomeres

With the telomeric repeat survey described below, terminal (TTAGG)*n* arrays were detected at 8 of the 28 chromosome ends: the 3′ ends of Chr1, Chr5, Chr7, Chr8, Chr9, Chr10 and Chr14 and the 5′ end of Chr3 (array lengths 0.7–3.9 kb; Table S5). One further array, 17.6 kb internal to the 3′ end of Chr12, was classed as subtelomeric-ambiguous. No chromosome carries detected arrays at both ends, and internal control windows yielded no detections. The assembly is therefore not telomere-to-telomere. Run with unchanged code on the provider assembly, the survey reproduces the earlier result (5 of 28 ends; Note 2).

##### Contamination screen

FCS-GX flagged 30 sequences or intervals, all assigned to bacteria (Table S14). Thirteen unplaced scaffolds (1.83 Mb in total; 45–557 kb each), assigned to Firmicutes over 40–

100% of their length, were removed. Terminal segments of four unplaced scaffolds (2.88 Mb in total; 99 kb to 1.40 Mb), assigned to Mollicutes, were trimmed. Two gene models on the trimmed segment of scaffold_26 (SMUT165800 and SMUT165810, neither a toxin) were removed with it, leaving 16,701 genes and 20,527 transcripts in the deposited annotation. The analyses in this paper used the annotation before cleaning (16,703 genes). The two removed genes are not in the toxin catalogue and belong to no gene family used for matched controls, and excluding them leaves the family-size-matched odds ratio unchanged (3.82). Three sequences flagged for review, each with 11–24% contaminant coverage (unplaced scaffolds of 27 and 142 kb and a 4.2-kb interval on Chr11), were retained unchanged.

The remaining ten intervals (1.11 Mb in total; 2.5–231 kb) lie on Chr1, Chr2, Chr6, Chr10, Chr13 (four intervals) and two unplaced scaffolds. They were assigned to *Spiroplasma* (Mollicutes) and flagged as contaminant sequence internal to a scaffold. We tested each interval with the three checks applied earlier to the single such interval of the provider assembly, on its Chr9 (Table S14). These were read depth against 200 length-matched host intervals of the same sequence, alignment to Mollicutes reference sequences, and HiFi reads spanning each of the 20 junctions with host sequence. Each of the 20 junctions is spanned by reads with MAPQ ≥ 20 (14–81 reads against control medians of 15–35; the two scaffold_24 junctions carry more reads than their controls). Thus, none of the junctions shows the loss of read support expected at a chimeric join. Depth over seven intervals is indistinguishable from the host (robust $z$ −0.4 to 0.8). The Chr2 interval has 1.26-fold and the Chr13 interval at 17.6 Mb 2.74-fold the host median depth. The scaffold_24 interval has 1.96-fold, with its reads uniquely placed (depth 82.0 at MAPQ ≥ 20, against 19.1 for the host intervals of that scaffold), and the Chr10 interval has host-level depth over all reads but 0.71-fold at MAPQ ≥ 20. Only 28–34% (63–69 kb) of each of the five intervals of about 200 kb aligns to the Mollicutes references, at a median identity of about 85%, and none of the flanking host sequence does. The intervals contain 16 gene models: 13 with best eggNOG annotation at the level of Tenericutes (among them FtsZ, the GroEL chaperonin, topoisomerase IV subunit ParC and purine-nucleoside phosphorylase), two at the metazoan level and one without annotation. None of them is a toxin.

Sequence from a free-living contaminant would be expected to have a depth set by its own abundance and no read continuity with host sequence. Host-level depth over most intervals, together with continuous junction-spanning reads at every junction, supports these intervals being physically joined to centipede sequence in the sequenced material. We interpret them as *Spiroplasma*-derived sequence integrated into the nuclear genome. We therefore retained the ten intervals and their 16 gene models, and the gene models are listed in Table S15 as lying in candidate bacteria-derived intervals (13 with Tenericutes-like orthology, 2 metazoan-like and 1 unassigned). Two caveats apply. First, the data cannot exclude that the same individual also carried a *Spiroplasma* endosymbiont whose reads map to these intervals; the elevated depth of the Chr13 interval at 17.6 Mb and of the scaffold_24 interval may partly reflect such reads. Second, no interval was confirmed by PCR or in a second individual, and whether the five intervals of about 200 kb are copies of one insertion was not tested. None of the conclusions of the paper depends on these intervals.

#### *Methods*

Contamination screen. The scaffolds, before chromosome naming, were screened with FCS-GX v0.5.5 against the NCBI GX database (build of 2023-01-24, 3,025,824 sequences) under the *S. mutilans* taxonomy. The database was split into six shards, and alignments from all shards were merged before taxonomic classification and action calling. Sequences with the action EXCLUDE were removed and segments with the action TRIM were cut, and the AGP file and annotation were updated accordingly (gene models overlapping a removed segment were deleted). Intervals with the action FIX were examined as described next, and sequences with the action REVIEW were retained.

Verification of FIX intervals. HiFi reads were mapped to the cleaned assembly with minimap2 v2.31 (preset map-hifi, -I 8G) and sorted with SAMtools v1.23.1. Mean depth of each interval was computed with samtools bedcov over all reads and over reads with MAPQ ≥ 20. It was compared with the depth of 200 intervals of the same length drawn at random (fixed seed) from the same sequence outside all flagged intervals, as a robust *z* score (deviation from the control median divided by 1.4826 × the median absolute deviation). Interval sequences and equal-length flanking sequences on both sides were aligned to Mollicutes reference sequences extracted from the FCS-GX database (minimap2 preset asm20), and aligned bases and median identity were recorded. Reads spanning each junction between a flagged interval and host sequence were counted with the junction script used for the provider assembly (Supplementary Code), at MAPQ ≥ 20 and ≥ 0, against 200 random host positions as negative controls. Gene models overlapping the intervals were annotated with eggNOG-mapper v2.1.15 (eggNOG 5.0.2).

Telomeric repeat survey. Telomeric arrays were surveyed with a purpose-written scan of the assembly (chromosomes Chr1–Chr14 only; unplaced contigs excluded). Candidate motifs were derived from 5-, 6- and 7-mer frequency spectra of terminal 50-kb windows relative to internal background windows on a discovery subset (odd-numbered chromosomes), and only motifs with zero internal-background detections were retained. The selected motif (TTAGG) and all thresholds were then frozen before the remaining chromosomes were scanned. Detection required, in at least one 500-bp window (100-bp step), union base coverage ≥ 0.25 by the motif and its rotations and reverse complements, and ≥ 0.60 of adjacent matches being contiguous. Arrays were classified by distance to the physical sequence end as terminal (< 1 kb), partial (1–10 kb) or subtelomeric-ambiguous (> 10 kb). Thresholds were calibrated using all chromosomes, and successive script revisions were informed by earlier whole-assembly results. This survey is therefore reported as exploratory, and it has not been validated against long-read support for subtelomere–telomere junctions.

#### Toxin catalogue and proteome support

##### *Toxin catalogue*

To define the venom arsenal, we searched the 20,529 predicted proteins of the v3.1 annotation against UniProt ToxProt (31) and required venom-gland-enriched expression (venom-gland/control ≥ 2; mean ≥ 5 TPM, using values transferred from the earlier annotation; Materials and Methods). This search yielded 194 candidates. Homology alone can misassign large physiological proteins that share a single domain with a toxin, so we curated the set by architecture. Six genes whose best Pfam (25) domain was non-toxin (receptor kinases, immunoglobulin-superfamily proteins, an ankyrin-repeat protein and a teneurin-type EGF protein) and that matched ToxProt over < 30% of their length were removed as spurious, independent of expression level (Table S6). The resulting 188 curated toxins comprise 121 with a predicted signal peptide (Tier A, predicted to be classically secreted) and 67 bearing a bona-fide toxin or venom-enzyme domain, or homology to a characterised scoloptoxin, without a detected signal peptide (Tier B, including cysteine-rich peptide toxins). More toxins carry a detected signal peptide than in the earlier annotation (121 against 77).

Twenty-eight genes derive from 12 earlier models, each of which corresponds to several complete v3.1 models (Table S6, Part B). When these genes were reduced to one locus per earlier model, the catalogue has 172 loci, and the family-size-matched odds ratio is 3.27 (2.18–5.22). Of the 180 earlier toxins, 171 have a counterpart. The nine without one lack a ToxProt hit (three), are not venom-gland-enriched (one), could not be projected completely (two) or have no transferable expression (three). The set is dominated by trypsin-type serine-protease, serpin, γ-glutamyltransferase, CAP/CRISP, Kazal protease-inhibitor and phospholipase-A2 domains, together with EGF modules. None of the 188 models contains an internal stop codon (median length 215 aa). In addition, 180 begin with methionine, and the eight that do not are models projected from the earlier annotation at toxin loci (Materials and Methods). These features reduce, but do not exclude, concern about fragmentary or clearly inactivated models.

##### *Shotgun venom proteomes*

To support the catalogue at the protein level, we re-mapped our *S. mutilans* shotgun proteomes onto the 188 loci by exact substring matching. Four samples were available (8,480 de-duplicated peptides in total). Sample identity was inferred by intersecting each sample's protein identifiers with the vendor's venom and body-tissue GO reference sets. Two samples are venom (A2, 20 proteins, venom fraction 0.400; B2, 175 proteins, 0.486), and two are body tissue (C2, 135 proteins, body fraction 0.541; D2, 1,205 proteins, 0.649). B2 is the previously published venom proteome (ref. 26; ProteomeXchange/iProX IPX0001117000). Its 879 peptides match the published count exactly, and on the earlier annotation it alone recovers the 52 peptide-compatible loci reported previously. The other three samples are deposited in the same iProX project (IPX0001117000) but were not analysed in that publication. Pooled, the four samples give 60 peptide-compatible and 37 locus-resolved toxins when uniqueness is assessed in the whole predicted proteome (Table S17). Per-sample contributions are not tabulated and were not used in the analyses.

In the published sample, 199 of the 879 peptides map to the toxins. The remaining 680 are not expected to map, because the proteome samples the whole venom and includes enzymes, non-toxin secretory proteins and peptides from loci outside the curated toxin set. Re-mapping identified 51 toxins as sequence-compatible with venom peptides, of which 34 were locus-resolved by ≥ 1 peptide unique among the 188 toxin proteins. The remaining 17 were supported only at the paralogous protein-group level; that is, a shared peptide places at least one family member in venom but cannot resolve the individual locus (Table S7). This sample is collected venom (subjected to 3-kDa ultrafiltration) and not dissected gland. It therefore provides physical evidence that these peptides are present in venom, including support for a subset of the signal-peptide-negative Tier B toxins. Collected venom may nonetheless contain minor secretory-epithelial components, and peptide detection does not itself demonstrate toxicity or classical secretion. The 3-kDa cut-off also imposes a directional bias. Components below roughly 27 residues are removed, so the smallest cysteine-rich peptides, the class for which confirmation of Tier B assignments would be most useful, cannot be supported by this proteome. The 34 locus-resolved loci are therefore a lower bound with a known direction. Support is described as mutually exclusive layers: 51 peptide-compatible toxins (53 under I/L-equivalent matching; 34 of them locus-resolved); of the remaining 137, 92 signal-peptide-positive; and 45 supported by expression and homology or domain only. We cite the peptide-level FDR of 1% from the original study, and no protein-level FDR is claimed for this re-search.

##### *Venom-gland DIA proteome*

To test the catalogue in the tissue that synthesises venom, we generated a data-independent-acquisition (DIA) proteome of dissected venom glands from 12 adults. Each treatment comprised three males and three females, and glands were collected 24 h after electrical venom extraction or from unstimulated controls sampled in parallel. The proteome was searched against all predicted proteins with tandem-cluster members kept as separate entries (Materials and Methods). The search identified 6,687 non-contaminant protein groups (1% FDR; 6,009–6,270 per individual), and contaminant proteins contributed 0.04% of precursor signal. Peptides were then mapped to the 188 toxins with the same exact-matching procedure used for the shotgun data, which reproduces the earlier counts when applied to those data with the earlier annotation (Materials and Methods). The gland proteome recovers peptides for 163 toxins and resolves 141 to a single locus by peptides found in no other gene of the predicted proteome, compared with 60 and 37 from the four earlier shotgun samples. Of the locus-resolved toxins, 106 have no locus-resolved support in those samples. Together, the two datasets give 164 peptide-compatible and 143 locus-resolved toxins (Table S17).

The gland data also bear on whether the tandem clusters are genuine duplicates. In 25 of the 32 clusters, at least two members are each resolved by their own genome-unique peptides (23 when support is required in at least four of six animals in each treatment), compared with 6 clusters from the shotgun data. These members are unlikely to be allelic variants of a single locus. Across the 182 pairs of resolved members in these clusters, protein identity ranges from 20.0% to 97.8% (median 39.5%). Every one of the 25 clusters contains a pair below 95% identity in which each member carries a unique peptide differing from the other copy at two or more positions. This divergence far exceeds that expected between alleles at the provider's genome-wide heterozygosity estimate of 0.51%. Independent peptides for several members of the same cluster are therefore protein-level evidence, separate from the assembly itself, that these clusters contain distinct translated copies, although their physical arrangement rests on the assembly.

Proteome support was correlated with gland transcription. Among locus-resolved toxins, protein abundance (summed intensity of unique peptides, median across animals) was correlated with venom-gland TPM (Spearman $\rho = 0.56$, $P = 7.8 \times 10^{-13}$, $n = 141$). The 25 toxins without any matching peptide have lower gland TPM than those with peptides (median 30.3 versus 294.1; one-sided permutation $P < 0.001$). Only one of them lies in the top quarter of gland TPM: SMUT163040 (2,112 TPM), a Mu-scoloptoxin(15) homologue in a mixed scoloptoxin–serpin cluster on chromosome 14. In contrast, 28 locus-resolved toxins lie in the bottom quarter (Table S17). Not all toxin genes expressed in sea anemone tentacles are represented in the venom proteome (32). In this gland-tissue proteome, the toxins that escape detection are mostly the weakly transcribed ones, a pattern more consistent with detection limits than with untranslated, highly expressed genes. However, SMUT163040 is one of several fragments of an earlier merged model and inherits that model's TPM, which may belong to another fragment.

Finally, the two treatments showed no detectable difference. No protein differed detectably between glands collected 24 h after venom extraction and unstimulated glands, or between sexes (limma, BH-adjusted $P \geq 0.59$), and a pre-specified competitive test found no overall shift of the toxin set (camera $P = 0.22$). The largest axis of variation among individuals instead tracked the toxin share of total protein (1.6–13.8%; Spearman $\rho$ with PC1 = −0.83), which did not differ between treatments. With three animals per sex and treatment, the size of effect that can be excluded is not defined.

#### The earlier provider assembly and the results that changed with the reference

##### *Role of the provider assembly*

The pre-specified analyses were first run on a chromosome-level assembly delivered by the sequencing provider (Hifiasm and ALLHiC, versions not stated; 979.96 Mb), with the provider's annotation and a toxin catalogue of 180 genes. The earlier version of this study was based on that assembly. Because it could not be reproduced from recorded commands, we re-assembled the genome from the same reads (Note 2). The re-assembly was first used as an independent check (below). It then replaced the provider assembly as the reference, the genome was re-annotated, and the toxin catalogue was rebuilt (188 genes). The analyses were re-run after each script had been shown to reproduce its earlier output on the provider assembly, with three exceptions. First, two code defects were corrected (below). Second, the GGT tree was rebuilt with single-threaded alignment, because the multi-threaded alignment of the earlier tree is not reproducible. Third, the filtered-set rows of Table S16 for the four comparison species could not be recomputed and are carried over from the earlier run. The results that changed are listed at the end of this Note. This Note also keeps our verification of the provider assembly, written when it was the reference.

#### *Verification of the provider assembly: contiguity, telomeres and completeness*

The provider assembly spans 979.96 Mb across 14 chromosomes, with a contig N50 of 57.6 Mb (2.4-fold that of the published assembly; ref. 33; Table S25) and only 1.3 kb of remaining gaps across the 14 chromosome-scale scaffolds. Six of the 14 chromosomes (Chr1, Chr7, Chr10, Chr11, Chr12 and Chr14) are gap-free, while the remaining 8 carry 13 gaps of 100 bp each (1,300 bp of N in total; no gaps in the 12 unplaced sequences). Telomeric repeat arrays anchored at the physical sequence end (distance 0) were detected at 5 of the 28 chromosome ends: Chr5, Chr7, Chr9, Chr10 and Chr14, all at the 3′ end (array lengths 0.7–3.8 kb). One further array on Chr12 lies 17.6 kb internal to the end and was classed as subtelomeric-ambiguous. No chromosome carries detected arrays at both ends, and no array was detected at any 5′ end; internal control windows yielded zero false-positive arrays. The assembly is therefore gap-poor and highly contiguous but is not telomere-to-telomere in the strict sense.

To examine whether the ends without detected arrays are array-free or are assembly truncations, we re-mapped the HiFi reads and examined outward-facing soft-clips at every chromosome end. HiFi read-back identified coherent ≥ 1-kb outward extensions supported by at least three independent reads at seven chromosome ends, all at the 5′ ends of Chr1, Chr2, Chr3, Chr6, Chr7, Chr8 and Chr11. None met the pre-specified TTAGG-rich soft-clip criterion. Only 9 of 28 ends had sufficient edge-alignment support; 21 showed low terminal coverage and 3 were not evaluable. Thus, the pronounced 5′-end asymmetry indicates missing sequence beyond seven assembled endpoints, while limited read mappability prevents the exclusion of additional incomplete or telomeric ends (Table S5). The assembly is highly complete and accurate, with 98.7% complete BUSCOs (98.2% single-copy, 0.5% duplicated; 0.2% fragmented, 1.1% missing; arthropoda_odb10, $n$ = 1,013) and a consensus quality value of QV 65.9 (per-base error $2.5 \times 10^{-7}$). Contig-level statistics were computed by splitting the assembled sequences at runs of N (26 sequences and 13 gaps yield 39 contigs; contig N50 57,639,668 bp). The provider's structural annotation comprised 30,063 transcripts from 26,089 genes, and transposable elements occupy 50.1% of the genome.

Contig count alone need not determine whether short-range toxin clusters remain observable, so we tested this directly. Under a coordinate-based counterfactual, exactly 313 additional breakpoints were placed independently over the non-N sequence of the provider assembly while its coordinates and observed cluster membership were retained. The expected number of within-cluster adjacencies no longer lying on the same contig was 0.44 of 74 (0.6%), and the expected number of split clusters was 0.44 of 29. TE-weighted sensitivity models ($\lambda_{te}$ = 1, 3 and 10, treated as sensitivity models and not as bounds) yielded 0.45–0.46 for both quantities after rounding. None of the 29 clusters spanned any of the 13 existing within-sequence gaps. This coordinate-only counterfactual did not simulate reassembly, reannotation or gene truncation, and it does not estimate how many clusters the published assembly missed. Under these modelled breakpoint placements, the analysis provided little support for a substantial contig-count-driven gain in the observability of local toxin clusters. The consensus accuracy of the provider assembly is high (QV 65.9; Table S25). No consensus quality value is available for the published assembly, and the two BUSCO assessments are not directly comparable.

#### *Verification of the provider assembly: genome size, read-back and k-mer completeness*

The assembly was produced by a commercial provider that did not record software versions or command lines (Materials and Methods), so we re-verified it from the primary data. k-mer analysis of the HiFi reads estimates a haploid genome size of 1.23 Gb at 0.40–0.45% heterozygosity, whereas the Illumina survey data give 1.45–1.49 Gb at 1.33–1.81%. The two estimates differ by 18% in length and 3.7-fold in heterozygosity, and both have modest model fit (lower bounds 46% and 32%). We therefore use k-mer size estimates only as supporting information and not to adjudicate assembly content. Mapping all HiFi reads back to the assembly aligns 99.84% of reads with a primary alignment (2,627,694 of 2,631,872). Of all reads, 76.9% (2,024,362 of 2,631,872) align with MAPQ ≥ 20, and this MAPQ-filtered set was used for all depth analyses below. The k-mer completeness (merqury) is 91.93%, so about 8% of read-derived solid k-mers are absent from the assembly. We do not attribute this shortfall to a single cause. A pseudo-haploid assembly represents one haplotype, so k-mers spanning heterozygous sites on the other haplotype are absent by construction. Under a simple diploid distinct-k-mer model, the expected completeness is $1/[2 - (1 - h)^k]$, which at $k = 21$ and the HiFi heterozygosity estimate ($h$ = 0.40–0.45%) is about 91.7–92.5%, the same magnitude as the observed value. Heterozygosity is therefore sufficient in magnitude to account for much of the gap, although this is a model expectation and not a measurement. Unassembled or divergent repetitive sequence may contribute additionally, as unplaced contigs carry 1.3–3× the chromosomal read depth, but the present data cannot partition the contributions. We have not measured completeness for the previously published assembly, so no comparison is made.

#### Verification of the provider assembly: haplotypic duplication

Haplotypic-duplication analysis (purge_dups) flagged 68.0 Mb across 114 intervals and would remove 55.1 Mb, reducing the assembly to 924.8 Mb, closer to the 903.9 Mb reported for an independent *S. mutilans* assembly (33). BUSCO scores are identical to the digit before and after purging (C:98.7% [S:98.2%, D:0.5%], F:0.2%, M:1.1%, $n$ = 1,013), and consensus quality is essentially unchanged (QV 65.94 versus 65.80). However, neither metric is informative here. The duplicated fraction is already 0.5% and has no room to fall, and QV measures consensus base accuracy and not redundancy. The metric that does respond to haplotypic duplication is k-mer completeness. Purging lowers it from 91.93% to 91.68%, a loss of 1,562,859 solid k-mers. This loss is small relative to the 55.1 Mb removed, indicating that most of the removed sequence is represented elsewhere in the assembly. It is not zero, however, because the removed set contains solid k-mers absent from the retained assembly. These may be alternative-allele k-mers and not biologically unique sequence, so the loss bounds, but does not identify, what a blanket purge would discard.

#### Verification of the provider assembly: low-depth intervals and Hi-C

Interval-level read depth shows that the flagged set is heterogeneous (Table S9). Twenty-four intervals totalling 30.6 Mb lie below 0.6× of their chromosome's median depth. Among them are three chromosome-terminal blocks of 8.7–9.5 Mb at the starts of chromosomes 3, 4 and 9 (depth ratios 0.34–0.39). purge_dups annotates the largest of these as a haplotig of chromosome 1, an assignment that the Hi-C data do not support (see below). These ratios fall below the ~0.5 expected of simple allelic partitioning, consistent with the additional loss of multi-mapping reads described below, and 27.7 Mb of the 30.6 Mb is concentrated in the three terminal blocks. A Hi-C contact map built from 143.9 million read-pair contacts (chromap/PretextMap) provides independent evidence for the three terminal blocks. Each block shows essentially no off-diagonal contact with the body of its chromosome, whose dense, continuous diagonal signal begins only distal to the block, together with an internal checkerboard structure typical of unordered fragments. Other chromosome termini show no such break (Fig. S3b–e).

These observations must be read against a mappability confounder that applies to both the depth and the Hi-C analyses. If a retained haplotig were as similar to its counterpart as the genome-wide heterozygosity of ~0.4% would imply, HiFi reads would align equally well to both copies and be removed by the MAPQ ≥ 20 filter used here, and the same mechanism would depress Hi-C contacts. We have not measured the self-alignment identity of these three intervals, so this mechanism could operate but has not been demonstrated. The depth signal is in any case not independent of the purge_dups annotation, which itself combines a read-depth cutoff with self-alignment similarity. In addition, the coincidence of the Hi-C break with the depth boundary is expected under a pure mappability model and is not evidence against it. The checkerboard interior is a property of fragment ordering and not of signal strength, and is less readily explained by uniform signal suppression. However, we have not controlled for sparse-count artefacts and do not treat it as decisive.

As an exploratory check, we examined which chromosomes each block contacts in trans, normalising by partner chromosome length and comparing each block with an equally sized control interval from the middle of its own chromosome. All three blocks carry more trans signal than their controls (trans fraction 30–53% versus 19–24%; strongest partner enrichment 1.95–3.17 versus 1.19–1.36). Enrichment for chromosome 1 is 1.18–2.11, and chromosome 1 is not the strongest partner for two of the three blocks, so the data give no positive support for the purge_dups assignment to chromosome 1. Chr3-head and Chr4-head are reciprocally enriched (3.17 and 2.27), although enrichments involving Chr9-head are weaker (0.77–1.95), so this pattern is not uniform across the three blocks. We report this as an exploratory anomaly and not as evidence of origin. Low mappability changes relative as well as absolute contact counts, and degenerate alignment can bias towards homologous partners. A single mid-chromosome control does not match the blocks for telomere proximity, compartment or alignability, and we have neither corrected for selecting the maximum partner nor tested significance by permutation (Table S11). We therefore flag these three blocks (27.7 Mb) as candidate anomalous or mis-scaffolded terminal sequence.

Depth stratified by mapping quality and an independent re-assembly (below) distinguish two cases. In the Chr3 and Chr4 blocks, the median depth of reads with MAPQ ≥ 1 is 2.2 and 3.1 times that of the chromosome body. About 69–76% of the reads in these blocks have MAPQ 1–4, compared with about 95% of body reads at MAPQ 60. These two blocks are therefore best explained as collapsed multicopy (repeat) sequence, whose low depth at high mapping quality reflects multi-mapping. The Chr9 block is below body depth at every threshold and near half of it without a mapping-quality filter (median ratios 0.60, 0.57 and 0.37 at MAPQ ≥ 0, 1 and 20), as expected of haplotype-specific sequence.

In the reference, each block is separated from its chromosome by a 100-bp gap. In the re-assembly, none of the three is joined to the end of the corresponding chromosome, so their placement at these chromosome ends is a scaffolding decision that the re-assembly does not support, and where the blocks belong remains unresolved. The provider assembly retains this sequence unchanged, and it is flagged in Table S9. The remaining flagged intervals on chromosomes lie at or above 0.6×. Unplaced contigs as a class lie at 1.30× the chromosomal median (up to 2.95×), with GC indistinguishable from the chromosomes (0.2997–0.3295), consistent with collapsed repeats and not haplotypic redundancy.

The purge_dups removal set does not coincide with the low-depth set (it would also remove intervals at normal depth) and improves no measurable quality metric, so we did not apply it. The assembly is reported at 979.96 Mb, with the 30.6 Mb of sub-0.6× sequence disclosed as candidate anomalous sequence; the three largest blocks are interpreted as two collapsed repeats and one haplotype-specific segment. None of the 180 curated toxin genes and none of the 29 tandem clusters overlaps these candidate regions (0 of 180 and 0 of 29 by direct intersection). The toxin catalogue and the cluster definitions therefore do not depend on this uncertainty, and genes in these intervals remain part of the genomic background from which random and size-matched controls are drawn.

#### *Verification of the provider assembly: contamination and a horizontally transferred segment*

The predicted proteome was screened against metazoan (222,528 proteins) and non-metazoan (375,566 proteins) references, and the results were combined with the depth and GC distributions. No sequence met two or more contamination criteria (Table S8). This screen distinguishes clearly non-metazoan contamination but does not assign fine-grained taxonomy, so we also completed a full FCS-GX screen against the NCBI GX database (v0.5.5; 2023-01-24 build) using a split-database workflow required by memory limits (see Methods below). The official FCS-GX action report flags a single interval for correction, Chr9:6,948,379–6,990,380 (42,002 bp, 0.004% of the assembly), assigned to *Spiroplasma poulsonii* (prok:mycoplasmas), and no sequence for exclusion or trimming. Read depth across this interval (36.75×, full coverage) is indistinguishable from the host background of 200 size-matched control intervals on the same chromosome (median 41.55×, MAD 4.41; robust $z = -0.73$; empirical quantile 0.245). This depth fits sequence integrated into host DNA better than free-living contaminant DNA, whose depth would reflect the abundance of the bacterium in the sample. However, the interval lies within the 9.5-Mb terminal block of Chr9 flagged above (Table S9), whose placement the re-assembly does not support, so its chromosomal position is uncertain. In the present assembly, FCS-GX flagged no interval on chromosome 9 (Table S14).

Junction-spanning reads argue against a chimeric join at the interval boundaries. Across the six host–foreign transitions that delimit the flagged region, 18–34 individual HiFi reads align continuously through each junction. These reads span a 2-kb window without clipping within 500 bp of the breakpoint and without indels above 100 bp, and reads carrying supplementary-alignment tags were excluded. This rate is indistinguishable from that at 200 random same-chromosome positions matched for window coverage (median 33, MAD 7; robust $z$ from +0.10 to −1.45), with identical conclusions at MAPQ ≥ 20 and MAPQ ≥ 0. Nucleotide alignment localises the transferred sequence more exactly than the per-sequence FCS action does. Against 60 Mollicutes reference sequences (63.1 Mb; *Spiroplasma*, *Mesoplasma* and *Mycoplasma*, extracted from the GX database itself), only

4,988 bp of the 87.8-kb region carries detectable homology (5.7%, median identity 89.7%), whereas two size-matched flanking host intervals align over 0 bp. All four aligned segments fall within Chr9:6,960,603–6,966,214 (5.6 kb). The largest of them (2,361 bp at 89.7% identity, MAPQ 60) spans the single annotated gene in the region, SQ017506. This gene carries the bacterial-specific domain architecture of a type II topoisomerase (Toprim, HATPase_c, DNA_topoisoIV, DNA_gyraseA_C and DNA_gyraseB_C; Pfam $E = 4.7 \times 10^{-44}$), and its best non-metazoan hit is DNA topoisomerase IV subunit A of *Bacillus subtilis* (bitscore 165.0 against 96.7 for the best metazoan hit). In contrast, three flanking host genes return no non-metazoan hit at all. SQ017506 is also interrupted by a 517-bp intron with canonical GT-AG splice boundaries, which bacterial genes do not possess.

The transferred gene is not detectably transcribed in the venom gland. In two RNA-seq libraries aligned to this assembly (HISAT2; 97.3% and 93.4% overall alignment rate), coverage over both exons, over the intron and over the entire 5.6-kb segment is zero, with no intron-spanning spliced read. In the same libraries, positive-control genes on the same chromosome reach 10.4× and 780.6× (SQ017805) and 6,624.4× (SQ017825). The alignment reproduces the vendor's independent quantification in both directions, including a gene expressed in one library and silent in the other (SQ017825: 0.00× here and FPKM 0.000 in VCK1, a resting venom gland; 6,624× here and FPKM 5,390.8 in V0h1, a venom gland sampled 0 h after venom extraction). The absence of signal at SQ017506 is therefore not a mapping artefact. Only venom-gland libraries were available, so expression in other tissues or developmental stages cannot be excluded. Together, assembly contiguity, host-like depth, a bacteria-specific gene with an acquired eukaryotic intron, 89.7% (not near-identical) sequence identity and the absence of detectable transcription are consistent with an old and apparently inactive horizontal transfer from a mollicute bacterium into the centipede genome, and not with contamination. Two limits apply. First, the donor can be placed only within the Mollicutes and not to species, because the alignments are distributed across six different named genomes. Second, the homology-supported transfer is ~5.6 kb, whereas the remaining 82 kb of the FCS-flagged region carries no detectable bacterial homology at either the nucleotide or the protein level, so the per-sequence FCS assignment over-states the extent of foreign sequence.

At a finer granularity than the official per-sequence criteria, 299 intervals (2.253 Mb, 0.23%) have more than 30% of their alignment coverage assigned to a non-metazoan division, predominantly mycoplasmas (81 intervals), plants (67), alveolates (35) and fungi (80). We report these as an exploratory subset and not as an FCS-GX determination, because the official thresholds (good-pct 30%, good-len 50 kb) retain a long sequence whenever the majority of it is host-derived (Table S8). Two sequences missed by the protein-level screen illustrate its limits. Contig17 fell just below the non-metazoan-fraction threshold (0.423 against > 0.5), and Contig18 was not detected at all, probably because the flagged interval contains little or no predicted coding sequence or lacks informative protein hits.

#### *Comparison made while the re-assembly served as a check*

In this and the following three subsections, "reference" denotes the provider assembly and "re-assembly" the present assembly before chromosome naming and contamination cleaning; toxin identifiers (SQ…) and the count of 180 refer to the provider annotation. The primary re-assembly spans 1.49 Gb in 488 sequences, with 14 chromosome-scale scaffolds. Its consensus quality is higher than that of the reference (Merqury QV 68.8 against 65.9), and Inspector reports 6.9 against 92.6 small-scale errors per Mb. Structural-error counts are of the same order (62 against 48), and gene-set completeness is unchanged (BUSCO 98.6% against 98.7% complete; duplicated 0.6% against 0.5%). Single-assembly k-mer completeness is 92.6% (reference 91.9%), and the union of the two phased haplotypes reaches 99.1%, so the shortfall of a single assembly reflects haplotype-specific k-mers.

The reference (0.98 Gb) is smaller than all k-mer-based estimates of genome size in Note 2 (1.23–1.49 Gb), and the re-assembly lies at their upper end. On our reading, the additional ~0.5 Gb is mainly repeat-rich, and whether it consists of further repeat copies or partly of redundant haplotype remains unresolved. k-mers present in the reads but absent from the assembly fall only from 44.5 to 40.6 million, far fewer than the ~40 million haplotype-specific k-mers expected if the extra sequence were a redundant haplotype at 0.4% heterozygosity, while k-mers at an assembly copy number above four rise from 14.5 to 17.5 million. This sequence also has almost no Hi-C signal and no counterpart with MAPQ ≥ 5 in the reference. This interpretation is an inference, and the additional copies were not validated individually. Like the reference, the re-assembly is not telomere-to-telomere. With one terminal-repeat criterion applied to both assemblies (which differs from the survey in Note 2), 9 ends of the 22 longest primary sequences are supported and no sequence is supported at both ends, compared with 7 ends, all at the 3′ end, in the reference.

#### *Toxin map on the re-assembly*

With the annotation transferred by Liftoff, 179–180 of the 180 toxins map completely to each version of the primary re-assembly, and none is unmapped. In haplotype 1, 177 map completely, one partially and two (SQ010851 and SQ010884) not at all. Both of these map completely to haplotype 2 (179 complete, one partial), so no toxin is missing from both haplotypes. The partially mapped toxin is SQ011341 (coverage 0.46 in the primary and haplotype 1, and 0.25 in haplotype 2). Each version of the primary re-assembly has 174 toxin pairs within 100 kb, as the reference does, and none of them spans a gap or scaffold join; the haplotypes have 177 and 171 such pairs.

#### *Family-size-matched clustering on re-assembly coordinates*

The primary test was repeated on the coordinates of the primary re-assembly (scaffolding with and without contig error correction, the corrected version and two minimum scaffold lengths) and of both haplotypes, under the canonical and unidirectional definitions and with and without the quality filter. The odds ratios were 3.06–4.17 over the 42 configurations with all mapped genes (family-block interval lower bounds at least 1.74) and 3.14–4.72 with strictly mapped genes only (lower bounds at least 1.81), compared with 4.18 (canonical) and 3.98 (unidirectional) on the reference. Without the quality filter, the estimate on re-assembly coordinates differed from that for the same genes on reference coordinates by at most 0.22. With the filter, the genes removed differ between the coordinate systems (by 13–90 genes), and the largest difference, 0.38, therefore mixes coordinate and gene-set effects.

#### *Other observations*

The three chromosome-terminal blocks of the provider assembly (above) are not joined to the corresponding chromosome ends in any version. In the corrected primary re-assembly, the Chr3 and Chr9 blocks each form one separate scaffold and the Chr4 block is split among three. In haplotype 1, all three are dispersed over several sequences. In haplotype 2, the Chr3 block forms one scaffold, and the Chr9 and Chr4 blocks lie mainly (96.5% and 84%) on one scaffold each. The genes of the first 4.4 Mb of reference chromosome 12 lie on unplaced scaffolds in all three re-assembly versions. The haplotypes also indicate a candidate inversion of the terminal ~4.9 Mb of chromosome 12. In haplotype 2, 10 and 12 reads join the two breakpoints in inverted orientation (both segments with MAPQ ≥ 20 and adjacent in read coordinates), while 12 and 20 reads span them continuously. In haplotype 1, however, only 0 and 2 such reads are found, so we do not classify it as a heterozygous inversion. None of these observations bears on the toxin catalogue or on the conclusions of the paper.

#### *Results that changed*

The table lists the principal values from the earlier version (provider assembly and annotation) that changed with the present reference and annotation; it is not exhaustive. The conclusion of the family-size-matched comparison is unchanged. Two defects of the analysis code came to light during the re-run and are reported here. First, the bootstrap analyses of the cross-species comparisons drew all species from a single random-number stream. Changing the centipede input therefore also changed the resampling, and hence the intervals, of species analysed after it (for example, the sea anemone interval of the family–linkage correlation moved from −0.90 to −0.08 to −0.90 to 0.11); point estimates of other species are unaffected. Second, the multiple-testing step of the cross-species within-family test used *P* values hard-coded from an earlier run, and it was recomputed from the current values.

| Analysis | Earlier version (provider assembly) | Present reference |
|---|---|---|
| Assembly | 979.96 Mb, 14 chromosomes; QV 65.9; BUSCO 98.7% | 1,488.0 Mb in 475 sequences, 14 chromosome-scale scaffolds (949.5 Mb); QV 68.8; BUSCO |

| Analysis | Earlier version (provider assembly) | Present reference |
|---|---|---|
| | complete | 98.6% complete |
| Chromosome ends with terminal telomeric arrays | 5 of 28 | 8 of 28 |
| Annotation | 26,089 genes, 30,063 transcripts | 16,703 genes, 20,529 transcripts |
| Curated toxins | 180 (Tier A 77, Tier B 103); 103 in 29 tandem clusters | 188 (Tier A 121, Tier B 67), 28 of them sharing 12 earlier models (22 as split fragments); 118 in 32 clusters |
| Largest toxin cluster (Chr7) | 13 genes in 271 kb | 15 genes in 271 kb |
| Venom-gland DIA proteome | 6,305 protein groups; peptides for 156 toxins, 133 resolved to one locus; protein–RNA abundance ρ = 0.63 | 6,687 protein groups; peptides for 163 toxins, 141 resolved to one locus; ρ = 0.56 |
| Toxin clustering, centipede (family-size-matched odds ratio) | 4.18 (2.60–7.48) | 3.82 (2.57–6.05) |
| Same, with the symmetric quality filter | 4.03 (2.54–7.50), 175 toxins | 3.62 (2.49–5.74), 183 toxins |
| Same, Tier A / Tier B toxins | 5.01 / 4.95 | 4.08 / 3.34 |
| Per-toxin excess clustering ($d(f)$) versus local duplication topology ($\Delta f$), centipede family level (ρ) | 0.66 (0.24 to 0.88); positive | 0.005 (−0.65 to 0.67); no detectable association (conclusion changed) |
| Local duplication topology, centipede toxin minus control (median $\Delta f$) | +0.01 (−0.19 to 0.78) | +0.20 (−0.19 to 0.51); uncertain in both |
| Share of expression from tandem copies, centipede: RNA / protein | +0.25 / +0.23 | −0.01 / +0.10; no longer both positive |
| RNA absolute ΔFC, tandem toxin pairs minus tandem control pairs, 0 | +0.23 (0.04–0.63) / +0.16 (0.03–0.79) | +0.22 (−0.03 to 0.39) / +0.13 (−0.03 to 0.41); intervals |

| Analysis | Earlier version (provider assembly) | Present reference |
|---|---|---|
| h / 1 h | | include 0; with one-to-one expression transfer only, +0.20 (0.02 to 0.42) / +0.17 (0.01 to 0.36) |
| Congener comparison: toxin arrays located in *S. cretica* | 23 of 65 versus 361 of 1,047; +0.04 (−0.12 to 0.21) | 34 of 90 versus 425 of 893; −0.17 (−0.30 to −0.03), Holm *P* 0.033 (conclusion changed) |
| Congener comparison, pre-specified E1 | not evaluable (7 arrays) | −0.77 (−1.51 to 0.05), *P* 0.087 |
| Congener comparison, post hoc homologous-loci discordance | −0.63 (−0.85 to −0.42) | −0.22 (−0.57 to 0.28); no longer supported (conclusion changed) |
| Collinear blocks; toxins in collinear blocks | 139; 1 | 7; 0 |
| Ks, toxin versus non-toxin tandem pairs | 0.069 versus 0.071, *P* 0.74 | 0.062 versus 0.079, *P* 0.0027 (new difference) |
| Penelope elements near toxin clusters | enriched at all three window widths | not enriched |
| Venom-gland co-expression module | *r* = 0.72; 35 toxins | *r* = 0.91; 114 toxins (expression inherited from the earlier annotation; see Note 7) |
| Scoloptoxin orthogroup, gene-wide selection | likelihood ratio ≈ 356; 33 MEME sites | likelihood ratio ≈ 262; 24 MEME sites |
| Coding-sequence pairs with ≥ 98% identity | 4 | 34 |
| Automated ToxProt toxin set, enrichment | 2.0 | 2.41 |

#### *Methods*

Independent assembly re-verification. Because the provider report does not state software versions or command lines, the assembly was re-verified from the primary reads. k-mer spectra were counted with meryl ($k$ = 21) separately for the HiFi reads and for the Illumina survey library and fitted with GenomeScope2 ($p$ = 2). The two libraries gave discordant estimates (HiFi 1.228–1.230 Gb at 0.398–0.446% heterozygosity, model fit 45.9–98.0%; Illumina 1.454–1.487 Gb at 1.33–1.81%, model fit 32.4–96.4%), and both differ from the 976.3 Mb quoted in the provider report. k-mer estimates are therefore reported as supporting information only and were not used to adjudicate assembly content. Haplotypic duplication was assessed with the complete purge_dups workflow including get_seqs (HiFi self-alignment with minimap2 v2.31, -I 8G). The purged assembly was re-scored with the same BUSCO version, lineage and gene predictor as the primary assembly (BUSCO v6.1.0, arthropoda_odb10, miniprot 0.18-r281) and with merqury against the HiFi meryl database, so that the two are directly comparable. Per-sequence depth was obtained by mapping all HiFi reads with minimap2 (preset map-hifi, -I 8G), retaining only primary alignments with MAPQ ≥ 20 (samtools view -F 0x900 -q 20) and computing per-base depth (samtools depth -a -Q 20). Medians and interquartile ranges were summarised per sequence. The overall mapping rate was computed from an unfiltered alignment of the same reads (samtools flagstat; primary alignments only) and is reported separately from the MAPQ ≥ 20 set. The k-mer completeness was taken from merqury (fraction of read solid k-mers present in the assembly), and the purged assembly was scored with the same meryl database and merqury version.

Trans-contact partners were counted from the chromap pairs file. For each region, contacts to each other chromosome were divided by that region's total trans contacts and normalised by the partner chromosome's share of assembly length, and control intervals of equal size were taken from the midpoint of the same chromosome. Each purge_dups-flagged interval was additionally profiled with the same filters (samtools depth -a -b, -Q 20), and its median depth was expressed as a ratio to its own chromosome's median (Table S9). Intervals below 0.6 are reported as candidate anomalous sequence (retained haplotype or collapsed repeat; see above). Overlap between these intervals and the curated toxin genes and tandem clusters was computed with BEDTools intersect.

Contamination was screened by DIAMOND blastp ($E < 1 \times 10^{-5}$) of the predicted proteome against two references assembled for the horizontal-transfer analysis of this study (222,528 metazoan and 375,566 non-metazoan proteins), with protein-to-sequence assignment taken from the GFF3 annotation. A sequence was called suspect only when at least two of three criteria were met: more than half of its proteins had their best hit in the non-metazoan reference (with at least five proteins assigned), median depth was below 0.2× or above 3.0× the chromosomal median, or GC content was below 25% or above 60%. This screen is not a full nt or UniProt search and cannot assign precise taxonomy; it identifies clearly non-metazoan contamination and does not exclude closely related eukaryotic contamination.

A full FCS-GX screen was additionally completed (gx v0.5.5; NCBI GX database build 2023-01-24, 465 GB, all eight files verified against the official md5 checksums). Because the database exceeds the memory of the available compute nodes, it was partitioned into three shards by md5 of the subject sequence identifier (1,009,476, 1,008,909 and 1,007,439 sequences; 3,025,824 in total, matching the source database exactly), each occupying 153–155 GiB. Queries were aligned against each shard in turn (gx align). Per-shard hits were merged with the self record placed first within each query block and passed through the complete official pipeline (gx taxify, classify_taxonomy and action_report) with the asserted division anml:insects, the division under which myriapods are catalogued in the GX database.

Read depth over the flagged interval was computed with samtools depth -a against the HiFi alignment and compared with 200 size-matched control intervals drawn at random from the same chromosome. Junction-spanning reads were counted with pysam at each host–foreign transition, requiring continuous alignment across a 2-kb window, no soft- or hard-clipped alignment terminus within 500 bp of the breakpoint, no deletion, skip or insertion longer than 100 bp inside the window, and no supplementary-alignment (SA) tag. The 200 random same-chromosome controls were filtered to within two-fold of the junctions' window coverage so that mappability was comparable, and the whole analysis was repeated at MAPQ ≥ 0. Homology was assessed by aligning the region and two size-matched flanking host intervals with minimap2 (preset asm20) against 60 Mollicutes sequences retrieved from the GX database across all three shards, and by DIAMOND blastp of the region's predicted protein against the same metazoan and non-metazoan references used above. Transcription was assessed by aligning two venom-gland RNA-seq libraries (VCK1, resting; V0h1, 0 h after venom extraction) to the assembly with

HISAT2 and measuring per-base coverage and intron-spanning spliced reads at the locus. Two independently quantified high-expression genes on the same chromosome were used as positive controls, and control coverage was required to exceed 1× before any absence of signal was interpreted. Because the control distribution is right-skewed by repeat-derived high-depth intervals (median 41.55×, maximum 179.01×), significance was assessed with a robust *z* statistic based on the median absolute deviation and with the empirical quantile, and not with a standard *z* score.

Two limitations apply. First, the equivalence of shard-wise alignment followed by merging to a single whole-database run is not guaranteed by the tool's documentation. A whole-database run was not feasible on the available hardware, so no direct comparison could be made, and the screen should be read as a split-database workflow and not as an official single-pass run. Second, official FCS-GX actions are assigned per sequence under fixed default thresholds (good-pct threshold 30, good-len threshold 50,000), so short foreign intervals inside long, predominantly host-derived sequences do not generate an action. The interval-level tabulation in Table S8 is reported alongside the official actions for this reason.

HiFi read-back at chromosome ends. HiFi reads were mapped to the assembly with minimap2 v2.31 (preset map-hifi) and sorted with SAMtools v1.23.1. For each chromosome end, we counted only primary alignments (MAPQ ≥ 20) whose aligned blocks strictly span the terminal base and measured the outward-facing soft-clip of those reads, and clip sequences were scanned for TTAGG/CCTAA repeats. Depth was computed from CIGAR aligned blocks only, and the per-chromosome baseline is the median over all non-overlapping 10-kb bins after excluding 1 Mb at each end. An end was called as carrying a coherent extension when at least three independent read names contributed soft-clips ≥ 1 kb, and additionally as telomeric when at least two of those clips were ≥ 50% TTAGG/CCTAA. Because the absence of an extension does not establish that an end is complete, ends were reported in four separate categories, together with coverage and evidence-sufficiency flags. Sensitivity was calibrated on controls built from the same reads. Thirty internal positions (15 per orientation) were converted into single-sided synthetic contigs with the source interval and a 60-kb outward buffer masked in the parent chromosome, and 6 telomeric spike-in ends plus 6 length-matched non-telomeric ends were added as semi-synthetic reads. Conditional sensitivity on the pre-screened internal positions was 29 of 30 (15 of 15 at 5′ and 14 of 15 at 3′). The telomeric link was recovered at 6 of 6 spike-in ends, with 0 of 6 false upgrades among the matched non-telomeric controls. This analysis is exploratory, and its negatives are not evidence that an end is complete.

### Note 3. Five-genome comparison: methods, detailed results and sensitivity analyses

#### Methods

##### *Comparative genomes, toxin sets and gene families*

Five genomes were analysed with one framework: *S. mutilans* (this study); *Crotalus adamanteus* GCA_039797435.1; *Stegodyphus dumicola* GCF_044657685.1; *Nematostella vectensis* GCF_932526225.1 (all chromosome-scale); and *Centruroides sculpturatus* GCF_000671375.1 (scaffold-level, scaffold N50 ≈ 0.54 Mb). For *S. mutilans*, the 188 curated toxins were used. For the other four species, toxin genes were proteins with a DIAMOND hit to ToxProt ($E < 1 \times 10^{-5}$, query coverage ≥ 50%, alignment ≥ 40 aa), mapped to genes. These uncurated ToxProt-hit sets are the primary input of all cross-species analyses. The architecture-curated sets (Table S10) and expression-filtered sets (Table S16) were used only in the sensitivity analyses described in this Note (checks run with an earlier implementation). Protein-coding genes with coordinates formed the analysis universe (non-coding-RNA loci excluded).

Three gene-family definitions were used. The primary definition was MMseqs2 (release 18-8cc5c; ref. 34) clustering of canonical-isoform proteomes (one protein per gene) at minimum sequence identity 0.3 with bidirectional coverage 0.5 and cluster reassignment. The second was MMseqs2 clustering with unidirectional coverage 0.5 (cov-mode 1). The third was homology groups formed by single-linkage clustering of all-against-all DIAMOND blastp hits (more-sensitive mode, $E < 1 \times 10^{-5}$) whose alignment covered at least 50% of the longer protein. This criterion was chosen because a laxer rule (30% of the shorter protein) chained 32–41% of all genes into one group; the largest homology group comprised 2.2–7.1% of genes.

##### *Family-size-matched clustering*

A gene in a set was called clustered if another member of the same set lay within 100 kb on the same sequence (interval distance; overlapping genes at distance 0), and all members within the window were searched. The observed statistic was the fraction of toxin genes clustered. The random-gene baseline drew equal numbers of random protein-coding genes (1,000 draws). In each of 1,000 draws, the family-size-matched baseline chose for every toxin family a non-toxin family of the nearest available size that contained at least as many genes as the toxin family contributed toxins. The family was chosen at random among equally near candidates without replacement within the draw (toxin families processed from largest to smallest), and as many genes as the toxin family contributed toxins were sampled from it. Clustering was then evaluated over the union of all sampled genes, so that neighbours from any matched family count, exactly as for toxins. Toxin families for which any draw lacked a candidate would have been removed from both sides; none was. For the centipede, the analysis universe was the 16,476 genes on the 14 chromosome-scale scaffolds. All 188 toxins lie on these scaffolds, so toxins and controls were drawn from chromosomal genes only, and the same gene table was used for the decomposition, cross-family homology, duplication-topology, expression and congener analyses. Effects are odds ratios of the observed fraction against the mean control fraction, with risk differences. One-sided empirical *P* values are $(x + 1)/(B + 1)$.

Uncertainty was estimated by resampling toxin families with replacement (2,000 resamples), carrying with each family its observed clustered count and its mean clustered count across control draws. As a check on spatial dependence, spatially connected components of families (families linked when any of their genes are neighbours) were also resampled. Both intervals are conditional on the observed spatial configuration. When run with the earlier neighbour search and matching order, the implementation reproduces the earlier result on the earlier assembly and annotation (centipede odds ratio 3.98, unidirectional definition). For the centipede, a quality-filtered universe excluded genes overlapping another gene model and genes within 10 kb of an assembly gap, for toxins and controls alike. The gap filter initially failed to match gap and gene coordinates because of a sequence-name mismatch; the corrected filter removes seven genes, none of them toxins (Note 1). The seed was 20260724. Spatial-component intervals are reported from a deterministic re-run that reproduces all point estimates.

The pre-specified sensitivity analyses (Table S24) repeated the test with windows of 25, 50 and 200 kb; separately for centipede Tier A and Tier B toxins, with the control pool excluding the families of all 188 toxin genes; with control families additionally matched on the median pairwise protein identity within the family; and, in the snake, with the authors' annotated venom genes (35) in place of ToxProt homologues. For identity matching, identity was computed identically for toxin and control families from all member pairs or from nearest paralogues, in quartile or quintile strata, and families were measurable when a minimum fraction of their pairs aligned (primary: all pairs, quartiles, fraction 0.5). For the snake, three sets were used: the 126 of 134 genes traceable to an annotated locus, the 78 submitted with venom-specific identifiers, and the 74 of these in the authors' venom-gland expression table. Each result was classed by a pre-specified rule. Robust support required the family-block and spatial-component intervals both to exclude 1 with at least 95% valid resamples. A result was classed as same direction but uncertain when an interval included 1, and as no positive support when the odds ratio was at most 1.

#### *Decomposition of the excess*

For each toxin family *f*, the excess *d*(*f*) is its observed number of clustered toxins minus the mean number of clustered genes among the genes sampled for it across control draws, each evaluated against the full observed or control set. Because every control draw has exactly as many genes as there are toxins, the *d*(*f*) sum to the total excess. We report the largest-family share, the top-3 and top-5 shares of the positive excess, and the positive and negative totals. The largest-family share was compared with the same quantity computed by treating each control draw in turn as pseudo-observed (500 draws). Influence was assessed by removing each of the five largest contributors and recomputing clustering and matching (200 control draws). Each clustered gene was further classified by whether it had a neighbour from its own family, from another family only, or both.

##### *Homology of cross-family neighbours*

For the observed toxin set and 200 control draws, every pair of neighbouring genes from different families was tested for homology with DIAMOND blastp (more-sensitive mode, $E < 1 \times 10^{-3}$, all hits retained). A pair was considered homologous if either direction gave a hit. Each gene was then scored for a same-family neighbour, a homologous neighbour from another family, a neighbour without detectable homology, and only neighbours without detectable homology. Observed fractions were compared with the distribution across control draws (one-sided empirical $P$).

#### Results in detail

##### *Toxin clustering against random and family-size-matched baselines*

Each species' toxin set was compared with two baselines using one statistic, the fraction of toxin genes that have another toxin gene within 100 kb. This fraction counted all such neighbours, including those beyond the adjacent gene in coordinate order (Materials and Methods). The first baseline draws random protein-coding genes. The second baseline draws, for every toxin family, a non-toxin family of the nearest available size and samples from it as many genes as the toxin family contributes. Against random genes, the odds of a toxin gene being clustered were 14–31 times higher in the four bilaterian species (centipede 17.5, snake 18.8, spider 30.8 and scorpion 14.2) and 3.3 times higher in the sea anemone (Fig. 1a). Against size-matched non-toxin families, the odds ratio fell to 3.82 in the centipede (95% family-block CI 2.57–6.05), 6.16 in the snake (3.94–10.42), 3.67 in the spider (1.89–6.54) and 2.79 in the scorpion (2.12–3.92). In the sea anemone, it fell to 1.21 (0.82–1.79; $P = 0.17$). Size matching was exact for 94–100% of family draws under the primary definition (88–100% across the three definitions). Confidence intervals that resample spatially connected groups of families, instead of single families, gave the same conclusions. Among the bilaterian species, the only exception was the snake under the unidirectional definition, whose spatial-component interval included 1 (0.98–3.87; Table S18). The canonical-isoform definition is used as primary here, whereas the analysis plan had named the unidirectional definition as primary (Note 1).

The excess in the four bilaterian species was also found under two other gene-family definitions. These were a coarser MMseqs2 definition with unidirectional coverage, and homology groups built by single-linkage clustering of proteins aligned over at least half of both sequences. The family-block interval of the matched odds ratio excluded 1 in all 12 species-by-definition combinations (Fig. 1b). The odds ratio ranged from 3.60 to 5.81 in the centipede, 2.09 to 6.16 in the snake, 3.38 to 5.16 in the spider and 1.93 to 2.79 in the scorpion. The snake estimate was the most sensitive to the definition, and its magnitude should be read as approximate. When the ToxProt homologues were replaced by the venom genes annotated by the authors of the snake genome (35), the matched odds ratio was 11.0–22.1 across the three author-annotated sets under the primary definition (largest set 22.1, 11.4–44.4) and 2.73–7.90 under the unidirectional definition (largest set 7.90, 2.92–22.1; Table S24). Positive support therefore persisted with the author-annotated set, although the estimate remained sensitive to the family definition and rests on 18–22 families. In the sea anemone, the estimate ranged from 1.21 to 2.14, with intervals that included 1 under two of the three definitions. The evidence is therefore insufficient to establish an excess in that species (Table S24 gives one further window and definition with intervals excluding 1). In the centipede, genes that overlap other gene models or lie within 10 kb of an assembly gap were removed from toxins and controls alike. This left 183 toxins and an odds ratio of 3.62 (2.49–5.74).

The comparator toxin sets are ToxProt homologues, whereas the centipede set was additionally filtered on expression. Further sensitivity analyses of this asymmetry were run with an earlier implementation (implementation A) and are summarised in Note 3; their scope differs from that of the analyses above. When the automated ToxProt definition was applied to the centipede, a smaller excess remained above 1. When the centipede expression filter was imposed on the comparator sets, too few gene families were left for the estimator. The intervals then spanned one to two orders of magnitude and included 1 for the scorpion, so this check is uninformative. Architecture curation, simulated fragmentation and distance-to-end matching addressed definitional and assembly asymmetries against the random-gene baseline. Gene-order and long-scaffold analyses tested the family-size-matched excess in implementation A and gave concordant results. These checks bound some asymmetries, but they do not validate the effect sizes reported here. Thus, the random-gene baseline exaggerates how exceptional toxin clustering is, but the family-size baseline does not account for all of it.

Three further pre-specified analyses addressed the main alternatives to a toxin-associated excess (Table S24). First, the result held across neighbourhood windows. With windows of 25, 50 and 200 kb, the interval of the matched odds ratio excluded 1 in all four bilaterian species under the canonical definition (for example, 3.95 and 3.40 in the centipede and 6.60 and 5.22 in the snake at 50 and 200 kb). The odds ratios were generally lower at 200 kb, as both observed and control fractions rose. Second, toxin families might be younger than other families of the same size. We therefore additionally matched each toxin family to non-toxin families of similar median within-family protein identity, a proxy for duplication history (matched families differed by a median of 2.8–3.2 percentage points). The excess remained in all four species (centipede 3.58, 95% CI 2.29–5.92; snake 9.47, 5.33–17.15; spider 2.76, 1.36–5.23; scorpion 2.39, 1.44–4.26), compared with 3.77, 6.63, 3.68 and 2.24 without this matching on the same measurable families. Similarity is an imperfect proxy for age and may itself absorb signal related to tandem duplication, so this analysis does not exclude confounding by family age. Third, in the centipede the excess was found both for the 121 toxins with a predicted signal peptide (Tier A; 4.08, 2.73–6.40) and for the 67 toxins without one (Tier B; 3.34, 1.80–7.59). The excess is therefore present in both parts of the catalogue, although this does not show that Tier B proteins are secreted. In the sea anemone, the estimates were in the same direction but uncertain in each window under the primary definition.

#### *Distribution and composition of the excess*

Because the excess is a difference between observed and expected clustered genes, it can be divided exactly among toxin families (Materials and Methods). The concentration of the excess varied among species (Fig. 2a). The largest single family carried 7% (centipede), 13% (snake), 32% (spider), 13% (scorpion) and 26% (sea anemone) of the positive excess, and 15, 5, 4 and 9 families were required to account for half of it in the four bilaterian species. None of these shares was clearly larger than expected from the same decomposition applied to control draws (largest-family share, $P$ = 0.088 in the spider and ≥ 0.43 elsewhere; Table S19). When clustering and matching were recomputed after removing each of the five largest contributors in turn, the excess remained in every bilaterian species (odds ratios 2.62–6.12; Table S19).

We next examined what kind of neighbour makes a toxin gene count as clustered. When neighbours were classified by family, toxins had a neighbour from their own family more often than matched controls (centipede 0.47 versus 0.26; snake 0.43 versus 0.16; spider 0.46 versus 0.23; scorpion 0.44 versus 0.28). Toxins also had many more neighbours assigned to other families (Fig. 2b). These other-family neighbours could lack detectable homology, or they could be homologues split across families by the clustering threshold. To distinguish the two, every pair of neighbouring genes from different families was tested for detectable homology (DIAMOND, $E < 10^{-3}$), in toxins and in control draws alike. A homologous neighbour from another family was found for 0.29 of centipede toxins (≤ 0.005 of controls), 0.29 of snake toxins, 0.26 of scorpion toxins and 0.10 of spider toxins. Thus, much of the apparent cross-family clustering is clustering of homologues. Neighbours with no detectable homology were modestly in excess in the centipede (0.19 versus 0.05), snake (0.10 versus 0.04) and scorpion (0.13 versus 0.08), but not in the spider or sea anemone. The fraction of toxins whose only neighbours lacked detectable homology showed no clear excess in any species ($P$ = 0.22–0.48; Table S20). When families were replaced by the coarsest homology groups, a same-group excess was found in all five species (for example, centipede 0.52 versus 0.18; sea anemone 0.34 versus 0.17). Under the family and homology criteria used here, the excess is therefore carried mainly by homologous toxin copies lying next to one another. A smaller, species-dependent excess of neighbours without detectable homology was also observed, but we found no evidence for it independent of homologous adjacency. These patterns do not by themselves identify how the arrays arose.

#### Cross-species comparison: checks run with an earlier implementation

The checks in this Note were run with an earlier implementation (A) of the cross-species comparison. Its own family-size-matched estimates are superseded by the main-text implementation (B; Fig. 1 and Tables S18–S20 and S24) and are not repeated here. This Note reports only checks that implementation B does not cover. The two implementations differ as follows.

| **Element** | **Implementation B (main text; Fig. 1, Tables S18–S20)** | **Implementation A (this Note)** |
|---|---|---|
| Toxin sets | Centipede 188 curated; comparators | Same primary sets; architecture- |

| Element | Implementation B (main text; Fig. 1, Tables S18–S20) | Implementation A (this Note) |
|---|---|---|
| | uncurated ToxProt hits (DIAMOND *E* < $10^{-5}$, query coverage ≥ 50%, alignment ≥ 40 aa) mapped to genes | curated and expression-filtered sets in sensitivity analyses |
| Gene families | Canonical isoform, bidirectional coverage (primary); unidirectional coverage; homology groups | Unidirectional coverage (point estimates); canonical isoform (bootstrap intervals) |
| Neighbour search | All set members within 100 kb | Adjacent gene in coordinate order only |
| Null model | Joint draw; nearest-size family with at least as many members as toxins; no shortfall | Nearest available size; draws truncated when a matched family was smaller |
| Intervals | Family-block and spatial-block bootstrap (2,000) on the same estimator | Family-block bootstrap with 30 control draws for the point estimate |
| Use | All family-size-matched estimates | Only the checks reported in this Note |

Several checks below (architecture curation, simulated fragmentation and distance-to-end matching) were made against the random-gene baseline, and they therefore do not test the robustness of the family-size-matched excess.

Toxin-set quantities for the comparator species:

| Species | ToxProt-hit proteins (input to architecture curation; Table S10) | Proteins passing query coverage ≥ 50% and alignment ≥ 40 aa | Genes after protein-to-gene mapping | Genes with coordinates and a gene family (primary analyses; Table S18) |
|---|---|---|---|---|
| *C. adamanteus* | 463 | 289 | 289 | 289 |
| *S. dumicola* | 693 | 406 | 249 | 249 |

| Species | ToxProt-hit proteins (input to architecture curation; Table S10) | Proteins passing query coverage ≥ 50% and alignment ≥ 40 aa | Genes after protein-to-gene mapping | Genes with coordinates and a gene family (primary analyses; Table S18) |
|---|---|---|---|---|
| *C. sculpturatus* | 740 | 467 | 374 | 374 |
| *N. vectensis* | 745 | 290 | 222 | 222 |

#### Toxin sets and architecture curation

After uniform architecture curation, the comparison sets comprise 459 toxin candidates in the snake, 681 in the spider, 711 in the scorpion and 710 in the sea anemone, compared with the 188 curated toxins of *S. mutilans* (Table S10). The sea anemone set is the one most sensitive to the ToxProt-homology definition used here. Cnidarian venoms are dominated by lineage-restricted families, and our Tier C analysis shows that the curation whitelist does not cover lineage-specific venom domains, so recall in this species is the weakest of the five. This caveat applies directly to the sea anemone results in the main text.

A common analysis framework was applied to five venomous lineages: the centipede (Chilopoda), the snake *Crotalus adamanteus* (Serpentes), the spider *Stegodyphus dumicola* (Araneae), the scorpion *Centruroides sculpturatus* (Scorpiones) and the sea anemone *Nematostella vectensis* (Anthozoa). Four assemblies are chromosome-scale. The scorpion assembly is scaffold-level (scaffold N50 ≈ 0.54 Mb), which we account for below. For *S. mutilans*, the curated 188-gene set was used. For the other four species, toxin genes were called by a uniform automated ToxProt homology search ($E < 1 \times 10^{-5}$; query coverage ≥ 50%, ≥ 40 aa). We then subjected these 2,641 candidate proteins (protein-level counts, before mapping to genes and before the hit-quality filter applied in the primary analyses) to uniform DeepSig- and Pfam-based architecture curation, using the same domain lists and thresholds applied to the centipede set. A total of 2,561 proteins passed a narrow physiological-domain artefact filter (80 removed, 3.0%). They were classified as signal-peptide-supported (Tier A, $n$ = 1,490), toxin-domain-supported

(Tier B, $n$ = 704) or homology-only (Tier C, $n$ = 367) candidates. Tier C denotes candidates lacking signal-peptide support and a match to the centipede-derived toxin-domain whitelist. It should not be interpreted as a false-positive class, because that whitelist does not cover lineage-specific venom domains. Tier C in fact contains recognised venom components absent from the whitelist, including Kunitz/BPTI protease inhibitors ($n$ = 35), L-amino-acid oxidase (Flavin amine oxidoreductase, $n$ = 36), and Pfam model variants of families that are on the list (for example, Reprolysin_5, annotated as metallo-peptidase family M12, and Phospholip_A2_2).

The architecture-curated sets were used only in sensitivity analyses, and the primary analyses use the uncurated ToxProt-hit sets. Unlike the focal centipede toxin set, these sets lack matched venom-gland expression support and should not be interpreted as equivalently validated venom-toxin sets. A homology-only definition is also expected to be incomplete in a specific direction. Highly divergent, silenced or pseudogenised family members that would still be informative about a lineage's venom-gene history fall below the homology threshold and are missed. The definition-sensitivity analyses below (curated versus automated for the centipede; raw versus architecture-curated for the other four species) bound how much this asymmetry affects the comparison, but they cannot recover members that no homology search would retrieve. When the 100-kb clustering test was repeated on the architecture-curated sets, enrichment changed by +2.0% (snake, 10.10 to 10.30), +4.2% (spider, 16.82 to 17.52), +5.4% (scorpion, 8.13 to 8.57) and +9.8% (sea anemone, 1.98 to 2.17). Every test remained significant (one-sided empirical $P$ = 0.001, 1,000 permutations), and the rank order among lineages was preserved. These results indicate that the clustering results are not driven by the physiological-domain artefacts that curation removes. Against random protein-coding genes, the clustering signal in implementation A was stable across a 10–250-kb window grid.

##### *Gene-order control, long scaffolds and the automated definition*

Two checks of the family-size-matched comparison are not covered by implementation B. First, a gene-order-matched control, in which clustering was scored by intervening protein-coding genes instead of base pairs, gave concordant significant enrichment in all four sampled bilaterian species (3.2–5.5; $P$ = 0.002). Second, for the fragmented scorpion assembly, restricting the analysis to long scaffolds (≥ 1 Mb; 49 of 374 toxins) retained both the raw clustering and the family-size-matched excess (gene-order enrichment 3.2; $P$ = 0.004), which argues against a simple assembly-fragmentation explanation. In addition, applying the same automated ToxProt definition to *S. mutilans*, which removes the curated-versus-automated asymmetry, gave a smaller but still significant excess (enrichment 2.4; $P$ = 0.002). This value is within the range of the other bilaterian lineages under the automated definition. The architecture-curated 188-gene set gives a stronger estimate (~3.8), because it removes shared-domain physiological homologues that dilute the toxin-specific signal.

##### *Assembly contiguity and position*

To assess sensitivity to assembly contiguity, we fragmented each of the four chromosome-level assemblies using the empirical scorpion scaffold-length distribution. Across the 80 simulated fragmented assemblies, the realised N50 was 533.8 kb (range 484.8–588.8 kb), closely matching the observed scorpion scaffold N50 of 537.5 kb. Relative to the intact assemblies, species-level median enrichment changed by −0.2% to +1.2% when breakpoint-spanning genes were retained according to their start positions (keep), and by +4.3% to +20.1% when such genes were removed (drop). Changes in individual replicates were more variable, ranging from −18.7% to +23.0% under keep and from −24.1% to +51.8% under drop. Simulated fragmentation did not cause universal convergence towards the scorpion enrichment value of 8.21. Median enrichment after fragmentation remained high for the centipede (17.60 keep, 18.14 drop) and spider (16.85, 20.09), whereas the snake was closer to the scorpion value (10.17, 10.72) and the cnidarian remained below it (2.02, 2.17). Under this simulation model, scorpion-level fragmentation therefore did not eliminate the elevated enrichment seen in the centipede and spider. However, the response was species-dependent and no formal between-species test was performed, so these simulations do not exclude a contribution of assembly fragmentation to individual cross-species differences.

The drop analysis was more variable for the spider (median change +20.1%, largest replicate-specific change +51.8%). It retained 286–314 of the 393 spider toxin genes in the implementation A input set and 13,339–13,490 of 17,182 protein-coding genes. Because both the size and the membership of the retained sets varied among breakpoint realisations, we treat the drop results as a sensitivity analysis supporting qualitative robustness and not as a precise estimate of fragmentation effect size. Finally, after random gene sets were matched by distance-to-sequence-end decile, snake enrichment decreased by 28.9%, whereas the other species changed by between −7.9% and −2.6%. Enrichment remained significant for every species after matching (one-sided empirical $P$ = 0.001). The greater attenuation in the snake is descriptive, because we did not formally test whether its reduction differed from those of the other species.

##### *Within-family comparison*

A family-size-matched control cannot detect clustering that is informative within a family, which is the pattern best documented in snake venom (PLA2 and SVMP arrays). We therefore also compared, inside each gene family that contains both toxin and non-toxin members, the clustered fraction of the toxin members with that of their non-toxin paralogues. No lineage showed a significant excess after correction for testing five species (spider raw $P$ = 0.014, Bonferroni- and Benjamini–Hochberg-adjusted 0.068; all others raw $P$ > 0.12; Table S12), and the spider signal disappeared as the family-size threshold was raised ($P$ = 0.13 at three members per side and 0.25 at four). However, this test is structurally blind to the arrays that motivated it. It requires at least two non-toxin paralogues in the same family, whereas the classic venom arrays are composed almost entirely of toxins. Thus, 91.0% of snake toxin genes (65 of 98 toxin-containing families are all-toxin) never enter the comparison, nor do 67.9%, 57.4%, 56.6% and 46.4% of those in the scorpion, centipede, spider and sea anemone. The absence of a within-family effect should therefore be read as the absence of evidence in the families this design can see, and not as evidence of absence in venom arrays generally.

##### *Applying the centipede expression criterion to the other four species*

Only the centipede set was filtered on venom-gland-enriched expression, and the effect of this asymmetry is quantified here. The data needed to impose the same filter on the comparator sets are available, although the filtered sets cannot carry the primary analysis. Matched venom-gland and control-tissue RNA-seq were obtained for all four species, processed identically to the centipede data, and filtered with the same thresholds. The filter is strongly selective. In the best-matched dataset (*C. adamanteus*, with same-individual controls), it behaves as expected: it retains 74 of 459 candidates (16.1%), and the retention rate tracks the strength of the prior evidence (31.0% for candidates with a signal peptide, compared with 1.4% for those supported only by a toxin domain and 5.1% for homology-only candidates). Retention is 84 of 711 in *C. sculpturatus*, 34 of 681 in *S. dumicola* and 34 of 710 in *N. vectensis*.

The unfiltered ToxProt-hit sets remain the primary input to the family-size-conditioned estimator, because the filtered sets do not support it. The number of gene families available as bootstrap units falls from 89 (centipede) to between 8 and 22 (filtered-set values from the earlier run, not recomputed; Table S16). The resulting 95% intervals span one to two orders of magnitude (*C. adamanteus* 1.38–108.0; *S. dumicola* 2.86–95.0) and include unity for *C. sculpturatus* (0.35–6.53). They are not estimable at all for *N. vectensis*, where every bootstrap replicate is undefined because the filtered set contains no clustered toxin locus; this is a failure of the estimator on that set and not evidence that toxins in this species are unclustered. Two caveats limit how far these filtered numbers can be read. First, control sampling differs among species, from same-individual multi-tissue controls in *C. adamanteus* to cross-project controls in *C. sculpturatus*. The same numerical thresholds therefore do not imply the same evidential quality, and we treat the filter as a comparability check and not as a replacement definition. Second, we do not interpret the differences in retention rate as biological differences, because control matching, tissue proxies and transcriptome sources differ among species (Table S16). The comparison across species therefore rests on the unfiltered sets, and the quantitative effect of the definitional asymmetry is reported here separately from the main estimates.

#### *Methods (implementation A)*

Cross-venom comparison. Five venom genomes (*S. mutilans*; *Crotalus adamanteus* GCA_039797435.1, chromosome-scale; *Stegodyphus dumicola* GCF_044657685.1, chromosome-scale; *Nematostella vectensis* GCF_932526225.1, chromosome-scale; *Centruroides sculpturatus* GCF_000671375.1, scaffold-level, scaffold N50 ≈ 0.54 Mb) were analysed with one framework. Toxin genes were called by DIAMOND searches against ToxProt ($E < 1 \times 10^{-5}$; query coverage ≥ 50%, ≥ 40 aa), and for *S. mutilans* the curated set of 188 was used. Clustering was the fraction of a gene set with another set member within a window (10–250 kb; overlapping-gene distance = 0), compared with 1,000 permutations of equal-size random protein-coding gene sets (non-coding-RNA genes excluded from the eligible universe). Gene families were inferred per genome with

MMseqs2 (release 18-8cc5c; minimum sequence identity 0.3; coverage 0.5, cov-mode 1). As a robustness check, canonical-isoform proteomes were also clustered with bidirectional coverage 0.5 and cluster reassignment. A matched non-toxin control drew, per permutation, non-toxin families of the nearest available size to each toxin family (random choice without replacement) under two estimands. Estimand A compared toxin-labelled genes with an equal number of randomly drawn members of the matched families, and estimand B compared whole toxin families with whole matched non-toxin families. Effects are reported as null-centred odds enrichment with risk ratios and two-sided empirical *P* values. Uncertainty on the family-size-matched excess was obtained by family-level block bootstrap (1,000 resamples of gene families, the non-independent unit; 95% percentile intervals).

Robustness to local gene density was assessed with a gene-order-matched control (clustering scored by intervening protein-coding genes), and the fragmented scorpion assembly was additionally analysed with only scaffolds ≥ 1 Mb. The gene-order-matched control, the long-scaffold restriction and the automated-definition sensitivity analysis each used 500 permutations instead of the 1,000 used elsewhere. Same-family and cross-family neighbours and gene order (0, 1, 5 and 10 intervening genes) were tabulated. To remove the curated-versus-automated definition asymmetry, the same automated ToxProt definition was also applied to *S. mutilans* as a sensitivity analysis.

The four non-focal candidate sets were additionally subjected to uniform architecture curation (Supplementary Code). Signal peptides were predicted with DeepSig (eukaryotic model), and Pfam-A domains with hmmsearch (HMMER 3.4, $E < 1 \times 10^{-5}$ per sequence). The best-scoring domain used for the physiological-domain filter was the domain with the lowest i-Evalue on that sequence, without an additional threshold. The domain set used for Tier B assignment was further restricted to domains with i-Evalue < $1 \times 10^{-5}$, and the same toxin-domain and physiological-domain lists and the same coverage threshold used for the centipede set were applied. A candidate was removed only when its best-scoring Pfam domain was a physiological domain (kinase, Ig, thyroglobulin, ankyrin, methyltransferase or teneurin EGF) and its ToxProt coverage was < 0.30. The centipede tiering additionally accepts high-coverage named-scoloptoxin homology, which has no counterpart in the four non-focal genomes and was therefore not applied. The tiers are evidence levels and not a true/false toxin classification.

The expression criterion applied to the centipede set was also evaluated on all four comparator genomes. Venom-gland RNA-seq (or, for *N. vectensis*, which has no venom gland, cnidocyte-rich tentacle RNA-seq as a proxy toxin-delivery tissue) and control-tissue RNA-seq were assembled for each species and processed with the same pipeline as the centipede data (HISAT2 to the same assembly used here; featureCounts on exon-union length; TPM). The same numerical thresholds were applied (venom-gland/control $\geq 2$ and mean $\geq 5$ TPM). Identical thresholds do not imply identical evidential quality, because control sampling differs among species. The ratio was taken as mean(venom gland)/max(per-control-tissue mean), so that a locus must exceed every control tissue. Applying the criterion reduces the candidate sets from 459 to 74 (*C. adamanteus*), 711 to 84 (*C. sculpturatus*), 681 to 34 (*S. dumicola*) and 710 to 34 (*N. vectensis*), and protein-to-gene mapping was complete (100%) in all four species. The main text uses the sets without the expression filter, for the reason given in the Results, and the architecture-curated sets were used only in sensitivity analyses. Per-species summaries are provided in Table S10, and the availability of the per-protein assignments and Pfam descriptions of Tier C domains is described in the Data Availability section. All permutation analyses in this section used seed 20260724.

Assembly-contiguity and position sensitivity analyses. Scorpion-like fragmentation was simulated by independently sampling lengths with equal probability from the empirical set of scorpion scaffold entries and tiling each target sequence until its full length was covered. The final sampled segment was truncated at the actual sequence boundary before the realised fragment N50 was calculated. Twenty independent fragmentation replicates were generated for each of the four chromosome-level genomes. Two pre-specified treatments of genes spanning simulated breakpoints were evaluated. Under the keep definition, a breakpoint-spanning gene was assigned to the fragment containing its start coordinate. Under the drop definition, it was removed, and for every drop replicate both the observed toxin set and the permutation pool were redefined on the retained fragmented coordinates (the toxin set as the intersection of the original toxin set with the retained genes, and the null pool as all retained protein-coding genes); the resulting sizes were recorded per replicate. Clustering was the fraction of toxin genes having another toxin gene on the same sequence within an interval-to-interval distance of 100 kb. Enrichment was the observed clustering odds divided by the mean clustering odds of 1,000 random protein-coding gene sets of the same size, and one-sided empirical *P* values were $(x + 1)/1{,}001$, where $x$ is the number of permutations with clustering at least as large as observed. For the sequence-end analysis, each gene's distance to the nearer sequence end was divided into deciles, and random gene sets were sampled within these deciles to reproduce the distance-to-end distribution of the toxin genes. This analysis used the intact gene sets and was therefore independent of the keep/drop treatment, which applies only to the fragmentation simulations. The script is included as Supplementary Code, and the availability of the per-replicate results is described in the Data Availability section.

### Note 4. Duplication topology, expression and congener arrays

#### Methods

##### *Local duplication topology*

All toxin families with 4–150 members having coordinates and protein sequences (canonical definition) and an alignment of at least 100 amino acids were analysed. Each was paired with a non-toxin family of the nearest size with 4–150 members, drawn without replacement (seed 20261024). Proteins of all family members were aligned with MAFFT (auto mode), and trees were inferred with IQ-TREE 3 (LG+G4; 1,000 ultrafast bootstraps). Sister pairs were unrooted cherries. The primary statistic used cherries with UFBoot ≥ 95, and cherries with UFBoot ≥ 80 were used in a sensitivity analysis on the same trees with the same permutation and comparison procedure. The statistic was the fraction of such pairs on the same sequence within 100 kb. $\Delta f$ is this fraction minus its mean over 1,000 permutations that reassign the family's members among the family's own coordinates with the tree fixed. Families without a supported cherry, with zero permutation variance or whose tree could not be inferred were not evaluable. Toxin and control $\Delta f$ were compared per species by the difference in medians, with 95% bootstrap intervals (5,000 resamples of families) and rank-sum *P* values, Holm-corrected across species. Species with fewer than eight evaluable families on either side were excluded from testing. The pre-specified bound for a near-zero difference was ±0.10. The family-level association between $d(f)$ per toxin gene and $\Delta f$ was summarised by Spearman's ρ with bootstrap intervals.

#### *Expression analyses*

A gene was defined as tandem if another member of its own family lay within 100 kb. Centipede RNA expression was the mean provider TPM across the nine venom-gland samples (three pooled replicates each at rest and 0 h and 1 h after venom extraction), computed on the earlier annotation and transferred to v3.1 genes as described above. Raw reads for most samples are no longer available, so expression could not be requantified. Copies were treated as distinguishable when at least half of their coding-sequence 31-mers were absent from other members of their family. Values transferred from an earlier model that several v3.1 genes share, or merged from several earlier models, are not locus-specific (31 of the 188 toxins). Against venom-gland TPM requantified on v3.1 from the three available libraries, such values gave a Spearman ρ of 0.65, compared with 0.96 for one-to-one transfers. The RNA endpoints were therefore repeated on the 11,996 genes with one-to-one transfer (Table S22). Centipede protein abundance was the median across the 12 animals of the summed intensity of proteotypic precursors mapping to a single protein. For the other four species, venom-gland TPM (tentacle TPM for the sea anemone) was computed from existing featureCounts tables (exon-union lengths; multi-mapping reads not counted).

Endpoint 1 was the difference between the median $\log_2(x + 1)$ expression of tandem and non-tandem members within each family containing both, where $x$ is TPM or summed precursor intensity; proteins not quantified were treated as missing and not as zero. Endpoint 2 was the share of family expression carried by tandem members minus its expectation when the same number of members is drawn at random (1,000 draws). Toxin families were compared with non-toxin families of the nearest size that also contained both tandem and non-tandem members (difference in medians; 2,000 bootstrap resamples of families; rank-sum $P$ with Holm correction across species within each layer and endpoint). Bounds for near-zero differences were ±0.5 $\log_2$ units and ±0.10 of share. Exploratory coordination measures in the centipede were the absolute difference in $\log_2$ fold change between members of tandem pairs (RNA, 0 h and 1 h versus rest; protein, stimulated versus unstimulated), the sign concordance of these changes, and the correlation of protein residuals after removing treatment-by-sex means, each averaged per family.

#### *Congener comparison*

All 16,703 *S. mutilans* primary proteins (one per gene) were aligned to the *S. cretica* genome (GCA_966189295.1) with miniprot 0.18 (19), keeping up to 20 alignments per protein with a secondary score ≥ 0.5 of the best. Arrays were connected components of same-family genes (canonical definition, all members) within 100 kb, with at least two members; toxin arrays contained at least one toxin. Anchors were up to five nearest protein-coding genes on each side, excluding members of the same family and all toxins. The best alignment of an anchor had to be unique (second-best score < 0.9 of the best), have identity ≥ 0.5 and cover ≥ 70% of the query, and distinct anchors had to map to distinct target loci. An array was located when the target sequence carrying most anchors held at least two anchors on each side, the two sides did not interleave, and the region between the innermost anchors spanned at most ten times the source array or 1 Mb (3 Mb in a sensitivity analysis). Arrays whose regions overlapped were excluded from the primary analysis, as were arrays in families with any protein reaching the 20-alignment limit (no v3.1 protein reached it; at most 15 alignments per protein).

Within a region, alignments of the array's family proteins with identity ≥ 0.4 and query coverage ≥ 0.5 were merged into loci (same strand, coding overlap ≥ 50% of the shorter model, greedy by score, not transitive). Loci better explained by another family's alignment were discarded. Coding-candidate loci were those whose best alignment covered ≥ 80% of the query without frameshift or premature stop. Discordance was $|\log_2((n_{\text{cretica}} + 0.5)/(n_{\text{mutilans}} + 0.5))|$. The pre-specified analysis matched each toxin array with up to three non-toxin arrays of identical member count from families of 0.5–2 times its family size, with each control family used once. Family-averaged differences were compared by family bootstrap (5,000) and sign-flip permutation (10,000), with Holm correction over discordance, location rate and identity. The near-zero bound was 0.3 (about one copy in a three-member array), and results were to be reported descriptively when fewer than eight toxin families were evaluable. This condition applied on the earlier annotation (seven arrays from six families). A post hoc stratified comparison was therefore added against all located non-toxin arrays of the same member count, without the family-size restriction and excluding regions that contain assembly gaps (included in a sensitivity analysis); it is reported here as exploratory.

#### Results in detail

##### *Local duplication topology of toxin and non-toxin families*

If toxin arrays grow by a distinctive mode of local duplication, the closest relatives within a toxin family should lie closer together than the family's spatial layout alone predicts, and more so than in non-toxin families. To test this, we analysed every toxin family with 4–150 members and an alignment of at least 100 amino acids, together with an equally sized non-toxin family. For each family, a maximum-likelihood tree was built and strongly supported pairs of sister copies were taken (unrooted cherries with UFBoot ≥ 95). The fraction of these pairs lying within 100 kb was compared with the same fraction after permuting copies among the family's own positions ($\Delta f$; Materials and Methods). Conditioning on each family's layout in this way separates topology from the clustering already measured in Fig. 1.

In evaluable families, sister copies were closer than their layout predicts in both toxin families (median $\Delta f$ 0.23–0.60 across species, including the snake, whose toxin–control comparison could not be tested) and non-toxin families (0.31–0.49; Fig. 3a). The toxin-minus-control difference was +0.20 in the centipede (95% CI −0.19 to 0.51), +0.19 in the spider (−0.22 to 0.47), +0.14 in the scorpion (−0.21 to 0.46) and +0.23 in the sea anemone (−0.11 to 0.80). None of these differences was detectable after Holm correction ($P \geq 0.41$), and the intervals were too wide to bound the difference near zero (Fig. 3b; Table S21). The snake could not be evaluated. Fourteen of its 28 control families had a strongly supported cherry, but in 13 of them the permutation statistic had no variance, leaving one evaluable control family.

Strong support was itself rarer in toxin trees than in control trees in the centipede, snake and spider (median fraction of unsupported cherries 1.00, 1.00 and 1.00 versus 0.50, 0.93 and 0.33; Fig. 3c). This is expected if toxin arrays contain many near-identical or rapidly diverging copies, and the resulting difference in evaluability is a limitation of this test. In a pre-specified sensitivity analysis on the same trees, relaxing support to UFBoot ≥ 80 made more toxin families evaluable (21, 14, 26 and 10 in the centipede, spider, scorpion and sea anemone, compared with 13, 10, 25 and 9). However, every toxin–control difference remained uncertain (Holm-adjusted $P \geq 0.165$), and the snake remained not evaluable. Across families, the excess per toxin gene ($d(f)$) showed no detectable correlation with $\Delta f$ in any species (centipede Spearman ρ = 0.01, 95% CI −0.65 to 0.67, $n = 13$; snake, spider and scorpion −0.15 to 0.08; sea anemone −0.65, −0.90 to 0.11, $n = 9$). Thus, no relationship between the excess and duplication topology was found. The association between phylogenetic proximity and physical proximity that marks local duplication is present in both toxin and non-toxin tandem families here, and the toxin–control difference was uncertain and could not be bounded near zero. Because gene conversion between neighbours can also make adjacent copies appear as sisters, we describe this pattern as an association between local lineage similarity and spatial proximity.

#### *Expression of tandem toxin copies*

If arrays are retained for their output or for shared regulation, tandem toxin copies should differ from non-tandem copies of the same family in expression, or respond to stimulation together, more than tandem copies of non-toxin families do. We first assessed which copies can be quantified separately. A total of 156 of the 188 centipede toxins passed a within-family 31-mer uniqueness filter, which required at least half of their coding-sequence 31-mers to be absent from other members of their family and was applied identically to control families. This filter does not guarantee unique genome-wide read assignment or correct the provider's quantification of shared reads. In addition, 125 of the 141 locus-resolved toxins were quantified by proteotypic precursors detected in at least six animals. Within families containing both tandem and non-tandem members, the per-copy expression of tandem relative to non-tandem copies did not differ detectably between toxin and size-matched non-toxin families in any species, in RNA or in protein. Nor did the share of family expression carried by tandem copies beyond their number (all Holm-adjusted $P \geq 0.067$; Fig. 4a; Table S22). The centipede share estimates were uncertain (RNA −0.01, 95% CI −0.16 to 0.30; protein +0.10, −0.20 to 0.43), and all other intervals spanned zero widely.

In an exploratory analysis of the centipede venom-gland time course and of the stimulated and unstimulated proteomes, the changes of tandem toxin pairs were not more coherent than those of tandem control pairs. The differences in their RNA responses at 0 h and 1 h after venom extraction (difference in |ΔFC| +0.22, 95% CI −0.03 to 0.39, and +0.13, −0.03 to 0.41) and the protein-level measures were all uncertain (Fig. 4b, c). The analyses were repeated on genes whose expression was transferred one-to-one from an earlier gene model (157 of the 188 toxins). In this subset, the primary RNA endpoints remained uncertain, whereas the differences in RNA responses at 0 h and 1 h became +0.20 (0.02 to 0.42) and +0.17 (0.01 to 0.36), with tandem toxin pairs changing less coherently than tandem control pairs (Table S22). RNA replicates were pooled, protein data came from 12 individuals, and the intervals extend beyond the pre-specified near-zero bounds. These data therefore neither establish nor exclude a link between the excess clustering of toxin genes and the dosage or coordination of the clustered copies.

#### *Toxin arrays in the congener Scolopendra cretica*

To compare the consistency of copy number between toxin and other arrays within the genus, *S. mutilans* arrays (connected components of same-family genes within 100 kb) were located in the chromosome-level genome of *Scolopendra cretica* by flanking anchor genes. Detectable homologous loci between the anchors were then counted (Materials and Methods). Of 983 arrays, 90 contained toxins. Toxin arrays were located less often than non-toxin arrays (34 of 90 versus 425 of 893; family-level difference −0.17, 95% CI −0.30 to −0.03; Holm-adjusted $P$ = 0.033; Fig. 5a). Most failures were due to the lack of two uniquely mapping anchors on each side. Thus, the arrays that can be compared are a selected subset in which toxin arrays are under-represented. Of the 34 located toxin arrays, 23 were neither count lower bounds nor in shared regions; 14 of these (12 families) were evaluable in the pre-specified comparison, and 21 without assembly gaps entered the post hoc comparison (Table S23). In the pre-specified comparison, each toxin array was matched to non-toxin arrays of the same size from families of similar size, and this comparison could be evaluated for 14 toxin arrays from 12 families. The family-averaged difference in the discordance of coding-candidate loci was −0.77 (95% CI −1.51 to 0.05; $P$ = 0.087), or −0.67 (−1.63 to 0.28) when matched regions containing assembly gaps were excluded. The data therefore leave open whether toxin arrays have lower discordance (Table S23).

In a post hoc comparison without the family-size restriction, each located toxin array was compared with all located non-toxin arrays of the same member count, excluding regions that contain assembly gaps (21 toxin arrays from 15 families; 318 control arrays in the matching strata). Discordance in the number of detectable homologous loci (difference in $|\log_2$ ratio$|$ −0.22, 95% CI −0.57 to 0.28) and in coding-candidate loci (−0.18, −0.58 to 0.30) did not differ detectably between toxin and control arrays. Nor did the amino-acid identity of hits to toxin arrays (−0.02; 95% family-bootstrap percentile interval −0.06 to 0.02; sign-flip permutation $P$ = 0.34). Including gap-containing regions gave similar estimates (Fig. 5b, c; Table S23). Homologous loci include fragments and models with coding disruptions. Searches started from *S. mutilans* proteins can miss divergent or *S. cretica*-specific copies, and anchoring excludes arrays that expanded or were rearranged strongly. These results therefore do not show whether toxin copies are retained or replaced, or at what rate, and do not place changes on either lineage.

### Note 5. Divergence times between venom origins

#### TimeTree queries

Pairwise divergence times between the 14 representative species were obtained from the TimeTree public interface (ref. 36; pairwise endpoint; precomputed median age). Species identifiers were taken from the ENA taxonomy service. When a species pair returned no value, the genus and then the family were queried, as registered (Note 1, D6). Values were obtained for 90 of the 91 pairs at species level. The spider–scorpion pair (*Argiope bruennichi*–*Centruroides vittatus*) returned an error at species, genus, family and order level (Araneae–Scorpiones). Three alternative species pairs, registered in a fixed order before querying, also returned no value.

#### Literature estimates for the spider–scorpion split

Published estimates were then used, following criteria registered before the search. Europe PMC was searched with a query combining arachnid taxon names with molecular-dating terms (317 records). Studies were included if they reported an original fossil-calibrated molecular-clock analysis whose sample contained at least one spider and one scorpion, and gave a point estimate for their most recent common ancestor or for crown Arachnopulmonata. Reviews, minimum ages from fossils alone and values cited from other studies were excluded, and one value per study was taken from the main analysis. Values were checked against the full text, and studies without open-access text were obtained by the authors before use.

The 238 open-access full texts and two sets of supplementary files yielded two eligible studies, and five further studies were read in full. Four studies were included: 397 Ma (37), 438 Ma (38), 473 Ma (39) and 509.98 Ma (40). In one study (4), spiders and scorpions did not form a clade, and the value is that of the node joining them, read from the dated tree. In another (37), the calibration intervals were cited in part from an earlier molecular estimate; the registered criteria did not exclude secondary calibrations, and the study was included. The excluded studies, with reasons, are listed in Dataset S4. The median of the four values (455.5 Ma) was used in the primary analysis, and the minimum and maximum in sensitivity analyses.

TimeTree gives 682 Ma for the cnidarian–bilaterian split and 721 Ma for the protostome–deuterostome split. These values come from different sets of studies and are not mutually consistent. The analyses use the TimeTree values as registered, which gives the sea anemone a correlation of 0.054 with each bilaterian; in Fig. 1A the sea anemone is placed at the root for display only.

### Note 6. Order of recruitment and array formation

#### Families and outgroups

The analysis was limited to toxin families with published venom-peptide evidence in the four venomous lineages of our earlier analysis that have suitable outgroups: 51 families in the centipede and 12 in the snake (*C. adamanteus*). The spider and scorpion genomes of our earlier analysis contributed no family, because no published venom proteome was available for these species, and the sea anemone was not paired with an outgroup by design. Outgroups were chosen by the same order as representative species: the millipedes *Archispirostreptus syriacus* (GCA_982267035.2; primary) and *Chamberlinius hualienensis* (GCA_054772095.1; sensitivity) for the centipede, and the non-toxicoferan squamates *Eublepharis macularius* (GCF_028583425.1; primary) and *Podarcis raffonei* (GCF_027172205.1; sensitivity) for the snake.

#### Criteria

For each family, the source array was the connected component (100 kb) containing the most peptide-supported toxins. It was located in the outgroup genome by flanking syntenic anchors (at least two unique anchors on each side, in consistent order). Copies were counted within the anchored interval extended by 100 kb. A family was classified as an ancestral array when the outgroup locus held at least two copies of the family within the window and a strongly supported duplication node (UFBoot ≥ 95) had outgroup copies in both daughter clades. It was classified as expanded after recruitment when the outgroup locus held at most one copy, the venomous species had at least two more copies and twice as many, and the relevant duplication node contained only venomous copies. Gene trees were built with MAFFT L-INS-i and IQ-TREE 3 (ModelFinder; 1,000 ultrafast bootstraps; seed 20260724). Pairs of copies with significant gene conversion (GENECONV 1.81a; global $P < 0.05$) could not support a node. Families with fewer than four sequences, failed trees, too few polymorphic sites for GENECONV or insufficient anchoring were classified as not orderable.

#### Results

With the primary outgroups, 4 of the 12 snake families were orderable (2 ancestral arrays and 2 expansions after recruitment), and none of the 51 centipede families was orderable. In the centipede, 25 families lacked two anchors on at least one side, 14 had no anchor and 12 had tied target sequences; 33 families also had no outgroup homologue in the gene tree. With the sensitivity outgroups, 5 snake families were orderable (4 ancestral arrays and 1 expansion), and no centipede family was orderable. Because 4 of 63 families (6%) could be ordered with the primary outgroups, which is fewer than half and fewer than six, the registered stopping rule applied, and no general conclusion on the order of recruitment is drawn. Family-level results are listed in Table S4.

#### Note 7. Further features of the centipede toxin repertoire

##### Summary

Intragenomic synteny analysis placed 111 of the 188 toxins in tandem gene pairs (MCScanX; a definition based on adjacent homologous genes, distinct from the 100-kb clusters) and none in a retained collinear block. The observable toxin arrangement is therefore dominated by tandem arrays, although an older segmental contribution whose collinearity has eroded cannot be excluded. Transposable-element analyses, orthogroup expansion and selection, a screen for microbial origin and candidate regulators are reported in Notes 7, 7, 7 (Figs. S1, S2 and S6). These analyses are descriptive or correlative. Because duplication boundaries could not be identified, the TE analyses were not used to infer a TE-mediated mechanism.

##### Methods

###### *Clusters, synteny and transposons*

Tandem clusters were defined as venom-gland toxins ≤ 100 kb apart on a chromosome. Family trees were built with MAFFT v7.526 (41) and IQ-TREE v3.1.2 (42) (LG model), and intragenomic synteny was analysed with MCScanX (43). TE co-localisation was analysed with BEDTools v2.31.1 (44) on the RepeatMasker annotation described above (1,233,765 elements after removing simple repeats, low-complexity regions and small-RNA genes, covering 64.0% of the assembly). Gene-level co-localisation was tested against 10,000 control gene sets drawn under a stratified null. Each toxin was matched to non-toxin genes on chromosome and on gene-length quartile, and controls were drawn without replacement within each of the 47 occupied strata. The analysis universe was all 16,703 genes, each of which has a defined nearest-TE distance. Quartile boundaries (1,837, 6,102 and 16,113 bp) were computed across that universe including the 188 toxins; the toxins were then excluded, leaving 16,515 candidate control genes. Significance is the one-sided empirical $P = (b + 1)/(N + 1)$, where $b$ is the number of control sets reaching or exceeding the observed proportion (seed 20260803).

###### *Orthology, gene-family evolution and selection*

Orthogroups were inferred with OrthoFinder v3.1.2 (45) from nine primary-transcript proteomes. Gene-family expansion and contraction were analysed with CAFE 5 (46).

Selection was analysed with HyPhy v2.5.100 using BUSTED (47), MEME (48) and, for the scoloptoxin orthogroup, BUSTED[S] (49).

##### Regulatory analyses

Co-expression modules were identified with WGCNA v1.74 (50) from 18 samples. MicroRNA targets were predicted with miRanda (51). Transcription factors were identified by Pfam DNA-binding domains and filtered for venom-gland enrichment, and de novo promoter motifs were found with STREME (52) (MEME Suite v5.5.9).

#### Toxin clusters, duplication mode and transposable elements

##### Tandem clusters and within-cluster relatedness

The 188 toxins are distributed non-randomly across the genome, with 32 tandem clusters accounting for 118 toxins (Fig. S5). The largest are a 15-gene scoloptoxin cluster on chromosome 7, an eight-gene cluster on chromosome 14 that combines tick-type anticoagulant serpins with scoloptoxin homologues, an eight-gene U-scoloptoxin cluster on chromosome 5 and a six-gene serine-protease cluster on chromosome 8. A maximum-likelihood tree was built for the 15-gene scoloptoxin cluster together with the five non-cluster toxin homologues scoring highest against it by maximum DIAMOND bitscore (22) (MAFFT L-INS-i; IQ-TREE LG+G4 with 1,000 ultrafast bootstrap replicates; support values are ultrafast bootstrap percentages throughout). On this unrooted tree, the 15 cluster genes do not form an exclusive group. Ten fall in one split (bootstrap 89), and the other five group with a single non-cluster homologue, SMUT148260, to the exclusion of the rest (bootstrap 100). No split contains all 15 cluster genes and nothing else. Within this cluster, 44 of the 105 gene pairs were alignable by DIAMOND, with amino-acid identity ranging from 26.9% to 100% (median 66.7%), while the remaining 61 pairs fell below the detection threshold. These observations are consistent with homologous copies occurring both within and outside the cluster, but they do not reconstruct the duplication history. The comparison set was chosen by similarity score and not by phylogenetic criteria, so the tree is not rooted and was not used to polarise duplication events.

To test whether the clusters carry the signature of local duplication, we compared within-cluster sequence relatedness with a permutation null. Within-cluster gene pairs were much more likely to yield a reported DIAMOND alignment than expected after random reassignment of the same 118 clustered genes among groups of the observed sizes (144 of 274, 52.6%, versus 3.6%; 14.74-fold enrichment; permutation $P$ = 0.0010, 1,000 permutations). Because that null permutes genes across chromosomes, we repeated it stratified within chromosomes. The enrichment was smaller but remained significant (52.6% versus 23.0%; 2.28-fold; $P$ = 0.0010). This aggregate enrichment is consistent with a substantial contribution of local duplication to the cluster set. However, it does not establish that every cluster, or every pair within a cluster, is paralogous. All members were assigned to a single MMseqs2 (34) family in 15 of the 32 clusters, and 10 of the 32 clusters had complete significant Pfam coverage with at least one domain shared across all members; Pfam coverage was incomplete for most clustered genes (51 of 118 annotated). Among alignable within-cluster pairs, the median amino-acid identity was 69.9%, which was not significantly higher than the permuted expectation of 59.4% ($P$ = 0.1538). These data therefore do not resolve duplication ages.

#### *Transposable elements at the gene level*

In a genome that is about two-thirds transposable elements, most genes lie near a TE, so we tested TE association at the level of individual genes. Of the 188 toxins, 131 (69.7%) overlap or lie at zero interval distance from a transposable element. Against 10,000 control sets of non-toxin genes matched to the toxins on chromosome and on gene-length quartile and drawn without replacement within stratum, the mean proportion was 74.6% (difference −4.9 percentage points; 0.93-fold; one-sided empirical permutation $P$ = 0.98; 9,815 of 10,000 control sets reached or exceeded the observed value). The corresponding proportion across all 16,703 genes is 74.2%, but this number is descriptive only and is not a valid null. Toxin genes are shorter than genes in that universe (median span 4,583 bp versus 6,102 bp) and are distributed unevenly across chromosomes, whose individual overlap proportions range from 68.3% (Chr5) to 78.0% (Chr12). Individual toxin genes therefore show no detectable excess in TE co-localisation relative to non-toxin genes on the same chromosome and in the same gene-length quartile. This is an absence of evidence for enrichment under the matched null, and it is not evidence that toxins and TEs are unrelated. We describe toxin clusters as TE-associated on the basis of the superfamily-level flanking analysis below. Establishing a causal, TE-mediated duplication mechanism would require duplication-breakpoint, TE-orientation and TE-age evidence beyond the associations reported here.

##### *Transposable elements in cluster flanks*

To examine whether particular TE superfamilies are enriched specifically around toxin clusters, we compared the flanking regions of the 32 tandem clusters (10, 20 and 50 kb on each side) with two controls. The first control was length-matched random intervals drawn on the same chromosome. The second was non-toxin tandem gene clusters (members of one MMseqs2 family, ≤ 100 kb between consecutive members, ≥ 2 members, containing no toxin gene), matched on chromosome and exact member count and drawn without replacement. Effects are reported as the ratio of union base coverage to the permuted mean (1,000 permutations). Each analysis was corrected within its own family of tests by the Benjamini–Hochberg procedure: 33 tests (11 TE classes × 3 widths) for the random-interval control, and 15 tests (5 TE classes × 3 widths) for each of the tandem-cluster control and the single-family sensitivity analysis, which were restricted a priori to the classes of interest. Against random intervals, total TE coverage in cluster flanks was not enriched (0.82–0.89; $q \geq 0.44$), and no superfamily passed correction ($q \geq 0.44$).

For the second comparison, control clusters were matched to each toxin cluster on chromosome and on exact member count, were required to accommodate full 50-kb flanks, and were drawn without replacement. Three toxin clusters had no eligible control: the 15-gene scoloptoxin cluster on chromosome 7, the eight-gene serpin–scoloptoxin cluster on chromosome 14 and the six-gene serine-protease cluster on chromosome 8. On their own chromosomes, no non-toxin cluster combined the same member count with enough room for full 50-kb flanks, so these three were excluded, leaving 29. Across the remaining clusters, Tc1 was enriched at all three widths (1.31–1.45; $q \leq 0.040$) and LTR/Copia at 10 kb (2.70; $q = 0.045$). Total TE coverage did not differ (1.05–1.20; $q \geq 0.063$), Penelope was not enriched (1.01–1.09; $q \geq 0.91$), and unclassified repeats did not differ (0.87–1.01; $q \geq 0.43$).

As a sensitivity analysis, we repeated the comparison on a more closely comparable subset. This subset comprised 27 single-family toxin clusters, defined for this analysis as runs of one MMseqs2 family and therefore not a subset of the 32 toxin clusters (81 of the 188 toxins), compared with single-family non-toxin clusters matched on chromosome and member count. Tc1 remained enriched at all three widths (1.34–1.50; $q \leq 0.039$). Total TE coverage was higher than in the controls at 10 and 20 kb (1.35 and 1.27; $q \leq 0.011$; 1.17 at 50 kb, $q = 0.065$), while Copia (1.63–2.14; $q \geq 0.080$) and Penelope (0.87–1.07) were not enriched. The Tc1 signal therefore recurs in a more closely matched subset and does not depend entirely on the definition used in the primary comparison. However, this subset analysis does not establish that the asymmetry of the primary comparison is irrelevant to the full cluster result. The two analyses cover different toxin sets, and the clusters on each side are built from different starting gene sets (toxin genes only, versus all genes with toxin-containing clusters then removed). In addition, the specific gene family and the cluster span are not matched between groups, and eligibility for the subset requires same-family tandem expansion, which may itself co-vary with TE content. With the earlier assembly and its provider-supplied repeat annotation, Penelope and Copia were also enriched. With the present repeat library applied to the earlier assembly, the Penelope signal disappeared, so it reflected the repeat annotation and not the assembly (Note 2). Neither control matches local gene density, GC content, recombination rate or distance to chromosome ends, and none of these associations is evidence of a TE-mediated mechanism.

#### *Duplication mode*

To characterise the duplication mode, we mapped intragenomic synteny with MCScanX (43) and recovered 7 collinear (segmentally duplicated) blocks and 1,809 tandem gene pairs genome-wide. Of the 188 toxins, 111 fall within tandem arrays, and none is a member of a retained collinear block. The current toxin architecture is therefore dominated by tandem arrays, although an ancient segmental or whole-genome-duplication contribution whose collinearity has since eroded cannot be excluded. Synonymous divergence of toxin tandem pairs (median Ks = 0.062, $n = 48$) did not differ from that of the few collinear pairs (0.070, $n = 23$; permutation $P = 0.51$) but was lower than that of non-toxin tandem pairs (0.079, $n = 480$; $P = 0.0027$). We report this difference descriptively and do not interpret it as dating, because gene conversion between tandem copies can lower Ks.

#### *Further transposable-element analyses*

Each toxin cluster was matched to 20 same-chromosome intervals of identical length by nearest neighbour on local gene density and GC content (31 of 32 clusters matched). Overall TE occupancy inside clusters did not differ from that of matched controls (0.448 versus 0.449; permutation $P = 0.96$), and cluster TEs were not younger than control TEs by RepeatMasker divergence (median 15.4% versus 14.9%; $P = 0.34$). Within toxin clusters, same-family TE copies were more often co-oriented than in matched controls (0.611 versus 0.479; Fisher exact $P = 0.0001$; family-level permutation $P = 0.0092$). The difference persisted when copies less than 20 kb apart or within 2 kb of a toxin gene were removed (Table S13). These orientation measures differ from those of the controls used. However, duplication breakpoints were not identified, and two quantities were not calibrated: the expected co-orientation under local direction bias (which is not 0.5 in general; under independent draws it is $p^2 + (1 - p)^2$ for a local forward fraction $p$) and the dependence among repeats. Co-oriented copies can also be passengers of local duplication as well as substrates for recombination. We therefore report these results as associations with the repeat environment and do not infer a TE-mediated duplication or recombination mechanism. A comparable association between TEs and toxin genes has been reported in a funnel-web spider (53). Intervals between adjacent toxins within clusters show no TE excess over random intervals (occupancy 0.465 versus 0.484), and no recombination map exists for this species.

#### *Methods*

Within-cluster relatedness of tandem toxin clusters. The 188 curated toxin proteins were compared all-against-all with DIAMOND blastp ($E < 1 \times 10^{-3}$, maximum 200 targets), and the highest amino-acid identity per gene pair was retained. For the 32 clusters (118 genes, 274 within-cluster pairs), we scored the fraction of pairs with a reported alignment and the median identity among aligned pairs. Both were compared with 1,000 permutations that reassigned the same 118 clustered genes at random among groups of the observed sizes (seed 20260803). A chromosome-stratified version restricted each permutation to reassignment within the chromosome of origin. One-sided empirical $P$ values were $(x + 1)/1{,}001$. Cluster members were identified by matching SMUT-prefixed numeric gene identifiers. Two auxiliary descriptors were also tabulated per cluster:

whether all members fell in one MMseqs2 family, and whether all members carried a significant Pfam domain (i-Evalue < $1 \times 10^{-5}$) with at least one domain shared. These descriptors are reported as descriptive counts and not as independent tests, and Pfam coverage was incomplete for most clustered genes (67 of 118 lacked any domain at this threshold). For the 15-gene chromosome-7 cluster, the five non-cluster toxin homologues with the highest maximum DIAMOND bitscore against any cluster member were included as comparison sequences (not as phylogenetically justified outgroups). These sequences were aligned with MAFFT L-INS-i and analysed with IQ-TREE (LG+G4, 1,000 ultrafast bootstraps), and the resulting tree is reported unrooted. The availability of the analysis scripts and outputs is described in the Data Availability section.

Transposable-element enrichment in cluster flanks. Flanking regions of the 32 tandem clusters (10, 20 and 50 kb on each side) were compared with two controls. The first control was length-matched random intervals drawn on the same chromosome. Each drawn interval was required to have its own $\pm W$ envelope disjoint from the $\pm W$ envelope of every true cluster, and every permutation was required to yield a complete set of 32 intervals with full-width flanks (seed 20260802). The second control was drawn from 1,068 non-toxin tandem gene clusters, defined as runs of ≥ 2 genes of a single MMseqs2 family with ≤ 100 kb between consecutive members and containing no toxin gene. There were 1,112 such clusters genome-wide, of which 44 contained a toxin gene and were removed; these clusters are defined differently from the same-family arrays of the congener comparison in the main text. Each toxin cluster was matched to one control cluster on chromosome and on exact member count, drawn without replacement, and candidates were further required to accommodate full 50-kb flanks within the chromosome (seed 20260803).

Matching was on chromosome and member count only. Cluster span, gene-family identity, local gene density and GC content were not matched, and control clusters were selected at random among eligible candidates and not by nearest span. Three toxin clusters (the 15-gene scoloptoxin cluster on chromosome 7, the 8-gene serpin–scoloptoxin cluster on chromosome 14 and the 6-gene serine-protease cluster on chromosome 8) had no eligible candidate, because no non-toxin cluster on the same chromosome combined the same member count with room for full 50-kb flanks. They were excluded, leaving 29. A sensitivity analysis (Supplementary Code) restricted both sides to clusters whose members belong to a single gene family, although the starting gene sets remain asymmetric. In this analysis, 29 toxin clusters qualified, of which two were dropped for lack of a candidate, leaving 27 containing 81 of the 188 toxins. The two sides are built from different starting gene sets (toxin clusters from toxin genes only, control clusters from all genes with toxin-containing clusters then removed), and the design does not remove this asymmetry.

TE intervals were merged per superfamily before coverage was computed, so the statistic is union base coverage and not summed annotation length. Effects are the ratio of observed to mean permuted coverage (1,000 permutations). Two-sided empirical *P* values were corrected by the Benjamini–Hochberg procedure within each analysis's own family of tests: 33 tests (11 TE classes × 3 widths) for the random-interval control, and 15 tests (5 TE classes × 3 widths) each for the tandem-cluster control and the single-family sensitivity analysis. The latter two were restricted a priori to the five classes of interest (all TEs combined, DNA/TcMar-Tc1, LINE/Penelope, Unknown and LTR/Copia). Bonferroni-adjusted values are reported for the random-interval analysis only, and the two cluster-based analyses report Benjamini–Hochberg *q* values without a Bonferroni column.

#### Orthogroups, expansion, selection and horizontal transfer

##### *Orthogroups and gene-family evolution*

Nine arthropod proteomes were compared: three centipedes spanning two orders, together with a scorpion, a spider, a tick, a fly, a water flea and a beetle. Of the 188 toxins, 182 were assigned to 72 orthogroups by OrthoFinder (45), and the remaining 6 were singletons left unassigned. In the OrthoFinder assignments of the annotated proteomes, 23 of these orthogroups were represented only in the three sampled centipedes (*S. mutilans*, *S. cingulata* and *Strigamia maritima*). A further 25 were also represented in each of *Drosophila*, *Tribolium* and *Daphnia*; we refer to them as *Drosophila–Tribolium–Daphnia*-shared, a purely operational label, because no ancestral-state reconstruction was performed. The remaining 24 were represented outside the centipedes but not in all three of those proteomes, and 17 of them had a member in at least one sampled chelicerate (Fig. S6). We therefore refer to the first category as annotated-proteome-restricted and not as centipede-specific.

A tblastn re-search of all 48 toxin proteins from these 23 orthogroups against the *Drosophila*, *Tribolium* and *Daphnia* genome assemblies was then performed. It detected a strong genomic-similarity hit for three orthogroups in at least one assembly (union query coverage ≥ 50% and $E < 1 \times 10^{-10}$; two were strong in all three assemblies), weak hits for one (coverage ≥ 30% and $E < 1 \times 10^{-5}$), subthreshold similarity for two, and no detected HSP at the search cutoff ($E \leq 1 \times 10^{-3}$) for 17. Thus, restriction in the annotated-proteome orthogroup assignment is not evidence of genomic absence. Even strong tblastn similarity does not by itself establish an intact gene or orthology, and failure to detect a hit does not establish true absence. The classification is therefore conditional on gene prediction and orthogroup inference.

Gene-family evolution analysis with CAFE 5 was applied to the 72 toxin-containing orthogroups defined above. CAFE requires a family to be inferred present at the root, and 28 orthogroups did not meet that criterion and were not tested, leaving 44. Of these 44 orthogroups, 17 changed significantly (family-wide $P < 0.05$), and 9 of them were expanded on the *S. mutilans* branch. Counting expansions irrespective of significance, 14 of the 44 tested orthogroups were expanded on that branch. The 28 untested orthogroups are not evidence of stasis. When the six orthogroups with the most *S. mutilans* toxin copies were tested with HyPhy BUSTED, all six showed significant episodic positive selection ($P < 0.001$). The scoloptoxin family (the 40 v3.1 counterparts of the 42 members analysed with the earlier annotation) showed strong gene-wide selection (likelihood ratio ≈ 262; 24 MEME (48) sites at $P \leq 0.05$), which was retained under BUSTED[S], a model that accommodates synonymous-rate variation. These results are consistent with expansion accompanied by sequence diversification.

#### *Horizontal transfer*

An Alien Index was computed for all 188 toxins against 375,566 reviewed bacterial and fungal proteins (non-metazoan reference) and nine arthropod proteomes (metazoan reference). It provides no evidence of horizontal acquisition detectable by this method: only 41 toxins have any microbial homologue, and none reaches the threshold for foreign origin (all AI < 0). A maximum-likelihood phylogeny of the most microbially similar family, γ-glutamyl transpeptidase, was rebuilt with the present annotation. It places all nine *S. mutilans* paralogues that pass the alignment-coverage filter on the arthropod side of a bipartition that separates arthropod from bacterial and fungal sequences with UFBoot 100 (Fig. S1).

#### *Methods*

Genome-level re-search of annotated-proteome-restricted orthogroups. All 48 *S. mutilans* toxin proteins belonging to the 23 orthogroups that were represented only in the sampled centipedes were searched against the *Drosophila melanogaster* (GCF_000001215.4), *Tribolium castaneum* (GCF_031307605.1) and *Daphnia pulex* (GCF_021134715.1) genome assemblies with tblastn (BLAST+ 2.17.0; seg filtering on; E-value cutoff 1e-3; up to 500 target sequences). High-scoring segment pairs were grouped into loci by subject sequence, strand and a maximum subject gap of 100 kb. Each new HSP was compared with the largest subject end so far in the group, so that nested HSPs do not split a locus. Query coverage per locus was the union of query intervals, so overlapping HSPs are not counted twice. A locus was called a strong hit at union query coverage ≥ 0.50 with best $E < 1 \times 10^{-10}$, a weak hit at ≥ 0.30 with $E < 1 \times 10^{-5}$, and subthreshold otherwise. Queries returning no HSP at the search cutoff were recorded separately as having no detected hit, and any species that could not be searched would have been recorded as unassessed and not silently omitted. Each orthogroup took the highest-ranking status among its queries and species. The availability of the per-HSP, per-locus, per-query and per-orthogroup tables and of the provenance record (BLAST version, assembly checksums and full commands) is described in the Data Availability section.

#### Candidate regulators of venom-gland expression

Integration of the toxin map with venom-gland RNA-seq and small-RNA data identified a venom-gland-associated co-expression module (WGCNA; $r = 0.91$, $P < 0.001$; 114 of the 188 toxins), 19 venom-gland-enriched transcription-factor candidates and a predicted miRNA–target network (455 miRNAs to 124 toxins) (Fig. S2). The expression values were transferred from the earlier annotation (main-text Materials and Methods). Thus, v3.1 genes that inherit the values of the same earlier gene, such as fragments of an earlier merged model, have identical profiles. This can inflate their co-assignment to one module, and module membership should be read with that in mind. De novo promoter-motif discovery found no motif enriched in toxin promoters at $E < 0.05$ (best $E = 0.48$). These assignments rest on expression correlation and sequence prediction and not on direct regulatory evidence. They are therefore presented as candidates (Fig. S2), and testing them requires chromatin-level data that this study does not provide.

#### Figures

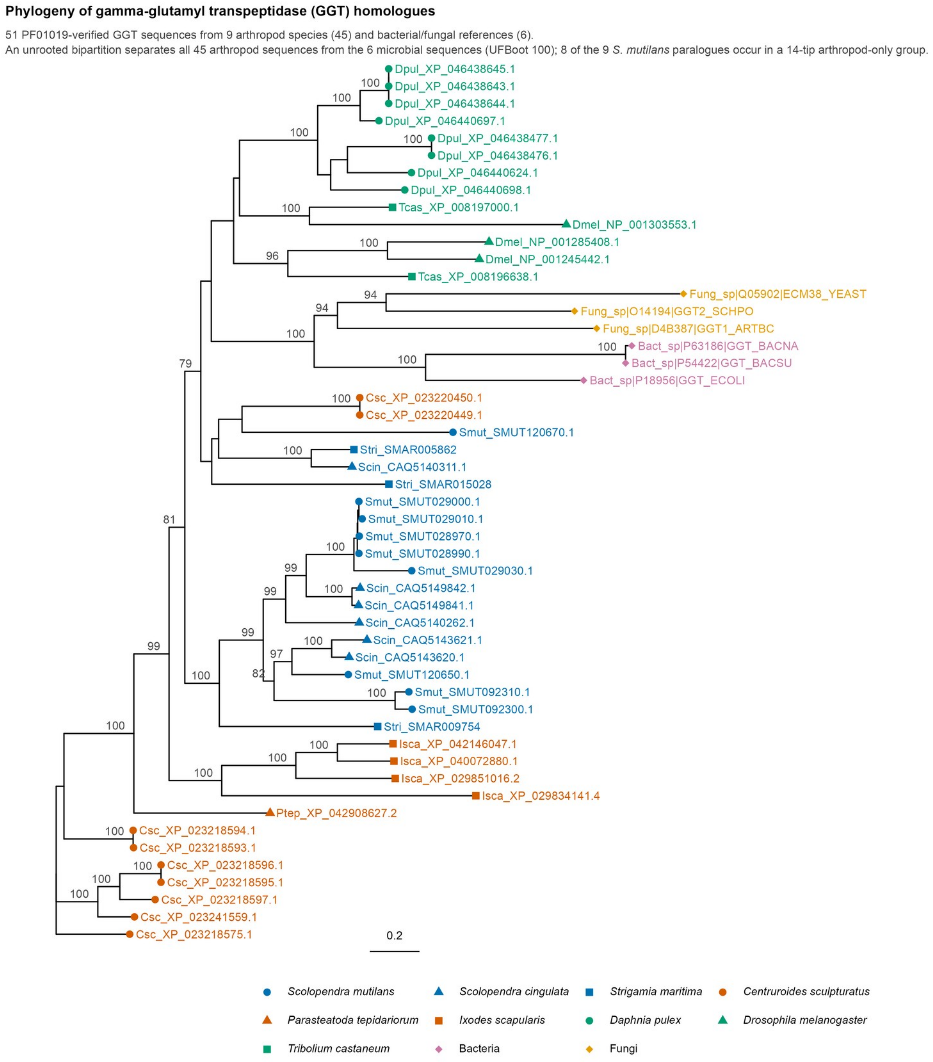


Fig. S1

**Fig. S1.** Maximum-likelihood phylogeny of γ-glutamyl transpeptidase (GGT), the toxin family with the strongest microbial homology. The 11 curated *S. mutilans* GGT toxins (all carrying PF01019) were aligned with 45 PF01019-verified GGT sequences from eight other arthropod species and bacterial and fungal references using MAFFT L-INS-i (single thread); columns with 50% or more gaps and sequences covering less than 50% of the retained columns were removed, leaving 51 sequences from nine arthropod species (45) and microbes (6) and 578 columns (3.7% gaps). The coverage filter removed two partial *S. mutilans* models (SMUT029020, SMUT029040) and three other arthropod sequences. IQ-TREE 3.1.2 LG+G4, ultrafast bootstrap 1000. An unrooted bipartition separates all 45 arthropod sequences from the six microbial ones (UFBoot 100); 8 of the 9 *S. mutilans* paralogues occur in a 14-tip arthropod-only group (UFBoot 100), and the ninth (SMUT120670) also lies on the arthropod side.

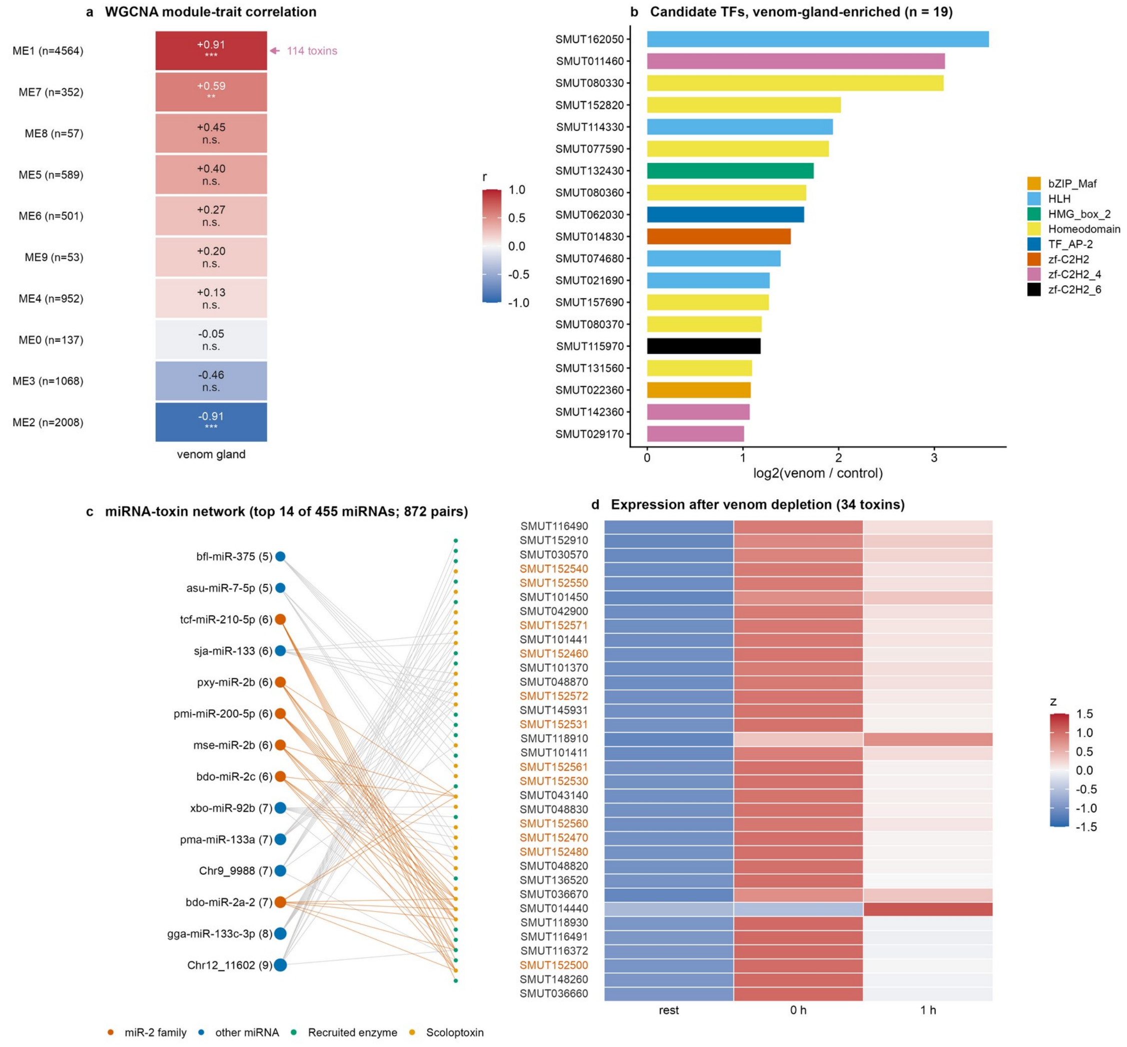


Fig. S2

**Fig. S2.** Candidate regulators associated with venom-gland expression. **a**, Eigengene correlation of the venom-gland-associated WGCNA module. **b**, Venom-gland-enriched transcription-factor candidates by Pfam DNA-binding domain class. **c**, Predicted microRNA–toxin regulatory pairs (top 14 of 455 microRNAs; 872 pairs); labels of the form ChrN_number are identifiers of novel microRNAs predicted from the small-RNA libraries, as assigned in the small-RNA annotation, and are not genomic coordinates. **d**, Expression of the 34 toxins with the largest increase from rest to 1 h after venom extraction (at least 5 TPM at 1 h and more than 1.5-fold the resting value), at rest, immediately after depletion (0 h) and 1 h later, shown as row-scaled *z* values; orange labels mark members of the largest tandem cluster (chromosome 7). Values are transferred from the earlier annotation, so models sharing an earlier source have identical profiles. WGCNA v1.74 on 18 RNA-seq samples; module–trait association by Pearson correlation ($r$ = 0.91, $P$ < 0.001; 114 of the 188 toxins in the module); transcription factors filtered by venom-gland/control expression ratio ≥ 2 and mean ≥ 5 TPM; microRNA targets predicted with miRanda. All associations are correlative and are reported as candidates.

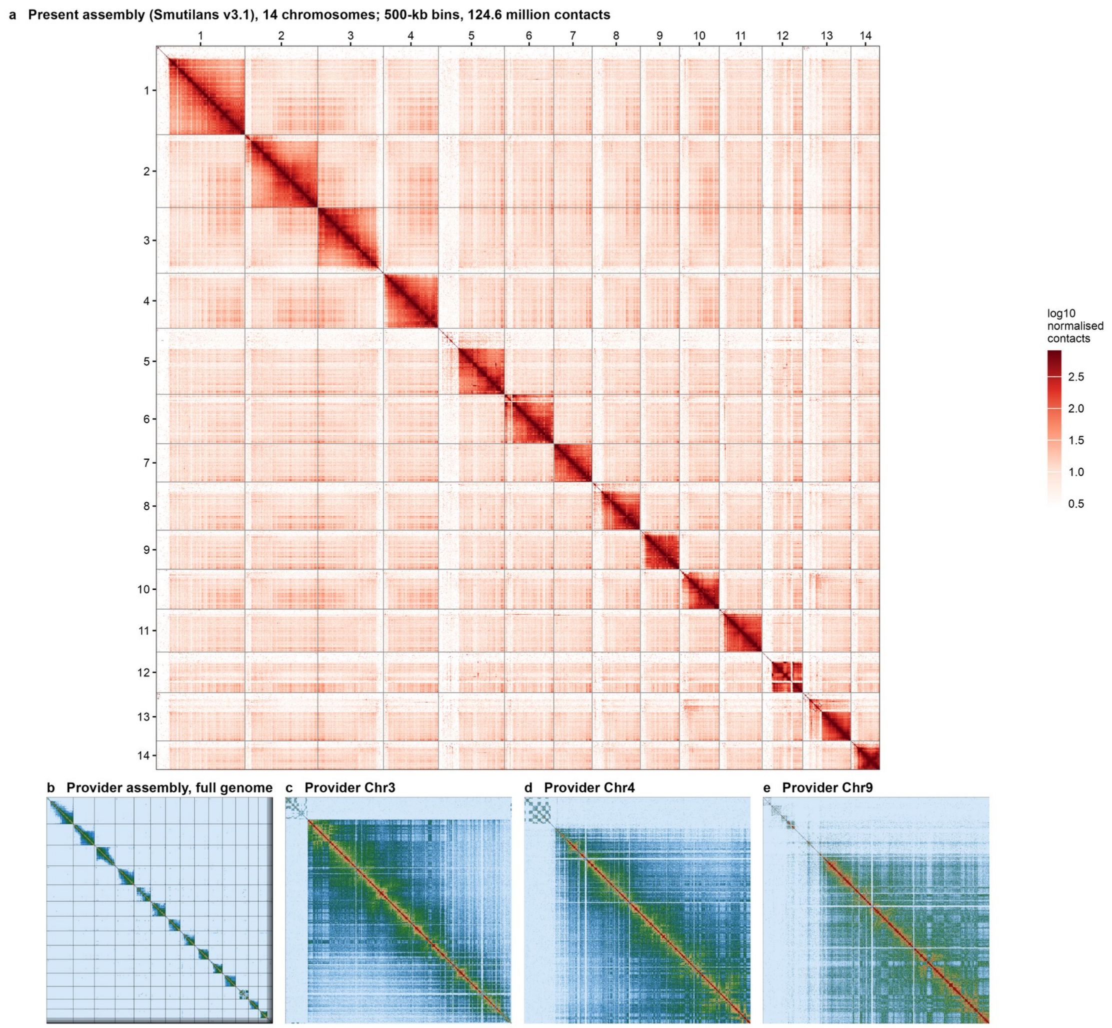


a, YaHS contact pairs (chromap --preset hic) on the present chromosome-scale scaffolds, coverage-normalised; for visual inspection only.
b-e, earlier provider assembly (143.9 million contacts; PretextMap; pale blue low to red high), retained for Supplementary Note 8.

Fig. S3

**Fig. S3.** Hi-C contact maps. **a**, Genome-wide contact map of the 14 chromosome-scale scaffolds of the present assembly (500-kb bins; 124.6 million contact pairs from YaHS scaffolding with chromap preset hic; coverage-normalised, log10 scale; for visual inspection only). Chromosomes are shown in the orientation of the provider assembly, after which they are named. **b–e**, Earlier provider assembly (143.9 million contacts; PretextMap): full-genome map (**b**) and chromosomes 3, 4 and 9 (**c–e**). In each of these three chromosomes the terminal 8.7–9.5 Mb block identified as sub-0.6x depth in Table S9 shows near-absent contact with the chromosome body and an internal checkerboard structure; the dense continuous diagonal of the chromosome body begins distal to the block (Note 2).

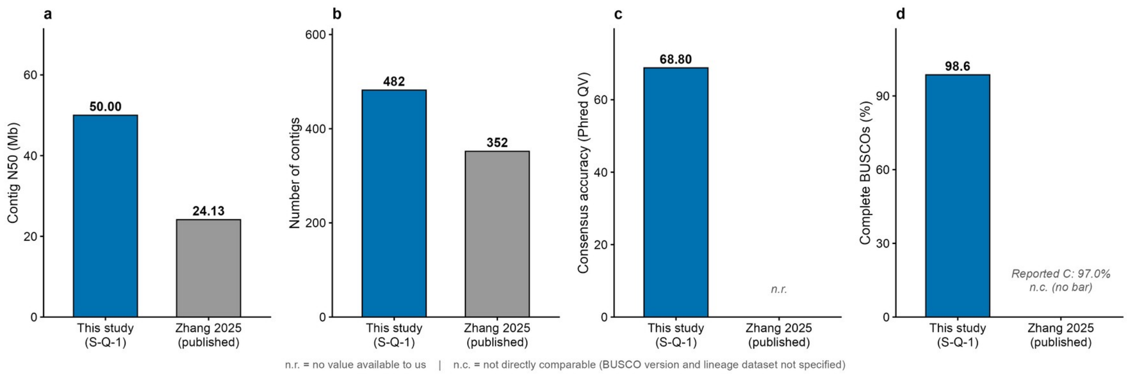


Fig. S4

**Fig. S4.** Assembly metrics of the present genome and the published *S. mutilans* assembly. **a**, Contig N50 (50.00 Mb versus 24.13 Mb). **b**, Number of contigs (482 versus 352). **c**, Consensus quality value of the present assembly (merqury QV 68.80; measured, like **d**, before the final one-contig correction and contamination cleaning). **d**, Genome-mode BUSCO completeness of the present assembly (98.6% complete BUSCOs; arthropoda_odb10, *n* = 1,013). For the published assembly no consensus quality value was available to us (*n.r.*), and its reported BUSCO completeness (97.0%) used an unspecified BUSCO version and lineage dataset and is not directly comparable (*n.c.*; no bar drawn). *No statistical test is applied; values are point estimates from this study's outputs or from Zhang et al. (2025).*

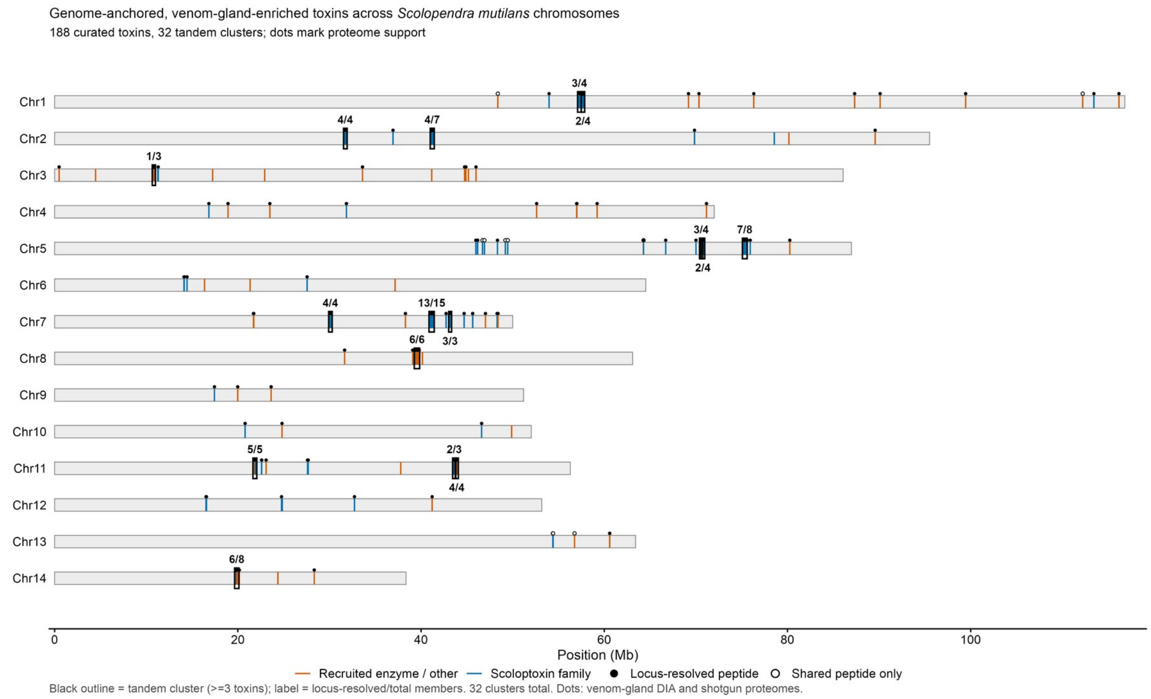


Fig. S5

**Fig. S5.** Chromosomal distribution of curated venom-toxin loci. Positions of the 188 curated venom-gland-enriched toxin genes across the 14 chromosomes. Of the 32 tandem clusters (toxins ≤ 100 kb apart on the same chromosome), the 16 with three or more members are boxed and labelled with the number of locus-resolved members over the total; the 16 two-member clusters are not boxed. Dots above each locus mark proteome support from the venom-gland DIA proteome and the earlier shotgun data combined: filled, at least one genome-unique (locus-resolved) peptide; open, only peptides shared with other genes. Chromosome lengths are drawn to scale. Per-gene support is listed in Table S17 (Table S7 for the published venom sample alone).

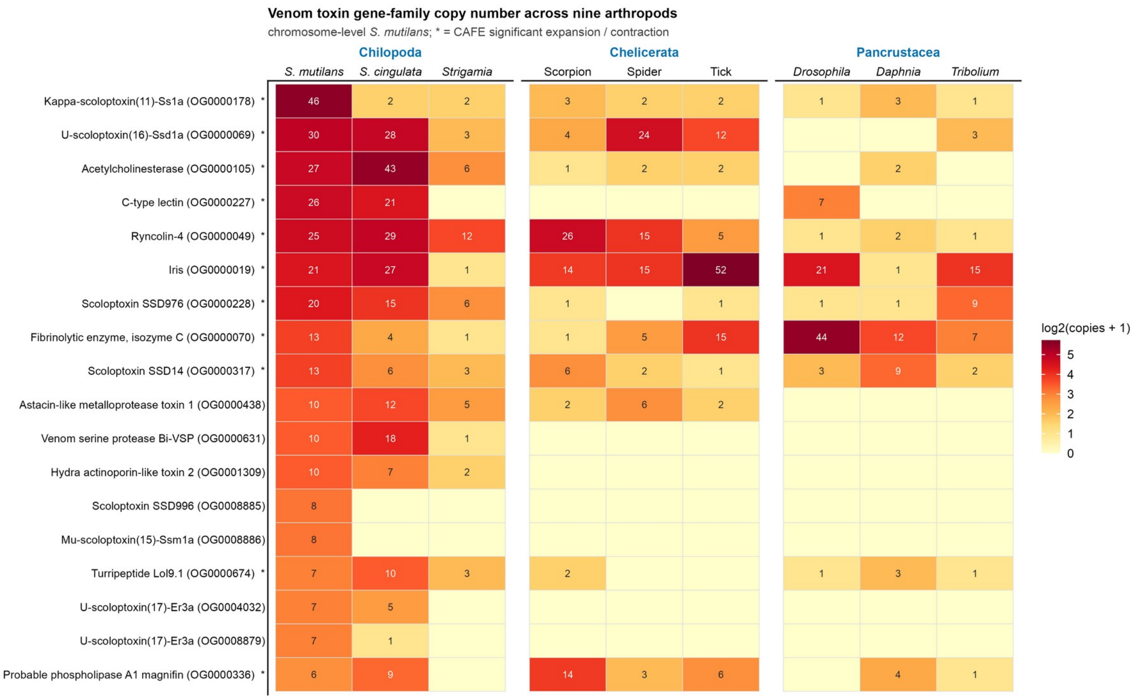


Fig. S6

**Fig. S6.** Copy number of toxin orthogroups across nine arthropod proteomes. Copy number per orthogroup for the largest toxin orthogroups in *S. mutilans* and eight other arthropods. Orthogroups were inferred with OrthoFinder v3.1.2 from primary-transcript proteomes; gene-family expansion and contraction tested with CAFE 5 across the species tree; asterisks mark orthogroups with a family-wide $P < 0.05$. $n = 9$ proteomes; 72 toxin-containing orthogroups in total. Zero cells denote absence from the annotated-proteome orthogroup assignment and do not establish absence from the corresponding genome (Note 7).

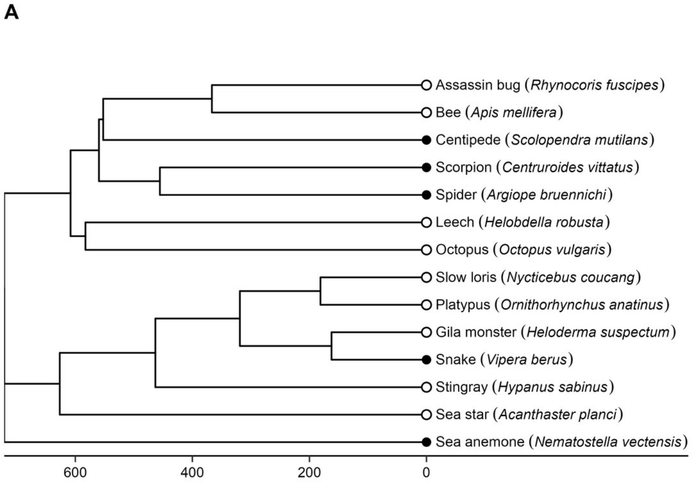


**B**

| Assembly | Strict (P) | Intermediate (P+E) | Broad (P+E+H) |
|---|---|---|---|
| Chr | 0 | 0 | 144 |
| Chr | 79 | 79 | 142 |
| Chr | 143 | 188 | 188 |
| Scaf | 0 | 0 | 259 |
| Chr | 0 | 0 | 309 |
| Scaf | 0 | 0 | 125 |
| Chr | 23 | 23 | 110 |
| Chr | 0 | 0 | 243 |
| Chr | 0 | 12 | 237 |
| Chr | 0 | 0 | 343 |
| Chr | 23 | 23 | 243 |
| Chr | 0 | 0 | 231 |
| Chr | 0 | 0 | 249 |
| Chr | 17 | 17 | 235 |

○ Added origin ● Lineage in earlier analysis Estimable (≥10 families with ≥2 toxins) Not estimable

Fig. S7

**Fig. S7.** Venom origins, representative species and toxin evidence tiers. (*A*) Time tree of the 14 independent venom origins, one representative species each, built from TimeTree median pairwise divergence times (million years ago). The spider–scorpion time is the median of four published estimates (455.5 Mya). TimeTree gives a younger time for the cnidarian–bilaterian split (682 Mya) than for the protostome–deuterostome split (721 Mya), and for display only the sea anemone was placed at the root; the correlation matrix used in the analyses retains the TimeTree values. (*B*) Assembly level (Chr, chromosome scale; Scaf, scaffold level) and number of toxin-associated genes in the strict (P, gene-unique venom peptide), intermediate (P or venom-gland expression, E) and broad (P, E or ToxProt homology, H) tiers. Blue cells are estimable tiers, with at least ten gene families containing two or more toxin-associated genes.

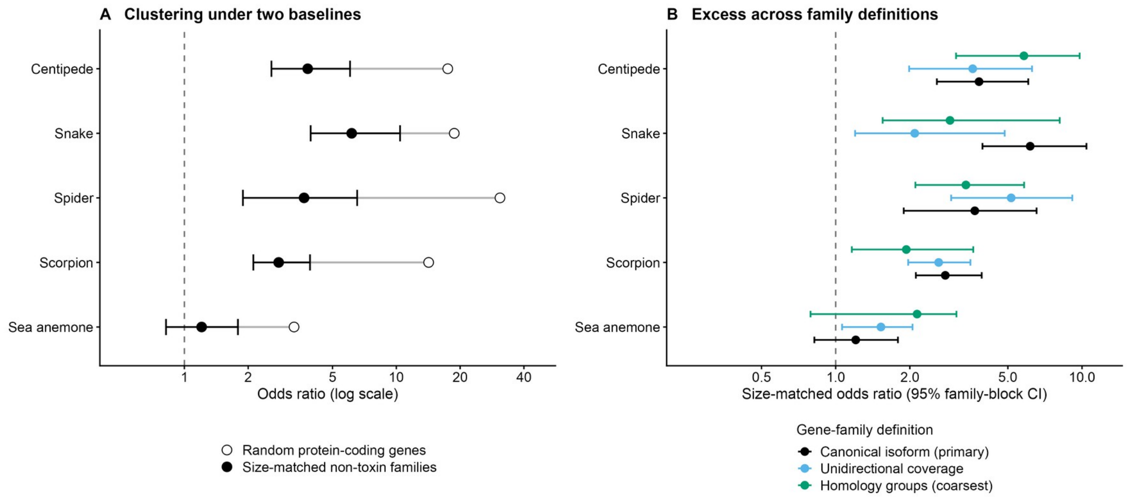


Fig. S8

**Fig. S8.** Clustering of toxin genes against random and family-size-matched baselines in five venomous genomes (the five-genome analysis). (*A*) Odds ratio for a toxin gene having another toxin gene within 100 kb, relative to random protein-coding genes (open circles; point estimates) and to genes drawn from non-toxin families of the nearest size (filled circles), under the primary gene-family definition. (*B*) Family-size-matched odds ratio under three gene-family definitions (canonical isoform, the primary definition; unidirectional coverage; homology groups). Each baseline used 1,000 draws; bars, 95% family-block bootstrap intervals (2,000 resamples of toxin families). One-sided empirical $P \leq 0.001$ for all bilaterian estimates; for the sea anemone, $P = 0.17$ under the primary definition. Numbers of toxin genes and families per species are given in SI Appendix, Table S18.

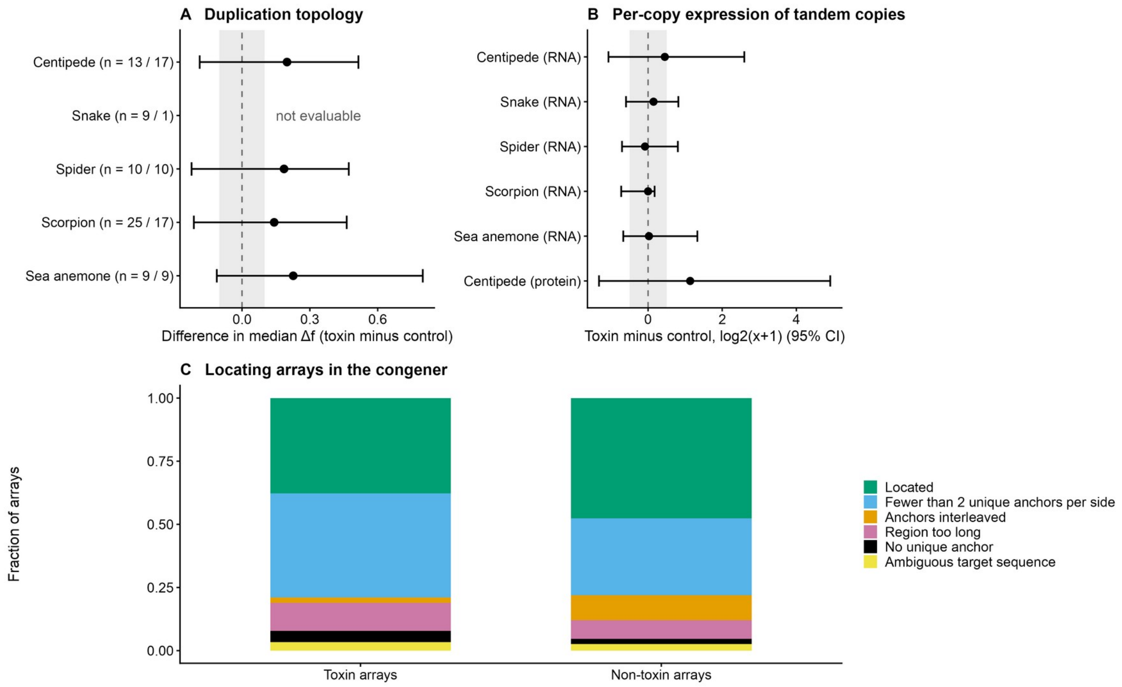


Fig. S9

**Fig. S9.** Duplication topology, expression and congener arrays of toxin and non-toxin families. (*A*) Difference between toxin and size-matched non-toxin families in the median Δ*f*, the fraction of strongly supported sister pairs (UFBoot ≥ 95) lying within 100 kb minus its expectation when members are permuted among the family's own positions; bars, 95% bootstrap intervals (5,000 resamples of families); shading, pre-specified near-zero bound (±0.10). Labels give the numbers of evaluable toxin and control families (n = toxin / control). The snake was not evaluable. Holm-adjusted $P \geq 0.41$. (*B*) Difference between toxin and control families in the per-copy expression of tandem relative to non-tandem members (venom-gland RNA; tentacle RNA for the sea anemone; centipede venom-gland proteome); bars, 95% bootstrap intervals (2,000 resamples of families); shading, pre-specified near-zero bound (±0.5). Holm-adjusted $P \geq 0.067$. (*C*) Outcome of locating *S. mutilans* same-family arrays in the congener *Scolopendra cretica* by flanking anchor genes (34 of 90 toxin arrays and 425 of 893 non-toxin arrays located). Sample sizes for all panels are given in SI Appendix, Tables S21–S23.

#### Legends for Tables S1 to S26

**Table S1.** Candidate venom origins, eligibility and representative species.

**Table S2.** Size-matched clustering in 14 venom origins and pooled estimates.

**Table S3.** Clustering of toxin-associated genes relative to secretory and defence (S) and chemoreceptor (L) control families.

**Table S4.** Order of recruitment and array formation: family-level classification.

**Table S5.** Per-chromosome-end telomere status of the present assembly (Smutilans v3.1) and of the earlier provider assembly.

**Table S6.** Toxin catalogue of the present annotation (Smutilans v3.1).

**Table S7.** Anchoring of the published venom proteome to the 188 curated toxins (Smutilans v3.1): the 51 peptide-compatible toxins are listed (the remaining 137 had no compatible peptide in this proteome); 34 are locus-resolved.

**Table S8.** Per-sequence contamination and depth screen of the earlier provider assembly of S. mutilans. Earlier provider assembly (Note 2): coordinates, chromosome names and the 180-gene toxin set refer to that assembly, not to Smutilans v3.1.

**Table S9.** Interval-level read depth of all 114 purge_dups-flagged intervals of the earlier provider assembly. Earlier provider assembly (Note 2): coordinates, chromosome names and the 180-gene toxin set refer to that assembly, not to Smutilans v3.1.

**Table S10.** Per-species toxin candidate sets after uniform architecture curation (b2_curate.py; Supplementary Code).

**Table S11.** Trans-contact partners of the three terminal blocks of the earlier provider assembly and of matched control intervals. Earlier provider assembly (Note 2): coordinates, chromosome names and the 180-gene toxin set refer to that assembly, not to Smutilans v3.1.

**Table S12.** Within-family control: clustering of toxin members versus their non-toxin paralogues.

**Table S13.** Repeat-environment measures in toxin clusters versus matched intervals (descriptive; Smutilans v3.1).

**Table S14.** FCS-GX contamination screen of the S. mutilans v3.1 assembly and verification of retained intervals.

**Table S15.** Gene models of the present annotation (Smutilans v3.1) that overlap the ten retained FCS-GX FIX intervals (Spiroplasma-assigned; Note 2, Table S14).

**Table S16.** Applying the centipede venom-gland expression criterion to the other four species.

**Table S17..**

**Table S18.** Toxin-gene clustering against random protein-coding genes and against size-matched non-toxin families, by gene-family definition.

**Table S19.** Decomposition of the family-size-matched excess among toxin families (canonical definition).

**Table S20.** Neighbour composition of toxin genes and of genes sampled from size-matched non-toxin families (200 control draws).

**Table S21.** Local duplication topology per family (toxin families with 4-150 members and size-matched non-toxin families).

**Table S22.** Expression of tandem copies: toxin versus size-matched non-toxin families.

**Table S23.** S. mutilans same-family arrays located in the S. cretica genome (GCA_966189295.1).

**Table S24.** Pre-specified sensitivity analyses of the family-size-matched clustering test (Q1).

**Table S25.** Assembly metrics of the present assembly (Smutilans v3.1), its phased haplotypes and HiFi-only contigs, and the earlier provider assembly.

**Table S26.** Non-venomous controls (post hoc; SI Appendix, Note 1, D13): ToxProt homologues analysed with the same procedure as the broad tier of the venom panel.

#### Legends for Datasets S1 to S4

**Dataset S1.** Frozen analysis plan (10 November 2026): original Chinese text (authoritative) and English translation.

**Dataset S2.** Dated deviation log (entries D1–D14): original Chinese text (authoritative) and English translation.

**Dataset S3.** Toxin-associated genes and evidence tiers for the 14 representative species.

**Dataset S4.** Literature screen for spider–scorpion divergence estimates: records, decisions and reasons.

#### Public genome assemblies analysed

*Crotalus adamanteus*, GCA_039797435.1, comparator genome of the five-genome analysis (Note 3) (54)

*Rhynocoris fuscipes*, GCA_040020575.1, representative species (Table S1) (55)

*Chamberlinius hualienensis*, GCA_054772095.1, outgroup for the order of recruitment (Note 6) (56)

*Octopus vulgaris*, GCA_951406725.2, representative species (Table S1) (57)

*Scolopendra cretica*, GCA_966189295.1, congener (Note 4) (58)

*Archispirostreptus syriacus*, GCA_982267035.2, non-venomous control (Table S26) (59)

*Drosophila melanogaster*, GCF_000001215.4, comparative genomics (Note 7) (60)

*Helobdella robusta*, GCF_000326865.2, representative species (Table S1) (61)

*Centruroides sculpturatus*, GCF_000671375.1, comparator genome of the five-genome analysis (Note 3) (62)

*Apis mellifera*, GCF_003254395.2, representative species (Table S1) (63)

*Ornithorhynchus anatinus*, GCF_004115215.2, representative species (Table S1) (64)

*Daphnia pulex*, GCF_021134715.1, comparative genomics (Note 7) (65)

*Podarcis raffonei*, GCF_027172205.1, outgroup for the order of recruitment (Note 6) (66)

*Nycticebus coucang*, GCF_027406575.1, representative species (Table S1) (67)

*Eublepharis macularius*, GCF_028583425.1, non-venomous control (Table S26) (68)

*Hypanus sabinus*, GCF_030144855.1, representative species (Table S1) (69)

*Centruroides vittatus*, GCF_030686945.1, representative species (Table S1) (70)

*Tribolium castaneum*, GCF_031307605.1, comparative genomics (Note 7) (71)

*Stegodyphus dumicola*, GCF_044657685.1, comparator genome of the five-genome analysis (Note 3) (72)

*Acanthaster planci*, GCF_054643075.1, representative species (Table S1) (73)

*Heloderma suspectum*, GCF_057233095.1, representative species (Table S1) (74)

*Nematostella vectensis*, GCF_932526225.1, comparator genome of the five-genome analysis (Note 3) (75)

*Argiope bruennichi*, GCF_947563725.1, representative species (Table S1) (76)

*Vipera berus*, GCF_964194415.1, representative species (Table S1) (77)

*Homo sapiens,* GCF_000001405.40, non-venomous control (Table S26) (78)

*Hemiscyllium ocellatum*, GCF_020745735.1, non-venomous control (Table S26) (79)

*Drosophila mauritiana*, GCF_004382145.1, non-venomous control (Table S26) (80)

*Crassostrea virginica*, GCF_053477285.1, non-venomous control (Table S26) (81)

*Asterias amurensis*, GCF_032118995.1, non-venomous control (Table S26) (82)

*Leiobunum rupestre*, GCF_980474605.1, non-venomous control (Table S26) (83)
